# SeCAT (Sequence Consensus Amplicon Trimming): a simulation-informed Nextflow pipeline for cross-study harmonisation of 16S rRNA metabarcoding datasets

**DOI:** 10.64898/2026.11.03.763908

**Authors:** David Terrey, Paula Huber, Emiliano Pereira, Maria C. Pelayo, Robert M. W. Ferguson, Alex J. Dumbrell, Michael Sweet, Alessia Bani, Sebastian Metz

## Abstract

Global syntheses of 16S rRNA metabarcoding data rely on combining datasets, yet amplicon sequence variants generated with different primer pairs can rarely be compared directly. Existing approaches correct batch effects statistically or aggregate taxonomically; none first establishes whether the sequences are positionally comparable. We present SeCAT (Sequence Consensus Amplicon Trimming), a Nextflow pipeline that maps every study’s variants onto a 16S rRNA full-length reference alignment, determines the largest window shared across studies, and tests by simulation whether trimming to that window degrades community structure before any data are merged. Applied to thirteen marine studies sequenced with three primer sets, twelve converged on a consensus window with a median trimmed read length of 249 bp (9,582 alignment columns). Degradation tests across twelve studies and six taxonomic ranks (72 study-rank combinations) returned 31 passes, 38 cautions and three confirmed failures. Eleven studies (682 samples after filtering) were harmonised (trimmed to the window and merged), collapsing 20,891 primer-specific variants into 12,667 MetaASVs, trimmed sequences deduplicated across studies. Harmonisation preserved community structure at family level (Mantel r = 1.00; Procrustes mZ < 0.001) without adding between-study variance (|ΔRZ| < 0.001). Alpha diversity, including observed richness, was recovered across all levels (intraclass correlations 0.93–1.00), whereas sequence-level congruence was moderate (Procrustes mZ = 0.52), as expected where between-study similarity at sequence level exists only after harmonisation. The harmonised output supports synthesis from aggregated ranks down to exact sequences. By establishing positional comparability before merging, SeCAT opens the archive of primer-fragmented 16S rRNA surveys to sequence-level synthesis.

## Introduction

Microbial communities underpin ecosystem function in every habitat on Earth and 16S rRNA metabarcoding is one of the principal means of characterising the taxonomic composition and diversity of their bacterial and archaeal communities, from single habitats to global syntheses (Tremblay *et al*. 2015). Two decades of high-throughput sequencing have filled public archives with amplicon datasets spanning many hosts, habitats and scales (Gilbert, Jansson & Knight, 2014; Thompson et al. 2017). Pooling these data offers statistical power and geographic reach (Thompson et al. 2017; Ma et al. 2022). Cross-study meta-analyses already exploit this, recovering consistent shared and condition-specific signals across independent 16S datasets (Duvallet *et al*. 2017; Ramirez et al. 2018; Fediajevaite et al. 2021). Whole fields are now synthesising their accumulated surveys into single, coherent pictures (Vadillo Gonzalez et al. 2026). The drive for coherent synthesis is not strictly academic: microbial life remains largely absent from global conservation frameworks, initiatives such as the newly launched IUCN Microbial Conservation Specialist Group aim to redress this (Gilbert et al. 2025), and the assessments they require will rest on evidence synthesised across many independent surveys. Harmonisation of accumulated data is therefore a priority for both policy and research.

Primers for 16S rRNA metabarcoding are designed against the conserved regions that flank the gene’s nine variable regions, V1 to V9 (Van de Peer, Chapelle & De Wachter, 1996; Janda & Abbott, 2007), so the choice of primer pair determines which segment of the gene, and which window of its information, a study captures (Klindworth et al. 2013; Bukin et al. 2019).

This approach, however, is not neutral: differences between the conserved regions, and therefore in primer annealing, result in unequal amplification of the bacteria and archaea present in samples (Abellan-Schneyder et al. 2021; Martinez-Porchas et al. 2017). Sequencing of the same samples using varying primers returns divergent downstream community estimates depending on the target region and no single primer set recovers the full diversity (Bukin *et al*. 2019; Tremblay *et al*. 2015; Lee *et al*. 2023). As such, hypervariable regions differ markedly in resolving power with no region considered optimal across different study contexts (López-Aladid *et al*. 2023).

Some of these differences in primer bias are underpinned by chemistry and currently lie beyond the reach of any post-hoc methods. Underrepresentation or complete omission of certain clades can be caused by template mismatches and even commonly used primers can miss roughly half of the community’s diversity in any given sample (Hong et al. 2009; Klindworth et al. 2013; Parada et al. 2016). Template genomic GC content can also skew recovered abundances (Aird et al. 2011; Laursen et al. 2017). These biases complicate cross-study comparison, yet studies sharing a primer set are routinely combined because the biases are held in common (Duvallet et al. 2017); what remains unresolved is comparison across primer sets, where each set carries its own biases and the datasets no longer share a feature space.

Positional issues, which stem from how sequenced reads become features, prevent cross-primer comparison altogether. Sequenced amplicons are denoised and reduced to a table of features before analysis. Early workflows clustered reads into operational taxonomic units (OTUs) at a fixed identity, usually 97% (Stackebrandt & Goebel, 1994). OTUs have now largely been displaced by denoising, with methods such as DADA2 and Deblur, modelling the sequencing error process to resolve reads to amplicon sequence variants (ASVs; exact sequences resolved at single-nucleotide resolution, equivalent to 100%-identity OTUs and also called exact or zero-radius sequence variants) (Callahan et al. 2016; Amir et al. 2017). Because an ASV’s identity is its exact sequence, ASVs are portable and comparable across independent studies without a shared reference or database, a central argument for their use (Callahan et al. 2017). However, this portability relies on a strictly conditional shared genetic window. Direct ASV merging is only possible where sequences derive from the same locus (Callahan *et al*. 2017). Studies using different but overlapping primer sets can only be combined after their ASVs are first trimmed to a standard, overlapping region. This issue arises from widely used primer sets binding at defined but non-identical positions on the gene (Klindworth *et al*. 2013). For instance, the commonly used set of primers for sequencing the V4 region (515F–806R; Caporaso et al. 2011) and V3–V4 (341F–805R; Herlemann et al. 2011) windows overlap only partially (Lee et al. 2023), and some pairings, such as V1–V2 (27F–338R; Lane, 1991; Amann et al. 1990) versus V4, do not share any overlap at all. The same organism is therefore recorded as a different ASV in each primer set, and where windows do not overlap there is no shared sequence to compare. Independently published multi-primer datasets therefore cannot be merged as sequences until they are brought into a common window. Callahan *et al*. (2017) prescribe exactly this trimming step, but currently there is no principled way to locate and trim to this shared window, and no means of confirming that such trimming has not distorted the community structure the comparison was intended to preserve.

Several solutions have been proposed to enable cross-study comparison, but none address the fundamental issue of position. Statistical batch-effect correction (MMUPHin, ConQuR, PLSDA-batch; Ma et al. 2022; Ling et al. 2022; Wang & Lê Cao, 2023) adjusts feature tables that already assume comparable features; closed-reference OTU picking certifies taxonomic identity rather than positional overlap (Rideout et al. 2014; Pollock et al. 2018); and phylogenetic placement (MaLiAmPi; Minot et al. 2023) reconciles studies by placing their sequences on a shared reference tree rather than on a shared gene window. None of these methods has become standard (Li et al. 2025), and none establishes that the studies are reading the same stretch of gene in the first instance; their respective strengths and limits are taken up in the Discussion.

To address this gap, we present SeCAT (Sequence Consensus Amplicon Trimming), a pipeline that enables the joint analysis of amplicon datasets generated with different primer pairs that amplify overlapping regions of the 16S rRNA gene. Using the sequences themselves, SeCAT first verifies that the datasets share a common gene segment and then returns a harmonised feature table anchored to that shared segment for downstream meta-analysis. SeCAT finds the largest window of coverage that is shared across studies by aligning their sequences to a common coordinate frame defined by a full-length 16S rRNA reference alignment. It then assesses, using several independent criteria, whether trimming to this window preserves community structure. It also quantifies, per study and per taxon, how trimming alters the data, so that inclusion decisions are informed. Studies that successfully pass these steps are trimmed and the ASVs are collapsed at exact identity into sequence units comparable across datasets (MetaASVs). A tiered validation framework, from sequence-level QC through alpha and beta diversity to co-occurrence networks, then allows the user to measure how well harmonisation preserves ecological signal and the between-study structure of the data, so that results from the pooled table can be interpreted against the original studies.

To validate SeCAT, we applied it to thirteen publicly available global marine 16S rRNA datasets (Supplementary Table S1). Each dataset was sequenced with one of three primer sets: 341F–805R (V3–V4; Herlemann et al. 2011), 515F(-Y)–926R (V4–V5; Parada et al. 2016) and 515F–806R (V4; Caporaso et al. 2011). Eleven of the datasets (682 samples after cleaning) satisfied the pipeline’s mapping and inclusion criteria and were harmonised into a single MetaASV table. The approach is general: any 16S meta-analysis drawing on different primer regions is a candidate, the reference database is interchangeable, and the logic extends to other markers such as 18S rRNA or ITS.

## Materials and Methods

### Overview and inputs

SeCAT is a Nextflow DSL2 pipeline (Di Tommaso *et al*. 2017) that harmonises independently published metabarcoding 16S rRNA datasets sequenced with different primer pairs into a single feature table of exact sequence variants (MetaASVs) comparable across studies.

The pipeline is executed in two phases (Figure 1). Phase 1 maps each study’s ASVs into the SILVA 138.2 full-length reference alignment, uses the alignment positions to establish a shared consensus window, builds a simulation null model and writes a diagnostic report on how each study degrades under both-end cropping. Phase 2 trims the selected studies to the consensus window, merges them into a combined feature table and validates the result. Every setting is customisable and is listed in Table 1.

**Figure 1.**
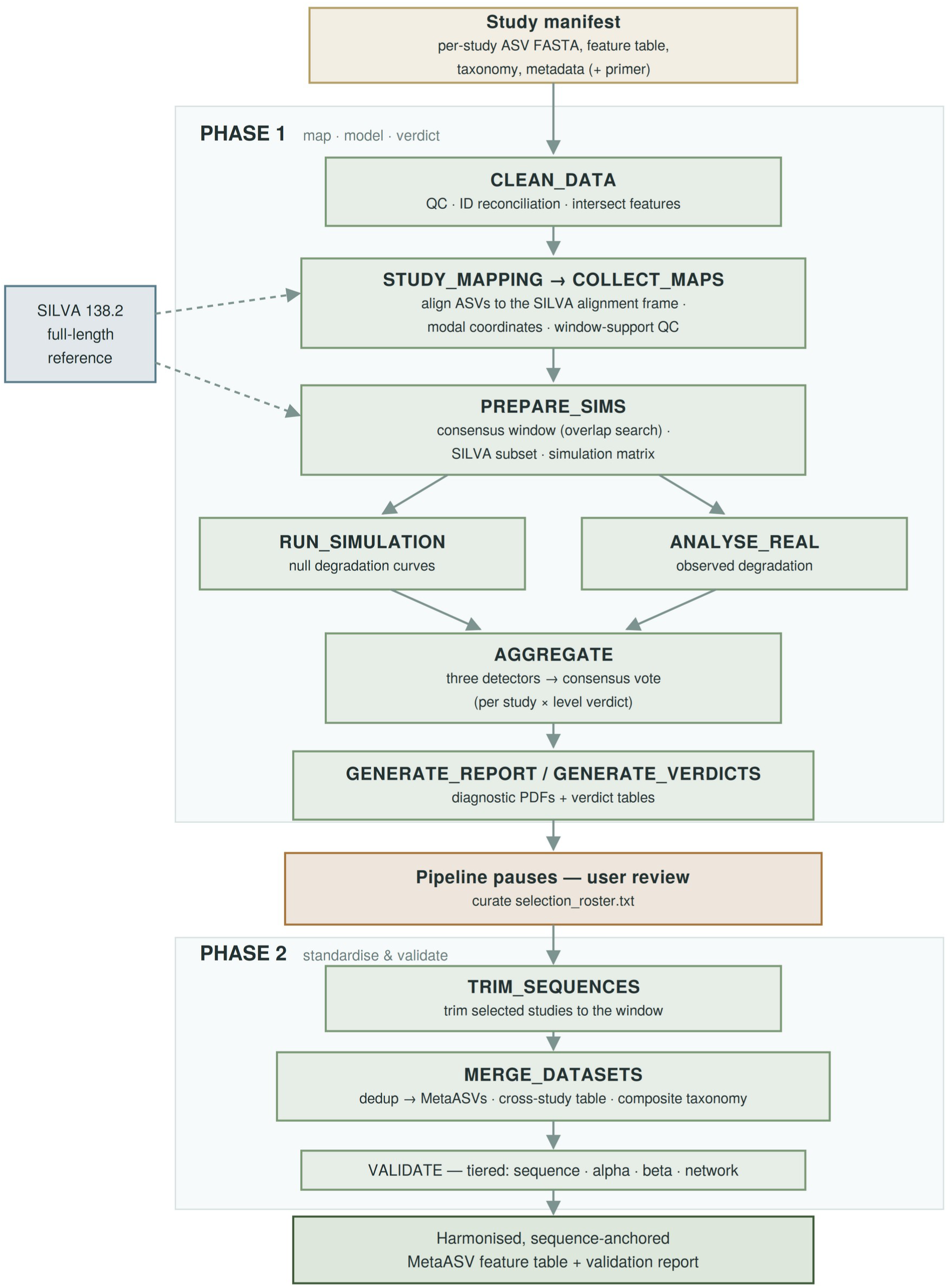
The SeCAT pipeline. A two-phase Nextflow DSL2 workflow. Phase 1 (map, model, verdict) maps each study’s ASVs onto a fixed SILVA 138.2 alignment frame, compares observed against null-model degradation, and records a per-study trimming verdict; the pipeline then pauses for the user to decide which studies to standardise. Phase 2 trims the selected studies to the consensus window, collapses identical trimmed sequences into MetaASVs with composite taxonomy, and runs tiered validation, yielding a harmonised MetaASV feature table and validation report. Boxes are pipeline processes; the SILVA 138.2 reference (blue) supplies the shared coordinate frame. ASV, amplicon sequence variant.

**Table 1.** SeCAT parameters used in this study. Parameter names are those of `params.yaml`; rows marked (fixed) are constants of the implementation. Each is described in the Methods subsection where it operates.

| Parameter | Value (validation dataset) | Description |
|---|---|---|
| analysis_mode | study | Coordinate mapping driven by each study's own ASV sequences (the only supported mode) |
| Minimum samples per study | 3 (fixed) | Studies with fewer samples are dropped (CLEAN_DATA) |
| Sample-reconciliation floor | 50% (fixed) | Study excluded if fewer than half its samples reconcile between feature table and metadata |
| min_sample_depth | 10,000 reads | Samples below this depth are dropped (CLEAN_DATA) |
| min_asv_reads / min_asv_prevalence | 50 reads / 2 samples | ASVs with fewer than 50 reads in fewer than 2 samples are removed, recomputed over surviving samples |
| remove_organelles | true | Chloroplast and mitochondrial ASVs removed (optional; enabled by default) |
| reference_alignment_mode | subset | SILVA reference subset used for profile alignment (values: subset, full) |
| reference_subset_size | 5,000 (seed 123) | Random SILVA subset size for STUDY_MAPPING alignment |
| use_all_asvs | true | All of a study's ASVs used in coordinate mapping |
| min_overlap | 50 columns | Minimum amplicon-window overlap (alignment columns) for a compatibility-graph edge |
| consensus_optimization_threshold | 20% | Minimum window widening required to drop a “constrictor” study |
| min_consensus_studies | 3 | Floor on studies retained in the consensus clique |
| sim_max_silva_subset | 10,000 (reservoir seed 42) | Shared SILVA community pool for all simulations |
| num_simulations | 100 | Simulated-community replicates per study |
| maxForks | 50 | Concurrent simulation replicates / process throttle |
| simulation_community_size | 0 (= each study’s ASV count) | Null-community size; 0 matches the study’s ASV count, with sequence families emitted whole |
| sim_structure_matched_null | true | Null community reproduces each study’s amplicon window, family-size distribution and within-family divergence (measured once per study in PREPARE_SIMS) |
| sim_cluster_identity | 0.97 | Identity defining a sequence family (near neighbour) when that structure is measured |
| sim_max_template | 1,500 | Template ASVs per study used to measure that structure |
| sim_abundance_model | lognormal (μ = 5, σ = 2) | Relative-abundance model for simulated communities |
| trim_step_mode | absolute | Trim depth in fixed base-pair increments |
| trim_increment | 100 bp | Trim step size |
| max_absolute_trim_steps | 2,000 | Upper bound on trim steps |
| consensus_buffer_steps | 20 | Extra steps beyond the candidate trim point |
| sim_add_pcr_bias | true (sim_pcr_gc_bias 0.65; sim_pcr_optimal_gc 0.50; sim_pcr_cycles 25; floor 10%) | efficiency = 1 − 0.65 × ((GC − 0.50)/0.15)², floored at 10%, over 25 cycles |
| sim_add_errors | true (sim_error_rate 0.003 at the 5′ end, rising linearly to 2× at the 3′ end; sim_error_position_bias true) | Per error: 90% substitution (transition 0.70, each transversion 0.15), 5% insertion, 5% deletion |
| sim_add_chimeras | false | Chimera simulation disabled by default |
| VSEARCH mapping identity | 0.97 (fixed) | Identity at which trimmed sequences are matched by VSEARCH (usearch_global) to their step-0 baseline centroids: the conventional 97% OTU threshold, which tolerates simulated sequencing error |
| changepoint_scan_min / changepoint_scan_max / changepoint_bootstrap_n | 0.4–2.4 (step 0.1); 50 bootstrap replicates | penalty = multiplier × log n; multiplier chosen by bootstrap cross-validation |
| distance_cutoff_threshold | 0.20 (≈ 20% turnover) | Step-wise Bray–Curtis change flagging degradation |
| null_model_p_threshold | 0.05 | Empirical p-value threshold for null-model detection |
| null_model_min_consecutive | 2 steps | Consecutive p-value steps below threshold to trigger a null-model degradation call |
| Initial trim excluded | first 10 columns (fixed) | Excluded from all three detectors |
| verdict_consensus_tolerance_bp | 500 bp | Tolerance window for combining detector thresholds |
| bc_ceiling_threshold | 0.20 | Hard Bray–Curtis ceiling (HARD_FAIL) |
| retention_floor_threshold | 0.70 | Hard taxon-retention floor (HARD_FAIL) |
| Minimum trimmed length | 50 bp (fixed) | Trimmed sequences below this length are discarded |
| Minimum trim yield | 50% (fixed) | Studies below this yield excluded at trimming |
| MetaASV dedup identity | 100% (fixed) | Exact-identity dereplication into MetaASVs |
| Taxonomy composite score | 0.70 × confidence + 0.30 × length (fixed) | min–max normalised, pre-trim; ties broken on confidence, then length |
| plot_dpi | 600 | Diagnostic-figure resolution |
| auto_trim | false | Pipeline pauses before trimming for analyst review |
| selection_mode | roster | Study inclusion from a curated selection_roster.txt |
| validation_levels | ASV, Genus, Family | Levels at which harmonisation is validated |
| SPIEC-EASI feature filter | 0.05% relative abundance (fixed) | Taxa retained for co-occurrence-network inference |

For each study SeCAT requires four inputs: a FASTA file of ASV sequences, an ASV-by-sample read-count table (feature table), both generated after denoising and chimera removal, a table of taxonomy assignments and a sample metadata file. File paths are given in a per-study manifest (see the README). Additionally, it requires a reference full-length alignment (SILVA 138.2 SSURef NR99; Quast et al. 2013), to define a common coordinate frame for ASV mapping across studies. SILVA is used because its SSURef alignment places every sequence in a fixed 50,000-column frame spanning bacteria, archaea and eukaryotes, so column coordinates are comparable across studies and domains; the reference is a run parameter (`reference_db`) and any fixed-coordinate full-length alignment can be substituted. Process and parameter names are set in monospace throughout. The first process, `CLEAN_DATA`, prepares inputs by filtering ASVs by abundance, removing chloroplast and mitochondrial sequences (`remove_organelles`, enabled by default), dropping samples with no reads, and removing those studies with less than a minimum number of samples (Table 1). Sample identifiers are reconciled between metadata and feature table before each study’s inputs are intersected to a common ASV and sample set.

### Reference coordinate mapping

`STUDY_MAPPING` (Figure 1) aligns each study’s ASVs in a common reference frame using DECIPHER v2.30.0 (Wright, 2015, 2016). To bound memory and runtime, ASVs from each study are first aligned to one another with `AlignSeqs` into a study-specific multiple sequence alignment (a profile). This profile is then aligned to the SILVA reference alignment using `AlignProfiles` (Figure 2a). The reference is a fixed random subset of SILVA (Table 1). Studies exceeding 65,000 ASVs (Table 1) are split into chunks, and each chunk is aligned to the reference in the same way (Figure 2a). After profile alignment, columns arising from insertions relative to the reference are removed, so that every alignment spans exactly the reference alignment columns. By treating the reference coordinate system as fixed, SeCAT anchors every ASV to the same positional framework regardless of the hypervariable region covered.

**Figure 2.**
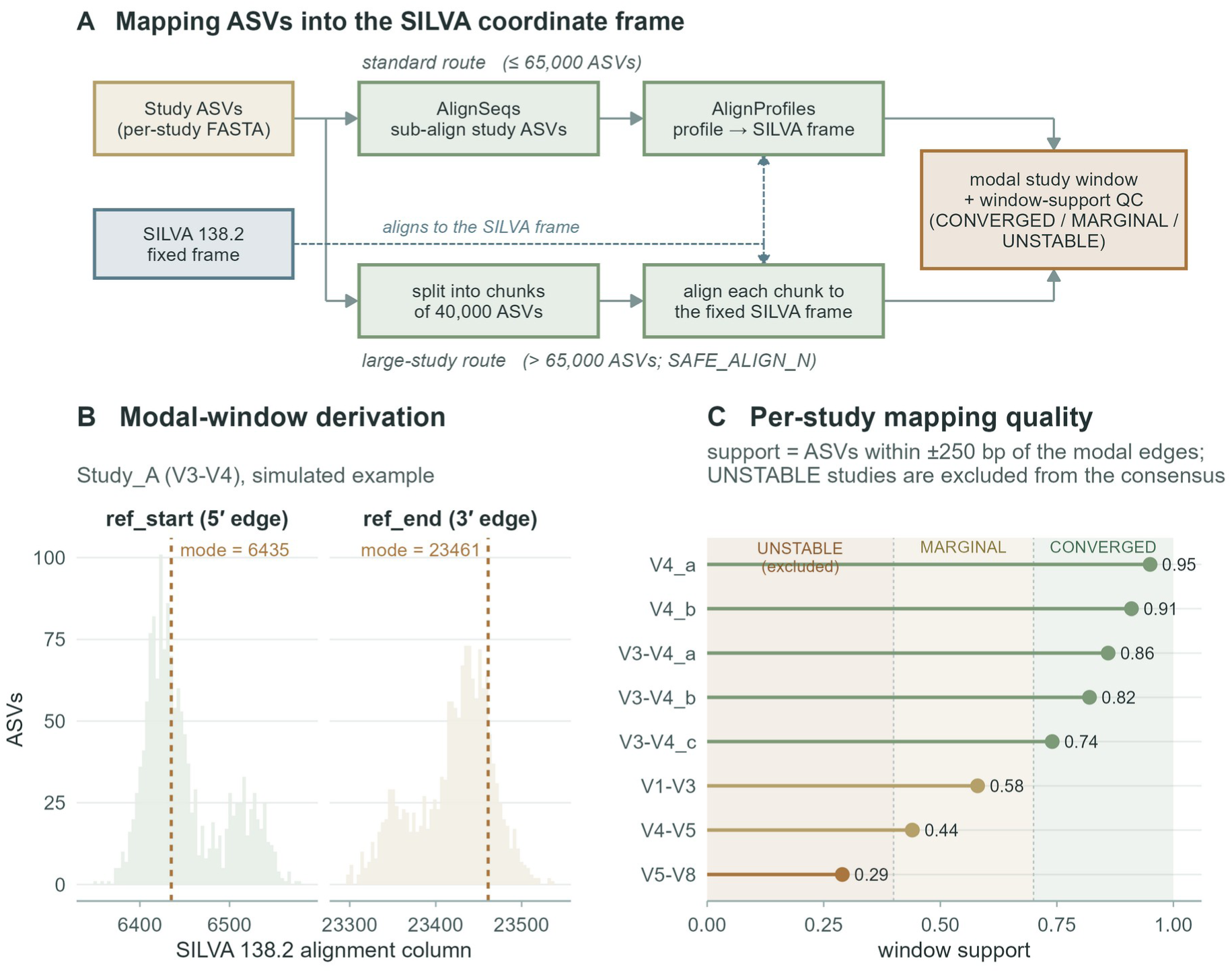
Mapping ASVs into the SILVA 138.2 coordinate frame. (A) ASVs are aligned onto a fixed SILVA 138.2 column frame by a standard route (≤ 65,000 ASVs) or, for larger studies, a chunked route (batches of 40,000; `SAFE_ALIGN_N` = 65,000); both feed modal-window derivation and window-support quality control (QC). Data shown are a simulated illustrative example. (B) Per-ASV 5′ (`ref_start`) and 3′ (`ref_end`) alignment-column positions for a simulated representative study; dashed lines mark the modal edges defining the study window. (C) Per-study window support for eight simulated studies (the fraction of a study’s ASVs within ±250 alignment columns of both modal edges), classified CONVERGED (≥ 0.70), MARGINAL (≥ 0.40) or UNSTABLE (< 0.40); UNSTABLE studies are excluded from the consensus. ASV = amplicon sequence variant.

For each ASV, the reference start and end position are read from the merged alignment as the first and last non-gap positions relative to the reference columns, and the study’s amplicon coordinates are derived from the statistical mode across all mapped ASVs. Modal derivation is used because amplicons from a single primer pair produce sharply peaked coordinate distributions (illustrated with simulated data in Figure 2b), whereas length variants could skew mean positions considerably. Each study is then scored for how tightly its ASVs converge on the modal window. The proportion of ASVs whose start and end fall within a fixed band of the modal coordinates classifies the study as CONVERGED (≥ 0.70), MARGINAL (≥ 0.40) or UNSTABLE (< 0.40) (Figure 2c). The support band and both class boundaries are tuneable parameters (`mapping_support_band`, `mapping_support_warn`, `mapping_support_fail`); the defaults are heuristic values chosen to separate clear convergence from ambiguous mapping, and the sensitivity of the downstream detectors to parameter choice is characterised in Supplementary Note S1. Studies flagged UNSTABLE, whose ASVs do not agree on a single window, are excluded from consensus determination and downstream analysis (“Consensus region determination” section). `COLLECT_MAPS` concatenates the per-study coordinate table.

### Consensus region determination

The global consensus region is determined by the `PREPARE_SIMS` process (Figure 1), in which studies are treated as nodes in a compatibility graph and these nodes are joined by an edge wherever their amplicon windows overlap by a set minimum overlap value (Table 1; Figure 3a,b). A greedy search grows a set of mutually overlapping studies and retains the largest of these sets. By construction every member of said set overlaps every other by at least `min_overlap`. A recursive optimisation step then tests whether removing a boundary “constrictor” study, one whose amplicon boundary lies inside the pooled window and thereby limits the shared region, widens the shared window by more than a set threshold (Table 1). Where it does, this constrictor study is dropped and the search recurses, subject to a floor on the number of retained studies (`min_consensus_studies`, Table 1). The final consensus region is the intersection of the surviving set, from the maximum start to the minimum end and is written with the included and excluded study lists (Figure 3a).

**Figure 3.**
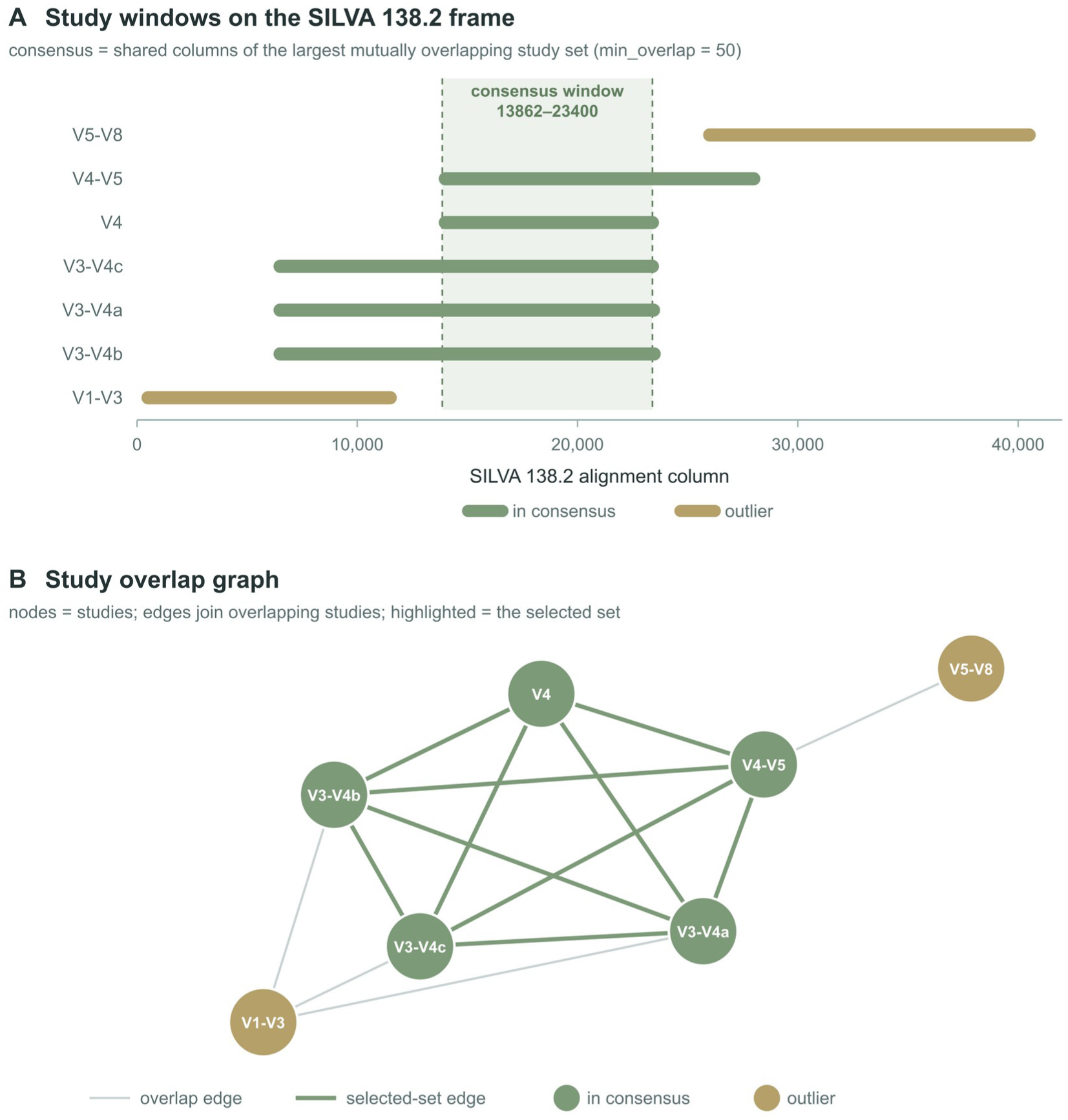
Selection of the consensus window. (A) Each bar is one study’s mapped window on the SILVA 138.2 frame (alignment columns); the shaded band is the consensus window, the shared columns of the largest set of mutually overlapping studies (`min_overlap` = 50 columns). (B) Study overlap graph: nodes are studies, edges join studies overlapping by ≥ `min_overlap`, and the selected set is highlighted. In both panels, green denotes in-consensus and gold denotes outlier studies. ASV, amplicon sequence variant.

### Simulation-based null model

To decide whether trimming a study to the consensus window distorts its community more than expected, SeCAT compares the study’s observed degradation with an empirical null distribution. The null is built from simulated communities that reproduce the study’s amplicon window and sequence-family structure and are trimmed in the same way, so they show the degradation expected from shortening the sequences alone; excess degradation in the real data therefore points to study-specific signal in the trimmed regions. `PREPARE_SIMS` sets up these simulations and `RUN_SIMULATION` executes them. `PREPARE_SIMS` first draws a random subset of the reference database by single-pass reservoir sampling (Knuth, Algorithm R), written once so that every simulation worker draws from an identical community pool (Table 1). It also measures, once per study, the sequence-similarity structure of that study’s aligned ASVs (pairwise identities among up to 1,500 template ASVs; `sim_max_template`): the fraction with a near neighbour at or above the clustering identity (`sim_cluster_identity`), the size distribution of the resulting sequence families and the per-column variability of the amplicon window. It then generates a simulation task matrix with one row per study and a random seed. Trim depth advances in fixed increments to a bound set by each study’s amplicon length plus a margin, so that degradation curves extend past the candidate trim point (Table 1).

`RUN_SIMULATION` independently generates one *in silico* community for each study and seed, then analyses its null degradation curve per replicate, following the amplicon-simulation structure of Grinder (Angly *et al*. 2012). A community is initialised as a structure-matched null (`sim_structure_matched_null`; Table 1): independent seed sequences are drawn without replacement from the SILVA subset and restricted to the study’s amplicon window, and each seed is expanded into a family whose size and within-family divergence are sampled from the study’s own measured structure, so that the null contains the close relatives that trimming can merge, up to the study’s ASV count (`simulation_community_size`). Relative abundances are then assigned from a lognormal distribution (Table 1), the species abundance structure expected for prokaryotic communities (Curtis et al. 2002; Shoemaker, Locey & Lennon, 2017). Three artefact layers are then applied. First, GC-dependent PCR amplification bias is applied to reproduce the quantitative skew documented for 16S amplicons (Aird *et al*. 2011; Laursen *et al*. 2017; Pinto & Raskin, 2012). Second, sequencing errors are simulated using a position-dependent model. The substitution rate and its increase towards the 3′ end of reads follow empirical MiSeq error profiles (Schirmer et al. 2015, 2016), the platform on which most publicly archived 16S datasets were generated. Transitions are favoured over transversions, to reflect the bias associated with polymerase misincorporation during amplification (Potapov & Ong, 2017). Third, chimera formation can be simulated but is disabled by default, as chimeras are routinely removed by modern denoisers (Haas *et al*. 2011). Each simulated community is then trimmed stepwise from both ends towards the selected window. At each trimmed step, sequences are matched back to the untrimmed baseline sequences cut to the same window using VSEARCH, at a fixed identity threshold (Table 1; Rognes et al. 2016). Bray–Curtis dissimilarity to baseline (Bray & Curtis, 1957) and taxonomic retention are then computed at each taxonomic rank (phylum, class, order, family, genus and ASV). Taxonomic retention is defined as the fraction of baseline taxa still detected after trimming at each rank. Real and simulated communities are trimmed identically, using the same fixed increments, steps and starting coordinates, so their degradation curves are directly comparable. Per-seed results are pooled into the empirical null distribution of change expected under trimming alone.

### Real data analysis and degradation detection

Each study’s community is analysed by the `ANALYSE_REAL` process (Figure 1). Stochastic operations are seeded from the study identifier for reproducibility. Sequences are trimmed inwards from both ends in fixed increments (Table 1). The total trimming depth is the sum of the distances from the study’s original 5′ and 3′ amplicon boundaries to the consensus boundaries, plus an additional margin (Table 1). At each step, the increment is divided between the two ends in proportion to how far each lies from the consensus. This ensures the end with the larger overhang is trimmed more and both ends reach the consensus boundary together. Per-taxon abundance trajectories are recorded for the diagnostic report. `AGGREGATE` then pools both real and simulated curves and, for each taxonomic level, applies three independent detectors to the real Bray–Curtis degradation curve before combining them by a consensus vote: PELT changepoint, empirical distance cutoff and null-model comparison (Figure 4). This produces a per-study, per-degradation status that is distinct from the later study-inclusion decision. A changepoint detector (PELT; `changepoint::cpt.meanvar`; Killick et al. 2012; Killick & Eckley, 2014) identifies the trim depth at which the mean and variance of the dissimilarity curve shift abruptly. Because its sensitivity depends on a penalty parameter, the curve is bootstrap-resampled across a scan of penalty multipliers (Table 1) and the multiplier whose changepoint location is most reproducible across resamples is adopted. An empirical distance cutoff flags degradation wherever the stepwise change in dissimilarity exceeds a set threshold (0.20 by default; Table 1). A null-model comparison computes at each step an add-one-smoothed empirical p-value, (k + 1)/(n + 1), where k is the number of simulation replicates whose step-wise change in Bray–Curtis dissimilarity equals or exceeds the observed change and n is the number of replicates (the Monte-Carlo empirical-p convention; North et al., 2002). Degradation is triggered when the p-value stays below threshold (0.05 by default) for a set number of consecutive steps (two by default; Table 1; Figure 4). The three triggers are combined into a consensus status with a tolerance window (Table 1). The result is CONFIRMED where at least two agree within tolerance, WARNING_SINGLE when exactly one method triggers, or when multiple methods trigger without agreement; and NONE when no method triggers. Two hard quality gates sit outside the vote and override it, a taxon retention floor and a Bray–Curtis ceiling, with a breach of either producing a HARD_FAIL. The per-method thresholds, consensus status, outlier flag and required trim depth are summarised by the GENERATE_VERDICTS process.

**Figure 4.**
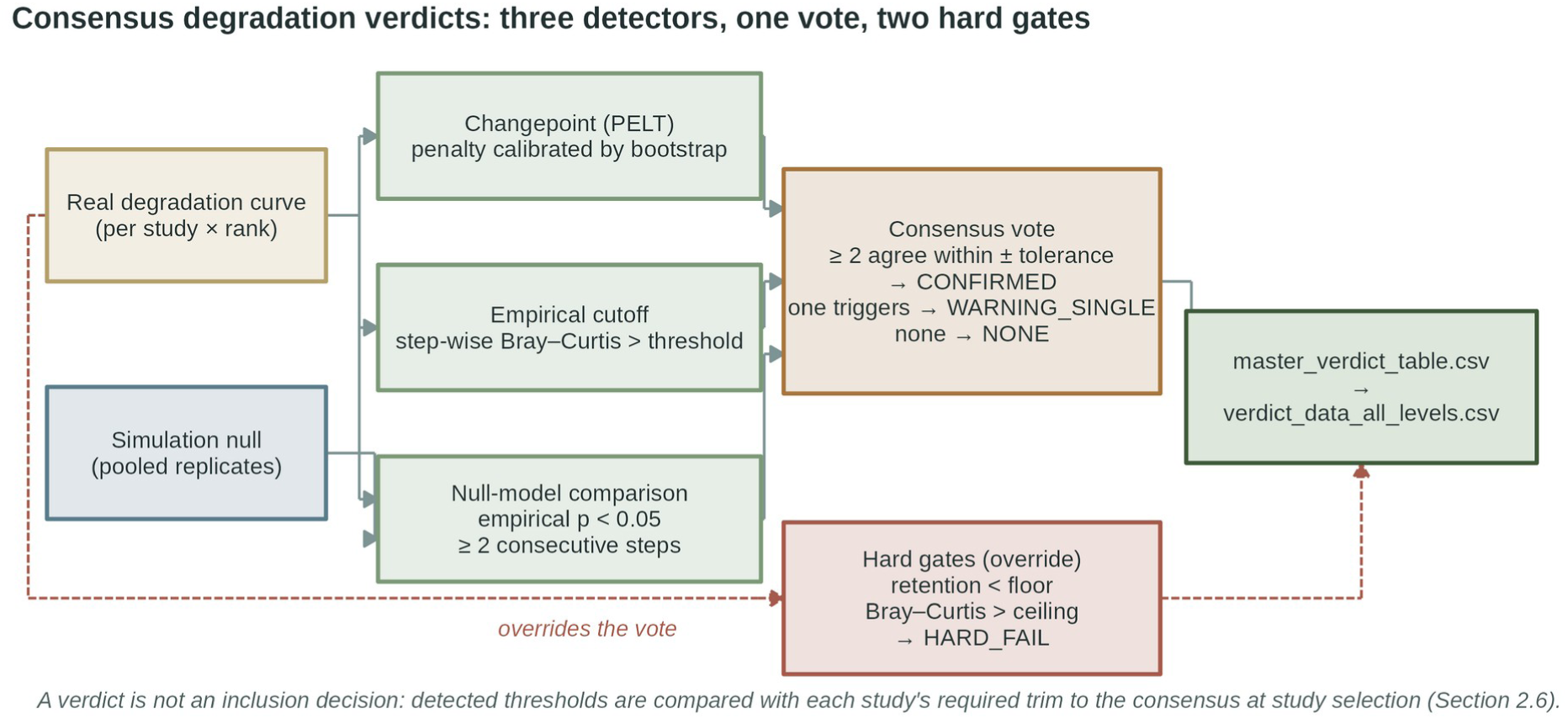
Consensus degradation verdicts. Each study’s real Bray–Curtis degradation curve is assessed per taxonomic rank by three independent detectors: a bootstrap-calibrated PELT changepoint, an empirical step-wise distance cutoff, and a null-model comparison against the pooled simulation replicates. Detector thresholds are combined by a consensus vote (CONFIRMED where at least two agree within tolerance; WARNING_SINGLE where one triggers; NONE otherwise). Two hard quality gates, a taxon-retention floor and a Bray–Curtis ceiling, sit outside the vote and override it (HARD_FAIL). Statuses and thresholds are written to `master_verdict_table.csv`. A verdict is not an inclusion decision: thresholds are compared with each study’s required trim to the consensus at study selection.

### Diagnostic reporting and study selection

`GENERATE_REPORT` (Figure 1) produces a diagnostic for each study at each taxonomic level with four graphical panels that help the user to decide which studies to include in downstream analysis (Figure 5): the real-data degradation curve against the simulation envelope, the taxonomic retention curves, the taxa whose relative abundances change most across trim depths, and core taxon trajectories or a pass/fail summary where these are unavailable. The ASV-level page of each per-study report for the demonstration run is provided as Supplementary Data S1. A master summary collates an amplicon-alignment map, a forest plot of safe trim thresholds, a study-inclusion guide and a comparison of the three detectors (Supplementary Data S2). Studies to carry forward are listed in a selection roster, one per line, keeping each inclusion decision explicit and auditable.

**Figure 5.**
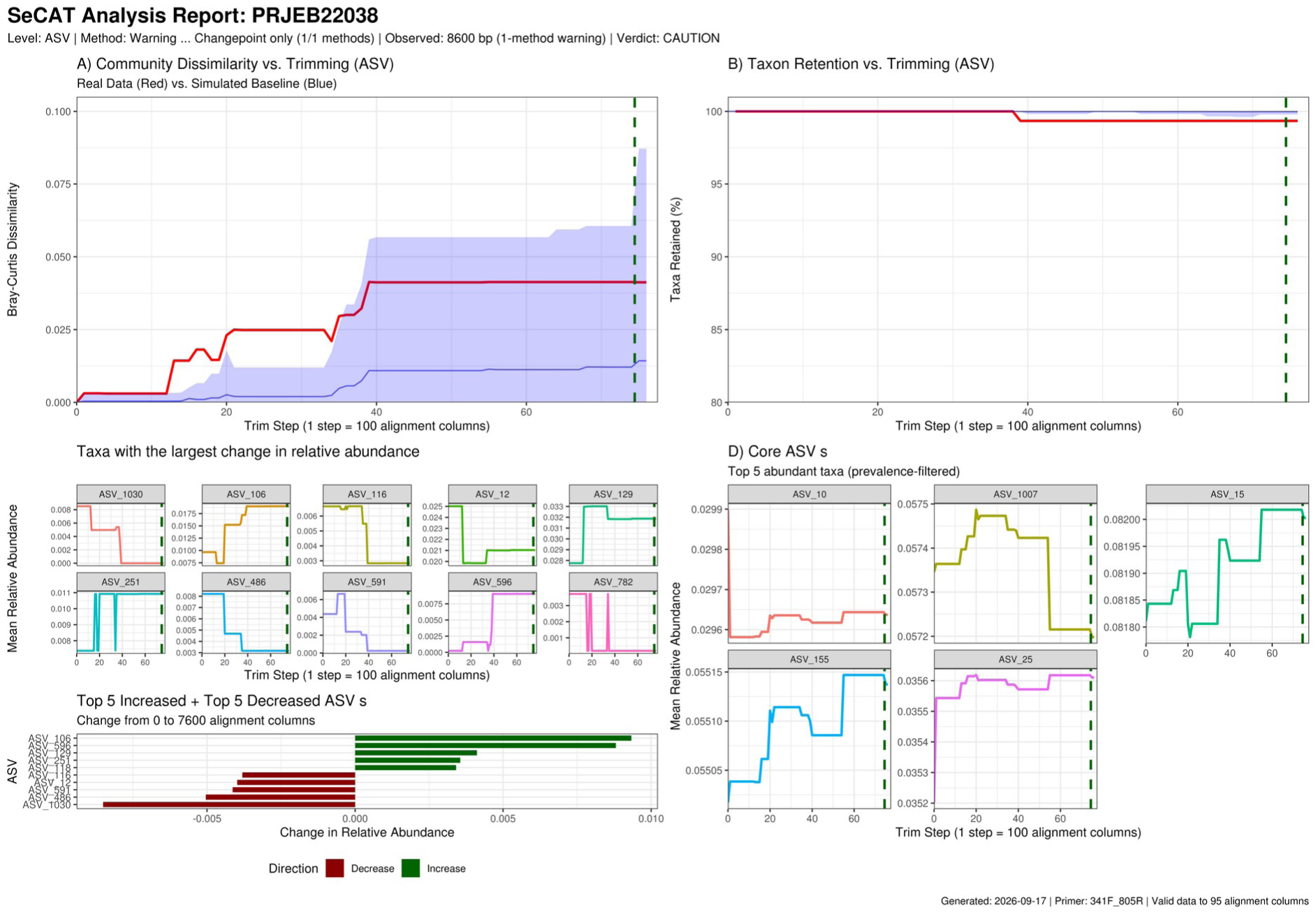
Exemplar per-study diagnostic report page (PRJEB22038, 341F–805R; ASV level): (A) real-data degradation curve against the simulated baseline with the detected threshold; (B) taxon retention with trimming; (C) taxa with the largest change in relative abundance and the top increases and decreases at the trim point; (D) core taxon trajectories. Panel headers report each detector’s raw status at the detected changepoint; the overall verdict (Supplementary Table S4) additionally accounts for whether the detected change lies beyond the study’s required trim, so a panel may display CAUTION while the study passes overall. The corresponding page for every study and taxonomic level is provided in Supplementary Data S1.

### Trimming, MetaASV merging and outputs

Standardisation is the second phase of SeCAT (`STANDARDIZE`). `TRIM_SEQUENCES` (Figure 1) extracts the intersection of each study’s mapped range with the consensus window from each study’s stored DECIPHER aligned FASTA and removes the alignment gaps. ASVs that are below a minimum length (50 bp by default) and studies that contain too few sequences after trimming (50% by default) are excluded (Table 1). MERGE_DATASETS pools the trimmed ASVs sequences from all selected studies and deduplicates them at exact 100% identity (exact string identity; `Biostrings::unique` on the pooled trimmed sequences) into MetaASVs. MetaASVs are defined by their shared-window sequence and so are directly comparable between primer sets. Every original ASV is linked to its MetaASV in a provenance table; within each study, ASV counts are summed onto MetaASVs, and the per-study tables are combined by a full outer join on MetaASV, with a MetaASV unobserved in a study’s samples filled with zero rather than NA so that the output remains a count matrix accepted by downstream tools; such zeros denote non-detection (see Limitations). Taxonomy for each MetaASV is assigned by a composite score combining classifier confidence (the per-ASV score supplied with each study’s input taxonomy table: per cent identity for the BLAST assignments used here, or bootstrap support at the assigned rank for naïve-Bayes and IDTAXA classifiers, rescaled to 0–1 whichever convention the classifier uses) and pre-trim sequence length (Table 1), with ties broken on confidence, then length (Figure 6). The score for each candidate ASV is 0.70 × classifier confidence (scaled 0–1) + 0.30 × pre-trim sequence length (min–max normalised among the ASVs collapsing into that MetaASV); the highest-scoring ASV donates its taxonomy. Where every contributing ASV has the same pre-trim length the length term is zero for all and confidence alone decides. The 0.70/0.30 weighting gives the length term at most 0.43 confidence points of leverage in principle; its realised effect on the demonstration dataset is quantified in the Results (the donor changed in 0.9% of MetaASVs, never at a confidence cost above 6.6 points). Because confidence values are compared across studies, taxonomy should be assigned to all input studies with the same classifier and reference database; scores from different classifiers (for example BLAST identity and naïve-Bayes bootstrap support) are not interchangeable. Sample metadata are then combined by prefixing sample identifiers and adding a `StudyID` provenance column (the manifest `study_name`). Both the trimmed (post-consensus) and untrimmed (pre-consensus) datasets are written as tab-delimited feature, taxonomy and metadata tables with a representative sequence FASTA and the ASV-to-MetaASV map, importable into phyloseq (McMurdie & Holmes, 2013).

**Figure 6.**
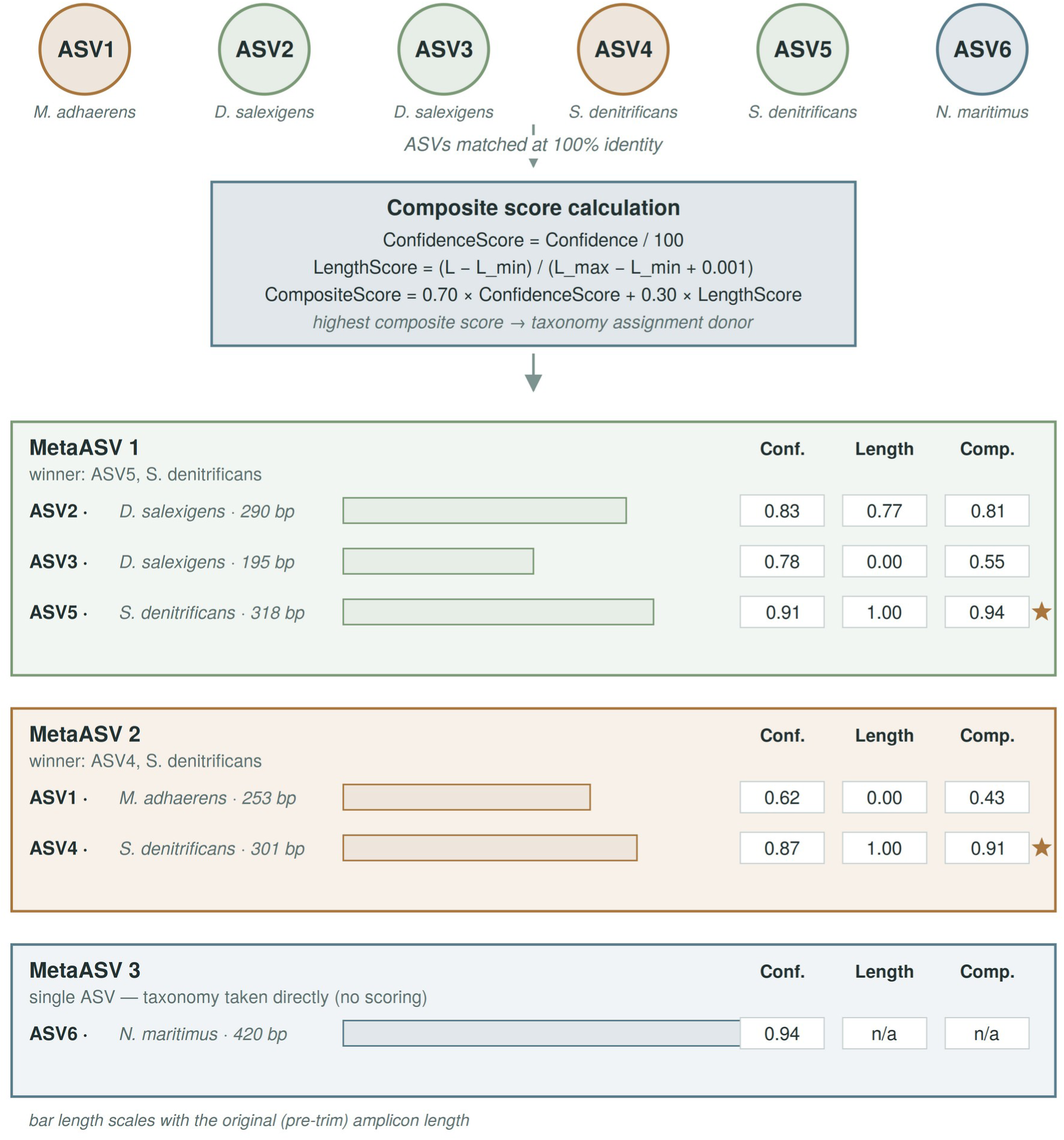
MetaASV construction and composite-score taxonomy. Trimmed ASVs matched at 100% identity collapse into MetaASVs, with read counts summed (circles coloured by MetaASV). Where contributing ASVs disagree on taxonomy, the donor is the ASV with the highest composite score (0.70 × confidence + 0.30 × length, length normalised within the MetaASV). Each panel shows contributing ASVs as bars (width ∝ original amplicon length) beside their Confidence, Length and Composite scores; ★ marks the donor. MetaASV 3 shows the single-ASV case: still assigned a MetaASV identifier, with taxonomy taken directly and no scoring (n/a). Taxa and scores are illustrative. ASV, amplicon sequence variant.

### Validation framework of MetaASVs

`VALIDATE` (Figure 1) runs automatically after merging and compares the pre-standardisation (cleaned, untrimmed) and post-standardisation (trimmed, MetaASV) datasets at each level in `validation_levels` parameter (e.g., ASV, Genus and Family), scoring each metric against a fixed criterion; the criteria are reported with the results. Sequence-level standardisation is checked through the coefficient of variation in sequence length, the shift in median GC content, the consensus coordinates and trim success (every rostered study returning SUCCESS from TRIM_SEQUENCES). Alpha-diversity preservation is quantified by a two-way random-effects, absolute-agreement, single-measurement intraclass correlation (ICC(A,1); irr v0.85, Table 2; reported following Koo & Li, 2016) between the pre- and post-standardisation values of observed richness, Chao1, ACE, Shannon, Simpson and Pielou evenness, and by Spearman’s ρ of the same indices. Beta-diversity structure is assessed between the pre- and post-standardisation Bray–Curtis distance matrices by a Pearson Mantel test (999 permutations; Mantel, 1967) and by symmetric Procrustes (mZ) with PROTEST (999 permutations; Gower, 1975; Peres-Neto & Jackson, 2001), together with a PERMANOVA (`adonis2` on `StudyID`, 999 permutations) comparing the variance explained by study before and after standardisation, which should not increase (Anderson, 2001; McArdle & Anderson, 2001). Within-group dispersion is tracked by the relative change in mean distance-to-centroid (`betadisper`; Anderson, 2006). Rank-abundance and abundance concordance are measured by Spearman’s ρ, and cross-study comparability by the MetaASV sharing rate at ASV level and mean pairwise Jaccard at genus and family. SPIEC-EASI networks (neighbourhood selection with StARS model selection; Kurtz *et al.,* 2015) are inferred through the microeco v2.3.0 `trans_network` framework (Liu et al. 2021; Table 2) on taxa filtered at a low relative abundance threshold (Table 1). The scored metrics are the change in vertex and edge counts and the retention of hub taxa (the top decile of vertices by degree, matched through the ASV to MetaASV map at ASV rank), with density, mean degree and transitivity reported. Topology and greedy modularity (`cluster_fast_greedy`) are computed with igraph (Csárdi & Nepusz, 2006), with networks exported as GraphML. Multivariate tests use vegan v2.7-5 (Oksanen et al. 2025). Results are summarised in a machine-readable file (value, criterion and pass status per metric) with diagnostic figures.

**Table 2.** Software and versions.

| **Software / package** | **Version** | **Role in SeCAT** |
|---|---|---|
| **Nextflow (DSL2)** | 26.04.3 | Workflow manager |
| **Singularity** | 4.5.0 (singularity-ce) | Container runtime (recommended on HPC) |
| **Docker** | not used in this run (supported alternative) | Container runtime (alternative) |
| **R** | 4.3.2 | Analytical environment (all pipeline logic) |
| **VSEARCH** | 2.23.0 | 97% identity mapping in degradation analysis |
| **DECIPHER** | 2.30.0 | Reference-frame profile alignment |
| **changepoint** | 2.3 | PELT changepoint detection |
| **vegan** | 2.7.5 | PERMANOVA, Mantel, Procrustes, betadisper |
| **microeco** | 2.3.0 | trans_network co-occurrence framework |
| **SpiecEasi** | 1.1.2 | Compositionally robust network inference |
| **igraph** | 2.3.3 | Network topology and modularity |
| **irr** | 0.85 | Intraclass correlation (alpha diversity) |
| **Biostrings, ape, data.table** | 2.70.3 / 5.8.1 / 1.18.4 | Sequence and data handling |
| **ggplot2, patchwork** | 4.0.3 / 1.3.2 | Diagnostic figures |

### Application to marine 16S rRNA datasets

To demonstrate SeCAT, thirteen marine 16S rRNA datasets sequenced with three primer sets spanning V3 to V5 region were downloaded from the European Nucleotide Archive (ENA). The description and accession number of each project are listed in Supplementary Table S1. The samples were pre-processed using DADA2 v1.10.0 (Callahan et al. 2016), after primers trimming by Cutadapt v1.18 (Martin, 2011). Each sequencing project was analysed separately with the same filtering parameters (`maxEE` = c(2,2); `truncLen` = c(250,220)). The taxonomic classification was performed using BLAST v2.17.0+ (Camacho et al. 2009) with the blastn algorithm (e-value = 0.0001) and the SILVA database (SSU Ref 138 NR99) as a reference. Samples with fewer than 10,000 reads were discarded and only ASVs with at least 50 reads in total and present in at least two samples were retained (thresholds adapted from Metz et al. 2022, who required three samples). The feature table, taxonomy assignation and metadata were then processed with SeCAT with the parameter values in Table 1. The pipeline was run on the JASMIN LOTUS2 SLURM cluster (the UK data analysis facility for environmental science). After the Phase 1, selected studies (Supplementary Table S1) were then trimmed to the consensus window and merged into MetaASVs and harmonised and original datasets were compared with the validation framework. Dataset composition, consensus coordinates and validation outcomes are reported in the Results.

### Implementation

SeCAT is implemented in Nextflow DSL2 (Di Tommaso et al. 2017) with one process per stage and all analytical logic in R (R Core Team, 2023); it is containerised and runs unchanged from HPC schedulers to workstations. Three executor profiles (slurm, sge, local) each combine with a container profile (singularity, which is compatible with Apptainer and recommended on HPC (Kurtzer et al. 2017), or docker). Operational detail (caching and resumption, per-process resources, container bind paths) is documented in the README. The pipeline was built and tested with Nextflow 26.04.3 (DSL2) and the Singularity (version 4.5.0) and Docker container runtimes. The version of R and every wrapped tool and R package used is given in Table 2. These versions are fixed by the container build (rocker/r-ver 4.3.2 base image, CRAN packages from a Posit Package Manager snapshot dated 2026-08-01, Bioconductor 3.18, SpiecEasi by release tag), so Table 2 lists the versions resolved in the published image.

## Results

### Coordinate mapping and degradation testing define a 12-study consensus window

SeCAT was demonstrated on a collection of thirteen publicly available marine 16S rRNA datasets drawn from studies worldwide, predominantly open-ocean and coastal water-column communities with two studies also including host-associated (sponge and kelp-biofilm) samples (Supplementary Table S1). These studies utilised one of three primer sets (341F-805R, 515F(-Y)-926R and 515F-806R; V3-V5). All datasets satisfied the cleaning thresholds (Table 1) and entered coordinate mapping (Table 3). Sample-level cleaning removed 276 of 1,083 input samples (25.5%) across the thirteen datasets, dominated by low-depth samples in PRJEB27168 (61 of 66 removed) and PRJEB50983 (80 of 401). Each study’s ASVs (25,402 in total; Table 3) were aligned to the SILVA 138.2 reference frame and reduced to a modal amplicon window (Table 3; Figure 7a).

**Figure 7.**
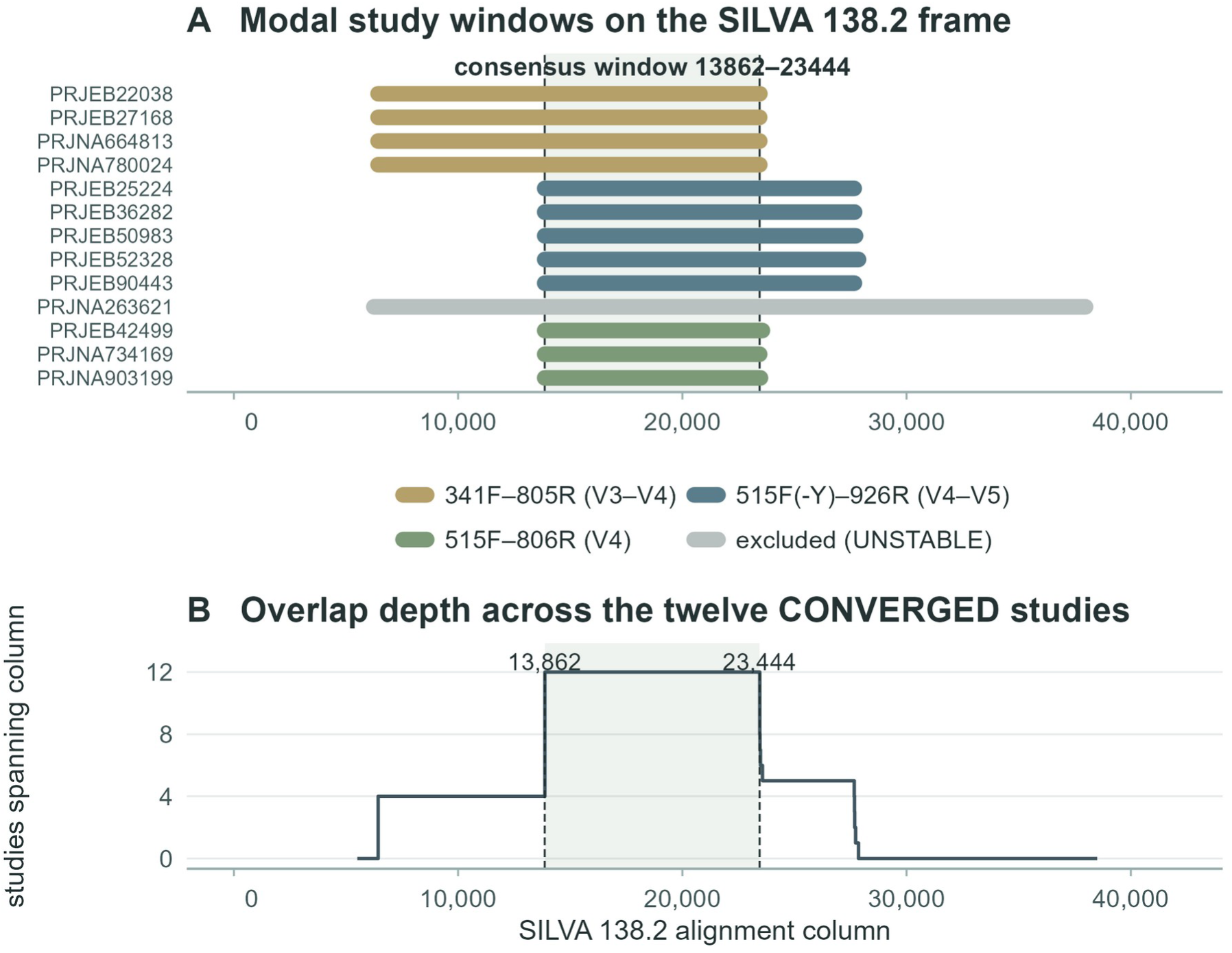
The real-data consensus window for the validation dataset. (A) Modal amplicon windows for the thirteen mapped studies on the SILVA 138.2 alignment frame, coloured by primer set, with the consensus window (columns 13,862–23,444; 9,582 columns) shaded; the excluded study (PRJNA263621, window support 0.24, UNSTABLE mapping) is greyed. (B) Column-wise overlap depth across the twelve CONVERGED studies; the plateau where all twelve overlap defines the consensus window, bounded below by the 515F primer start (13,862) and above by the shortest V4 amplicon end (23,444).

**Table 3.** Validation dataset: coordinate mapping, window support and inclusion. Modal windows are SILVA 138.2 alignment columns. Window support is the fraction of a study’s ASVs within the modal band; mapping quality is CONVERGED/UNSTABLE. Sample counts (n) are per study after cleaning (minimum depth 10,000 reads); the two withheld studies were excluded before merging (mapping instability or confirmed ASV degradation). ASVs, ASVs mapped to the reference frame; n, retained samples; Consensus and Roster, membership of the consensus overlap set and the inclusion roster.

| ***Study*** | ***Primer*** | ***Modal window*** | ***ASVs*** | ***Support*** | ***Quality*** | ***n*** | ***Consensus*** | ***Roster*** |
|---|---|---|---|---|---|---|---|---|
| PRJEB22038 | 341F_805R | 6,428–23,454 | 814 | 1.00 | CONVERGED | 19 | Yes | Yes |
| PRJEB25224 | 515F-Y_926R | 13,862–27,666 | 1,407 | 0.99 | CONVERGED | 102 | Yes | No |
| PRJEB27168 | 341F_805R | 6,428–23,444 | 103 | 1.00 | CONVERGED | 5 | Yes | Yes |
| PRJEB36282 | 515F-Y_926R | 13,862–27,684 | 3,273 | 1.00 | CONVERGED | 59 | Yes | Yes |
| PRJEB42499 | 515F_806R | 13,862–23,567 | 2,782 | 1.00 | CONVERGED | 121 | Yes | Yes |
| PRJEB50983 | 515F-Y_926R | 13,862–27,728 | 5,145 | 1.00 | CONVERGED | 321 | Yes | Yes |
| PRJEB52328 | 515F-Y_926R | 13,862–27,859 | 4,988 | 0.99 | CONVERGED | 56 | Yes | Yes |
| PRJEB90443 | 515F-Y_926R | 13,862–27,675 | 1,191 | 1.00 | CONVERGED | 15 | Yes | Yes |
| PRJNA263621 | 515F_806R | 6,238–37,980 | 3,104 | 0.24 | UNSTABLE | 23 | No | No |
| PRJNA664813 | 341F_805R | 6,428–23,448 | 623 | 1.00 | CONVERGED | 18 | Yes | Yes |
| PRJNA734169 | 515F_806R | 13,862–23,444 | 665 | 1.00 | CONVERGED | 31 | Yes | Yes |
| PRJNA780024 | 341F_805R | 6,428–23,449 | 363 | 1.00 | CONVERGED | 10 | Yes | Yes |
| PRJNA903199 | 515F_806R | 13,862–23,479 | 944 | 1.00 | CONVERGED | 27 | Yes | Yes |

The modal windows separated cleanly by primers. The window coordinates determined for each primer set are SILVA 138.2 alignment columns onto which every study is mapped (“Reference coordinate mapping” section) so windows are directly comparable between studies. The 341F–805R studies mapped to ≈6,428–23,450, the 515F(-Y)–926R studies to ≈13,862–27,700, and the 515F–806R studies to ≈13,862–23,500 (Table 3; Figure 7a). Twelve of the thirteen studies converged tightly on a single window (window-support 0.99–1.00; modal start and end median deviation = 0 alignment columns) and were classified as CONVERGED. One study, PRJNA263621, failed to converge (window support 0.24; start and end median deviation 892 and 2,473 columns) and was flagged UNSTABLE and excluded from consensus determination; its ASVs did not converge on one window, the pattern expected when a deposited dataset contains amplicons from more than one primer scheme (see Limitations).

The twelve CONVERGED studies formed a single, fully connected overlap set, so no boundary constrictor study required removal. Their intersection defined a consensus window of SILVA columns 13,862–23,444, 9,582 alignment columns wide, corresponding to a median trimmed read length of 249 bp (Figure 7b). The window is bounded below by the studies whose primers share the 515F forward site (the V4 and V4–V5 primer sets; modal start position 13,862) and above by the shortest V4 amplicons (minimum modal end position 23,444). The alignment overlap was largely representative of the V4 core common to all three primer sets.

Each study’s Bray–Curtis degradation curve was tested using three independent detectors that scanned for degradation within the trim length needed to reach the consensus region. These required trims were determined by primer set: 0–123 alignment columns for 515F-806R studies, which began on or near the trim boundary; 4,222–4,415 alignment columns for 515F(-Y)–926R studies; and 7,434–7,444 for the 341F-805R studies (Table 3). Across the 72 study × level tests (twelve studies × six taxonomic ranks), 31 (43%) showed no degradation signal at or before the required trim, including all six levels of the three 515F-806R studies. Thirty-eight carried a single detector warning before the required depth with no second corroboration. Three showed corroborated (CONFIRMED) degradation before the consensus was reached: PRJEB25224 at ASV level (1,700 columns against 4,222 required), PRJEB52328 at class level (1,550 against 4,415) and PRJNA780024 at phylum level (2,450 against 7,439). At ASV level specifically, four of twelve studies showed no degradation signal before their required trim, seven carried an uncorroborated warning and one (PRJEB25224) degraded before the consensus (Figure 8).

**Figure 8.**
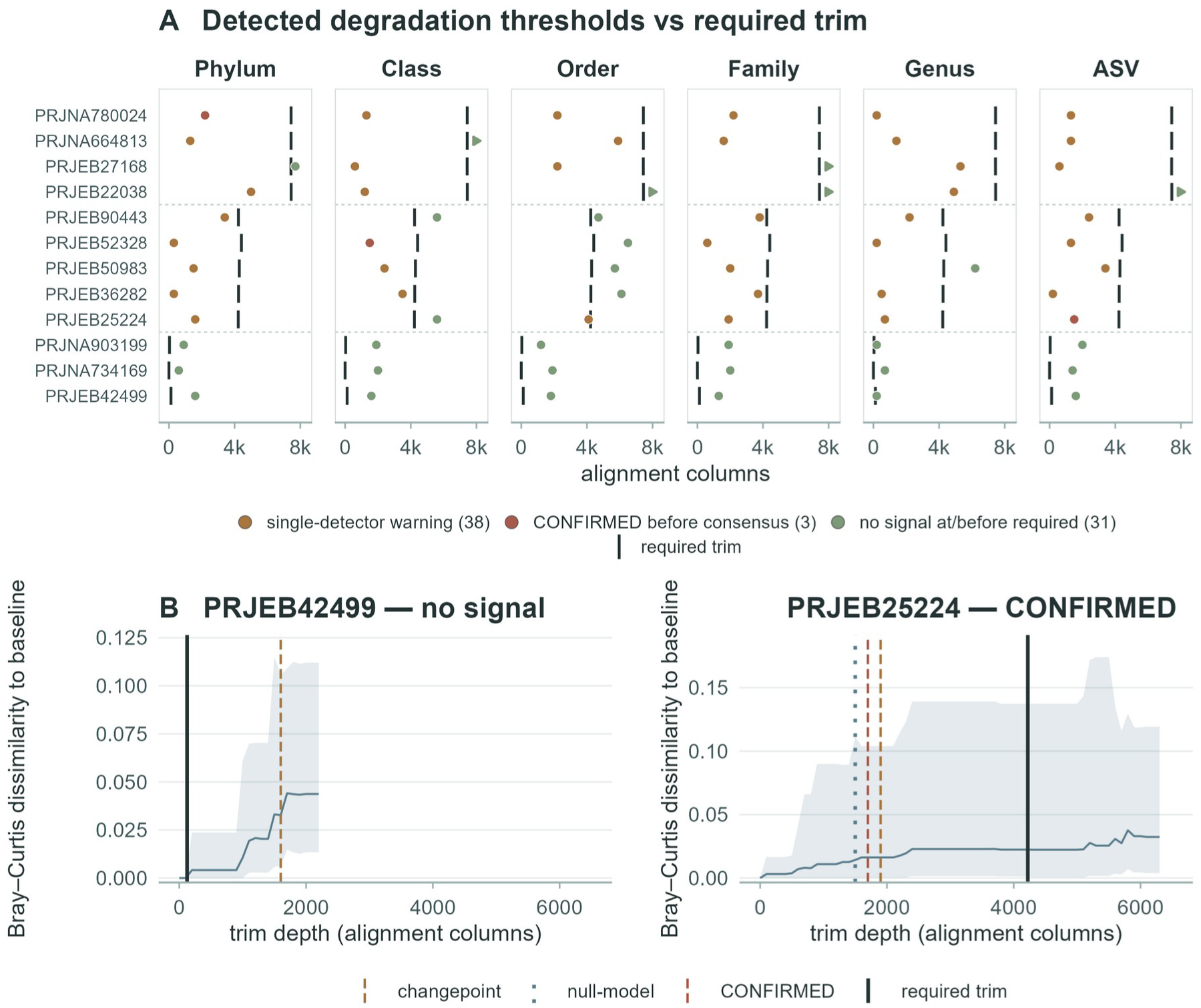
Consensus degradation verdicts for the twelve eligible studies. (A) Earliest detected degradation threshold for each of the 72 study × level tests (points, coloured by verdict class) against the required trim to the consensus window (vertical bars). Detections lying beyond the plotted range are drawn as arrows at the panel margin, coloured by verdict class; all lie far beyond the required trim. Studies are grouped by primer set (dotted rules). (B) Exemplar ASV-level degradation curves (mean simulated Bray–Curtis dissimilarity to the untrimmed baseline, with 95% band) for a passing study (PRJEB42499, detection far beyond the 123-column requirement) and the confirmed failure (PRJEB25224, CONFIRMED at 1,700 columns against 4,222 required). Vertical lines mark the detector thresholds (dashed, changepoint; dotted, null-model), the consensus CONFIRMED threshold (dashed) and the required trim (solid), identified in the key.

The three detectors behaved as designed (Figure 8; Supplementary Table S4; Supplementary Data S1). The changepoint detector, which locates the strongest structural break in every curve, proposed a break in all 72 tests (median location by rank 1,550–5,200 columns). The simulation null detection, which fires only where observed changes exceed the null, corroborated a break in 17 of 72 tests (median 1,500 columns). The empirical distance cutoff was never triggered among the eligible studies. Requiring agreement between at least two detectors therefore restricted the confirmed degradation to the three cases above, with single uncorroborated breaks recorded as warnings. PRJNA263621, the single UNSTABLE study was flagged during convergence testing (Table 3) and excluded from consensus determination.

### Harmonisation yields a cross-comparable MetaASV dataset

After the Phase 1 pause, eleven studies were curated into the inclusion roster. PRJEB25224 was withheld for corroborated pre-consensus degradation at ASV level; PRJNA263621 was excluded on its mapping instability. PRJEB52328 and PRJNA780024 were retained at ASV level as their corroborated shifts were confined to class and phylum respectively with all other levels clean or uncorroborated. All eleven studies trimmed successfully to the 9,582-column aligned window, with no sequences removed by the minimum length filter.

The eleven studies contributed 20,891 trimmed ASVs, which deduplicated at 100% identity into 12,667 MetaASVs across 682 samples (`combined_feature_table.tsv`). Retained studies contributed between 5 and 321 samples each (Table 3). Relative to the raw union of 108,165 study-specific ASVs held by the eleven studies before quality control, harmonisation produced a single coordinate-comparable feature table an order of magnitude smaller. Of the 3,511 MetaASVs built from more than one ASV, 1,061 had contributors with discordant taxonomy; the length term of the composite score changed the donor relative to a confidence-only choice in 112 of them (0.9% of all MetaASVs), at a median confidence cost of 0.4 points (maximum 6.6). Cross-study comparability increased accordingly: 21.1% of MetaASVs were observed in two or more studies (the remaining 78.9% were confined to a single study) and 2,004 MetaASVs were shared between studies from different primer sets (mean between-study Jaccard 0.047 within and 0.025 across primer sets). At aggregated ranks, the mean pairwise Jaccard similarity was 0.207 at genus and 0.335 at family, each meeting the minimum below baseline criteria (Table 4).

**Table 4.** Signal-preservation validation of the harmonised dataset at three taxonomic levels. Values are quoted from `validation_summary.csv`; criteria are the pipeline’s fixed pass thresholds. ICC, intraclass correlation; ΔRZ, post-minus pre-standardisation PERMANOVA RZ on `StudyID` (should not increase).

| *Metric* | *ASV* | *Genus* | *Family* | *Criterion* |
|---|---|---|---|---|
| *Beta-diversity (Mantel r)* | 0.790 | 1.000 | 1.000 | > 0.85 |
| *Procrustes m²* | 0.522 | <0.001 | <0.001 | < 0.10 |
| *PROTEST correlation* | 0.692 | 1.000 | 1.000 | > 0.95 |
| *PERMANOVA ΔR² (StudyID)* | −0.039 | −0.001 | <0.001 | ≤ 0.05 |
| *Dispersion change* | 1.9% | 0.1% | <0.1% | < 10% |
| *Alpha ICC (Shannon)* | 0.927 | 0.999 | 1.000 | > 0.90 |
| *Rank abundance (Spearman ρ)* | 0.943 | 0.995 | 0.998 | > 0.90 |
| *Abundance concordance (ρ)* | n/a | 0.974 | 0.988 | > 0.90 |
| *Network vertex/ edge change* | −0.6% / −15.9% | −1.2% / −7.9% | +1.3% / +1.6% | <20% |
| *Hub node retention* | 0.23 | 0.89 | 1.00 | ≥0.80 |
| *Cross-study sharing / Jaccard* | 0.211 | 0.207 | 0.335 | ≥ baseline |

### Community structure is conserved across taxonomic levels

At the sequence level, all 20,891 rostered ASVs across the eleven studies trimmed successfully to the consensus window. The pooled length distribution narrowed from a median of 372 bp (range 230–430) to 249 bp with the coefficient of variation falling from 0.147 to 0.019, and nucleotide composition was unchanged (median GC content 0.515 before and 0.514 after trimming).

Harmonisation did not introduce between-study structure and preserved community structure, with fidelity increasing from ASV to family (Table 4; Figure 9; Supplementary Table S3). Under PERMANOVA on `StudyID`, the variance explained by study did not increase appreciably after standardisation at any level (ΔRZ = −0.039 at ASV, −0.001 at genus and +0.0002 at family; `adonis2`, 999 permutations), indicating that trimming removed rather than added apparent batch structure.

**Figure 9.**
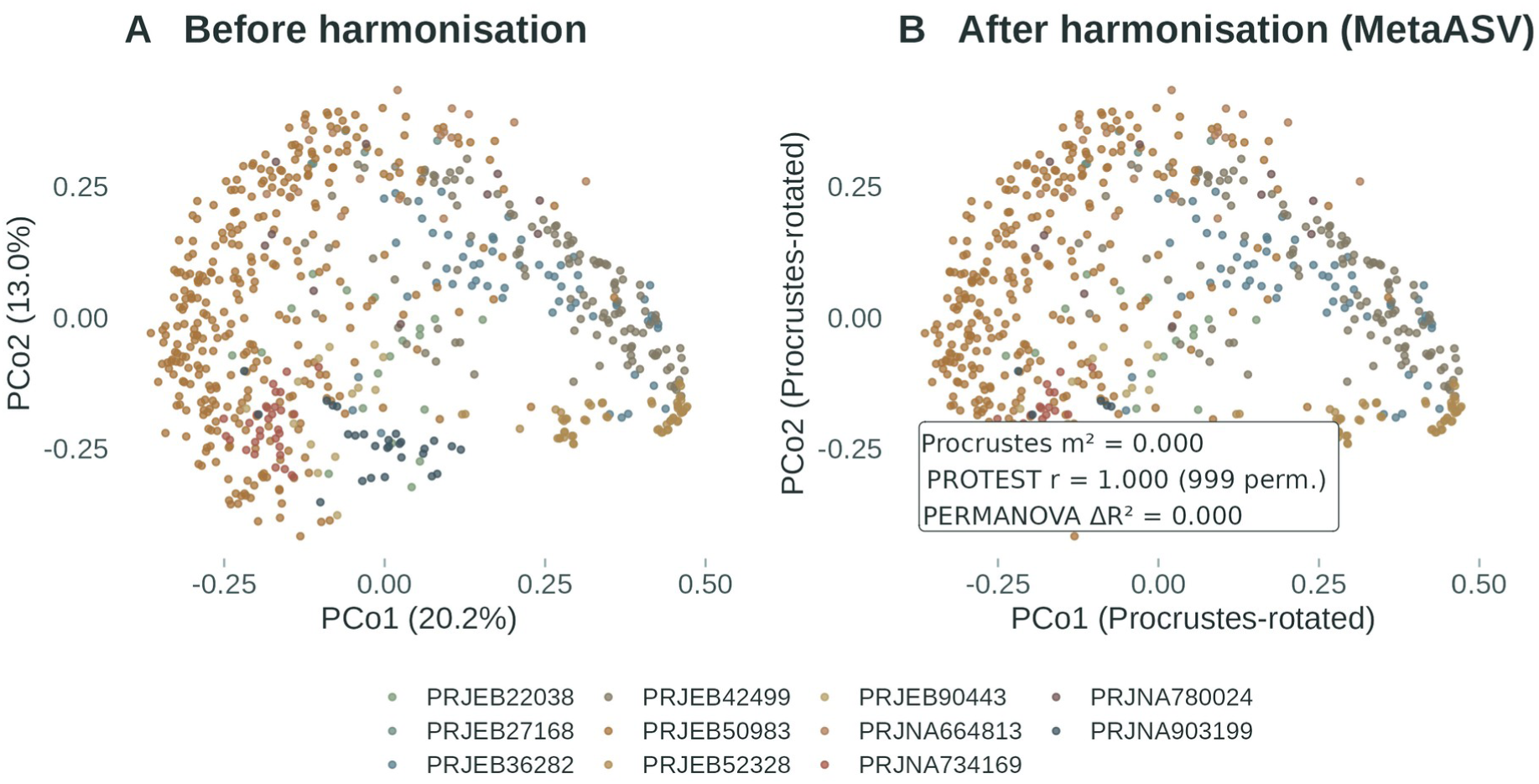
Community structure is preserved through harmonisation. Family-level principal coordinates analysis (Bray–Curtis dissimilarity on untransformed counts) of the 682 harmonised samples, before (A) and after (B) trimming to the consensus window and collapse to MetaASVs, coloured by study. The post-harmonisation ordination (B) is shown after Procrustes rotation onto the pre-harmonisation ordination, so panel orientations and scales are directly comparable. Procrustes and PROTEST statistics (999 permutations) compare the two ordinations over the first three principal coordinate axes; ΔRZ is the change in the variance explained by study (PERMANOVA on `StudyID`). Both panels show a weak arch (PCo2 depends quadratically on PCo1, RZ = 0.09); it is identical before and after harmonisation and does not affect the Procrustes comparison. The corresponding ASV-and genus-level ordinations are shown in Supplementary Figure S2.

Beta diversity concordance between the pre- and post-standardisation distance matrices rose with higher taxonomic levels. Mantel correlation rose from r = 0.790 at ASV level to effective unity at genus and family (both ≥ 0.9997; 999 permutations). Ordination geometry followed the same trend, with Procrustes mZ improving from 0.522 at ASV level to below 0.001 at genus and family (PROTEST correlation 0.692, 1.000 and 1.000; Figure 9; Supplementary Figure S2). Within study dispersion remained minimally affected with the relative change in mean distance-to-centroid ≤ 1.9% across all taxonomic levels (Table 4; Supplementary Figure S3). Study ASVs are study-specific features, so every between-study sample pair had a Bray–Curtis dissimilarity of exactly 1 before harmonisation (all 169,050 between-study pairs, within and across primer sets); the ASV-level Mantel and Procrustes statistics therefore compare within-study geometry only. After harmonisation, 89.8% of between-study pairs within a primer set and 58.7% across primer sets shared at least one MetaASV (mean dissimilarity 0.932 and 0.949), whereas family-level dissimilarities were unchanged by harmonisation.

Alpha diversity agreement was high for every component measured (Table 4; Supplementary Figure S1). Abundance weighted indices were in agreement between the original and harmonised datasets across all levels (intraclass correlation 0.93–1.00 for Shannon and 0.93–1.00 for Simpson). Richness agreement was similarly high (observed richness 0.97 at ASV level and ≥ 0.999 at genus and family; Chao1 and ACE 0.97–1.00). Rank abundance structure showed increased preservation at higher taxonomic levels (Spearman ρ = 0.94 at ASV level, 0.995 at genus and 0.998 at family) and per taxon abundance concordance at the higher taxonomic levels was similarly strong (mean ρ = 0.97 at genus and 0.99 at family; Supplementary Figure S4).

Co-occurrence network architecture was equally stable. SPIEC-EASI networks inferred from prevalence filtered tables before and after harmonisation differed by less than 2% in vertex count at every level (333 to 331 vertices at MetaASV level; 86 to 85 at genus; 78 to 79 at family), with edge counts stable at family (122 to 124) and genus (139 to 128) but reduced by 16% at MetaASV level (1,734 to 1,459). The most connected taxa were retained at genus (8 of 9 hub nodes) and family (8 of 8), whereas at MetaASV level the network was internally reorganised: all mapped hub features remained present in the post harmonisation network, but only 5 of 22 remained among the most connected, and vertex degree was only weakly correlated between the two networks (Spearman ρ = 0.28 across the 219 features present in both).

### Performance

The pipeline ran in two phases on the JASMIN LOTUS2 SLURM cluster in published container images (Supplementary Table S2). Phase 1 (coordinate mapping, simulation and verdict generation) for the thirteen mapped studies completed in 3 h 39 m wall-clock and 65 CPU h, of which the 1,200 simulation replicates (100 per converged study; single-core, mean 3.3 min, maximum 12.5 min, under 4 GB) accounted for 97%. The memory-limiting step was the per-study DECIPHER profile alignment (peak 28.9 GB; at most 7 min per study); all other Phase 1 stages completed within minutes. Phase 2 (trimming, MetaASV merging and validation of the eleven rostered studies) completed in 1 h 30 m, with validation the limiting step (1 h 17 m, peak 4.2 GB), dominated by network inference and permutation testing. The complete `final_outputs` tree archives to approximately 0.3 GB compressed. Full per-task traces and the execution timeline are provided in Supplementary Table S2.

## Discussion

SeCAT harmonises 16S rRNA metabarcoding datasets generated with different primer sets into a single table of cross-comparable MetaASVs, by mapping every study onto a common reference alignment and validating, by simulation, a trim to their shared consensus region.

The framework is general and applies to any harmonisable set of studies whose primer windows overlap on a suitable full-length reference. The validation stage presented is not a test of the functioning of the tool, but a per-dataset comparability test and for each new synthesis quantifies any drift of the pooled table compared to the original communities, so that results from the combined data can be interpreted against the source studies rather than in isolation. Even where a study drifts substantially from its original community, it may still be included in a meta-analysis. Comparisons within the harmonised table remain valid, and SeCAT documents (where such drift has occurred) can be weighed explicitly.

### Interpreting the validation: what is preserved and where the data drift

The validation separates two components of community structure that respond differently to harmonisation. Abundance weighted structure assesses which taxa dominate the community and how evenly abundance is distributed among them (Hill 1973; Jost 2006). These metrics are preserved at every rank, including ASV resolution. The moderate ordination congruence at sequence level follows from redefining the units rather than from any loss of fidelity: study-specific ASVs cannot be shared between studies, so between-study similarity at sequence level exists only after harmonisation. The dominant component of each community is therefore conserved through harmonisation, consistent with prior demonstrations that abundance-weighted structure is insensitive to the delimitation of sequence units (Glassman & Martiny, 2018; Nearing *et al*. 2018). Co-occurrence architecture followed the same trends. Network size and connectivity were stable at every level, and hub identity was conserved at genus and family but reorganised at exact sequence level.

Richness was also recovered, although it is the component most exposed to harmonisation, because trimming to the consensus window redefines the units over which richness is counted and any departure from the original values therefore reflects the procedure itself rather than degradation of the underlying data. ASVs are resolved over the full amplicon, so two ASVs may differ only within the flanking regions that trimming removes. These variants become identical over the consensus regions and are therefore combined into a single MetaASV (Johnson *et al*. 2019; Bukin *et al*. 2019). Across primer sets the same consolidation is the intended outcome: reads from one organism amplified over different windows unify into one MetaASV. Mergers conserve reads (constituent abundances are summed) while reducing the number of units. Abundance-weighted indices are dominated by abundant taxa and are insensitive to the rare tail, so summing counts within merged units leaves them largely unchanged. Richness weights every unit equally so each merger registers as a loss, and estimators that extrapolate from singleton counts (Chao 1984; Chao & Lee 1992; Chiu & Chao 2016) are the most sensitive to the reorganisation of the rare tail, which is itself susceptible to residual sequencing artefacts (Schloss *et al*. 2011). Agreement between pre- and post-harmonisation richness improves when MetaASVs are aggregated to genus or family because variants that merge during harmonisation typically derive from the same taxon. The count of the exact sequences falls, but the count of the named taxa that they represent remains largely unchanged. The accompanying decrease in pairwise taxon overlap between studies at these ranks arises where taxa represented by a few reads fall below detection after trimming and merging.

These results are the verdict of this validation, not fixed properties of the method, and they guide use of the harmonised output. Exact sequence comparison across studies, previously unreliable due to differing primer sets and lack of feature sharing (Klindworth *et al*. 2013; Callahan *et al*. 2017), is supported for compositional, abundance-weighted and richness-based inference (Gloor et al. 2017) where validation shows preservation of original metrics; where any metric falls short of its criterion, the harmonised table should be treated as its own baseline rather than as a proxy for the per-study values it replaces. Aggregation to genus or family remains an analytical option rather than a prerequisite. A different combination of studies may yield a different verdict; the validation report generated for each synthesis defines which comparisons its dataset supports.

### Position relative to existing approaches

Existing approaches to cross-study comparison each leave the positional problem unaddressed. Statistical batch-effect correction (MMUPHin, ConQuR and PLSDA-batch; Ma et al. 2022; Ling et al. 2022; Wang & Lê Cao, 2023) is applied after a feature table has been assembled, to remove study-associated variation while preserving biological signal. It suits syntheses in which every study used the same primer pair, but it presupposes a shared feature space and has no mechanism for creating one. Closed-reference OTU picking (Rideout et al. 2014) does create shared identifiers by assigning reads to reference centroids, which is why it has been the usual route to combining multi-region datasets; the cost is that sequences absent from the reference are discarded (Lloyd et al. 2018) and that the shared label certifies taxonomy rather than positional overlap, so two reads can carry one identifier while spanning non-overlapping windows (Pollock et al. 2018). Phylogenetic placement (MaLiAmPi; Minot et al. 2023) is the closest alternative in intent: ASVs from each study are placed with EPA-ng onto a reference phylogeny of full-length 16S rRNA alleles, and the resulting phylotypes are comparable across primer regions because the tree, not the amplicon, supplies the common frame. Its features are defined by position on the tree: robust to region choice, but not exact sequences, and bounded in resolution by the reference package. SeCAT differs from all three in establishing positional overlap from the sequences themselves before any merging, and in returning sequence-defined MetaASVs that import directly into standard workflows (phyloseq; McMurdie & Holmes 2013) and remain exact, portable sequences. A MetaASV table remains a valid input for subsequent batch correction or tree-based aggregation, so SeCAT complements rather than replaces these tools. Its distinctive contribution is the explicit, simulation-referenced test that trimming preserves community composition and structure; the paired pre- and post-harmonisation validation and empirical null give a quantified fidelity statement that none of the alternatives provides.

### Practical guidance for users

Three recommendations follow for the users. First, the diagnostic pause and roster should be used as intended. The window-support gate and per-study verdicts make each inclusion decision explicit and auditable; marginal cases should be resolved from the per-study reports, and taxonomy should be assigned to every input study with one classifier and reference database, for the reason given in the Methods. Second, the scope of legitimate cross-study comparison should be taken from the validation report generated for each synthesis rather than assumed: comparisons with the original studies’ conclusions should be restricted to the components the report shows to be preserved, or expected drift documented from the taxonomic-impact outputs. Third, for co-occurrence network analysis, low prevalence MetaASVs should be filtered before inference, because correlations involving features observed in only a few samples cannot be estimated reliably (Friedman & Alm 2012; Kurtz *et al*. 2015). Because many MetaASVs are recovered from only a subset of studies, the pooled table is also enriched for features whose absences reflect study membership rather than ecology, which network inference can misread as co-occurrence structure (Berry & Widder 2014; Carr et al. 2019). Network inference is also better supported at aggregated ranks, where the present validation found conserved hub structure.

### Limitations

SeCAT is designed to establish and validate positional comparability. Fidelity is bounded by the quality and the coverage of the reference database the package is run on. Taxa poorly represented in the reference alignment map less reliably, and the consensus window can be located only as precisely as the reference allows (Quast et al. 2013). Separately, the window-support gate removes studies whose ASVs do not converge on a single window, as when a deposited dataset contains amplicons from more than one primer scheme, at the cost of otherwise usable data; it is retained because such a study has no single required trim, and the remedy is to split the dataset by primer scheme in the manifest. SeCAT harmonises features but does not recall them, so it inherits the denoising decisions of each source study (Nearing et al. 2018), and the roster step, which improves auditability, remains a subjective element of the workflow. The recoverable window and achievable sharing rate depend on the primer composition of each synthesis, and a formal head-to-head benchmark against batch correction and closed-reference approaches has yet to be performed; such a benchmark is feasible only within a primer set, because across primer sets no shared feature space exists until harmonisation has created one. Zeros introduced by the full outer join also deserve caution: a MetaASV recorded as absent from a given study may be undetected rather than truly absent, a false negative driven by primer bias or sequencing depth, so cross-study absences should be read as non-detection. Where absence matters, such false negatives can be partly accommodated downstream, for example with occupancy models that estimate detection probability explicitly (MacKenzie et al. 2002; Ficetola et al. 2015).

The wider opportunity is set by the scale of what already exists. Coordinated programmes such as the Earth Microbiome Project showed what unified primers and protocols make possible (Gilbert, Jansson & Knight, 2014; Thompson et al. 2017), but most archived datasets were never coordinated, and syntheses built on them have worked around primer fragmentation rather than through it: aggregating to coarse ranks (Ramirez et al. 2018), restricting synthesis to studies sharing a primer set, or treating methodological heterogeneity as a nuisance term (Fediajevaite et al. 2021). By establishing positional comparability first and quantifying its cost per study, SeCAT converts that fragmented archive into usable sequence-level material: harmonised MetaASV tables can place historical baselines beneath present-day surveys, widen the reach of trait and biogeography analyses (Thompson et al. 2017; Ma et al. 2022), and let monitoring programmes that changed primers mid-series recover continuity. The same fragmentation affects human-microbiome archives, where surveys of one body site have targeted V1–V3 and V3–V5 (Human Microbiome Project Consortium, 2012) as well as V4 (Thompson et al. 2017), so sequence-level harmonisation would also widen the evidence base for meta-analyses relating microbiome composition to health outcomes. Because every inclusion decision is validated and documented, such syntheses also inherit an explicit statement of what the merged data can and cannot support.

Looking forward, the consensus-window logic generalises to any marker gene with a full-length reference alignment; extending SeCAT beyond 16S rRNA as further primer sets enter the archive is a natural next step.

## Acknowledgements

This work used JASMIN, the UK collaborative data analysis facility. The authors acknowledge the use of the High Performance Computing Facility (Ceres) and its associated support services at the University of Essex in the completion of this work. We thank the µSudAqua network (https://microsudaqua.netlify.app/en/) for creating a space where colleagues from across South America connect and exchange ideas. The original concept for this pipeline emerged from one such exchange between P.H., E.P. and S.M. We thank the authors of the thirteen source studies for making their sequence data publicly available.

## Funding

This work was supported by internal doctoral funding from the University of Derby. No additional external funding was received for the development of this software.

## Conflict of interest

The authors declare no conflict of interest.

## Data Accessibility and Benefit-Sharing

SeCAT v5.0.6 is open source at github.com/DETerrey/SeCAT under the MIT licence and is archived at Zenodo (v5.0.6 release DOI: https://doi.org/10.5281/zenodo.22286369; concept DOI resolving to the current release: https://doi.org/10.5281/zenodo.22113431). It requires Nextflow (DSL2) and a Singularity/Apptainer or Docker runtime; R and all package dependencies are supplied in the published container image (ghcr.io/deterrey/secat). The reference database is SILVA 138.2 SSURef NR99 (Quast *et al*. 2013). The study manifests, params.yaml, selection_roster.txt and the harmonised MetaASV outputs are provided in Zenodo (https://doi.org/10.5281/zenodo.22871720), and the source 16S datasets are available from their original accessions (Supplementary Table S1). Study-associated sample metadata are included in the Zenodo outputs deposit and accompany the original sequence archives.

This study analysed publicly archived sequence data generated by other research groups; no new biological samples were collected. Benefits from this research accrue from the sharing of our data, results and software on public repositories, and all outputs are released under open licences to the wider scientific community.

## Author contributions

David Terrey: conceptualisation, methodology, software, formal analysis, investigation, visualisation, writing - original draft. Paula Huber: conceptualisation (original concept), data curation, writing - review and editing. Emiliano Pereira: conceptualisation (original concept), data curation, writing - review and editing. Maria C. Pelayo: data curation. Robert M. W. Ferguson: resources, writing - review and editing. Alex J. Dumbrell: resources, writing - review and editing. Michael Sweet: supervision, writing - review and editing. Alessia Bani: supervision, writing - review and editing. Sebastian Metz: conceptualisation (original concept), supervision, data curation, methodology, writing - review and editing.

## Notes

https://github.com/DETerrey/SeCAT

https://doi.org/10.5281/zenodo.22871720

### Competing Interest Statement

The authors have declared no competing interest.

## References

Abellan-Schneyder, I., Matchado, M. S., Reitmeier, S., Sommer, A., Sewald, Z., Baumbach, J., List, M., & Neuhaus, K. (2021). Primer, pipelines, parameters: Issues in 16S rRNA gene sequencing. mSphere, 6(1), e01202–20. 10.1128/mSphere.01202-20

Aird, D., Ross, M. G., Chen, W.-S., Danielsson, M., Fennell, T., Russ, C., Jaffe, D. B., Nusbaum, C., & Gnirke, A. (2011). Analyzing and minimizing PCR amplification bias in Illumina sequencing libraries. Genome Biology, 12(2), R18. 10.1186/gb-2011-12-2-r18

Amann, R. I., Binder, B. J., Olson, R. J., Chisholm, S. W., Devereux, R., & Stahl, D. A. (1990). Combination of 16S rRNA-targeted oligonucleotide probes with flow cytometry for analyzing mixed microbial populations. Applied and Environmental Microbiology, 56(6), 1919–1925. 10.1128/aem.56.6.1919-1925.1990

Amir, A., McDonald, D., Navas-Molina, J. A., Kopylova, E., Morton, J. T., Xu, Z. Z., Kightley, E. P., Thompson, L. R., Hyde, E. R., Gonzalez, A., & Knight, R. (2017). Deblur rapidly resolves single-nucleotide community sequence patterns. mSystems, 2(2), e00191–16. 10.1128/mSystems.00191-16

Anderson, M. J. (2001). A new method for non-parametric multivariate analysis of variance. Austral Ecology, 26(1), 32–46. 10.1111/j.1442-9993.2001.01070.pp.x

Anderson, M. J. (2006). Distance-based tests for homogeneity of multivariate dispersions. Biometrics, 62(1), 245–253. 10.1111/j.1541-0420.2005.00440.x

Angly, F. E., Willner, D., Rohwer, F., Hugenholtz, P., & Tyson, G. W. (2012). Grinder: A versatile amplicon and shotgun sequence simulator. Nucleic Acids Research, 40(12), e94. 10.1093/nar/gks251

Berry, D., & Widder, S. (2014). Deciphering microbial interactions and detecting keystone species with co-occurrence networks. Frontiers in Microbiology, 5, 219. 10.3389/fmicb.2014.00219

Bray, J. R., & Curtis, J. T. (1957). An ordination of the upland forest communities of southern Wisconsin. Ecological Monographs, 27(4), 325–349. 10.2307/1942268

Bukin, Y. S., Galachyants, Y. P., Morozov, I. V., Bukin, S. V., Zakharenko, A. S., & Zemskaya, T. I. (2019). The effect of 16S rRNA region choice on bacterial community metabarcoding results. Scientific Data, 6(1), 190007. 10.1038/sdata.2019.7

Callahan, B. J., McMurdie, P. J., & Holmes, S. P. (2017). Exact sequence variants should replace operational taxonomic units in marker-gene data analysis. The ISME Journal, 11(12), 2639–2643. 10.1038/ismej.2017.119

Callahan, B. J., McMurdie, P. J., Rosen, M. J., Han, A. W., Johnson, A. J. A., & Holmes, S. P. (2016). DADA2: High-resolution sample inference from Illumina amplicon data. Nature Methods, 13(7), 581–583. 10.1038/nmeth.3869

Camacho, C., Coulouris, G., Avagyan, V., Ma, N., Papadopoulos, J., Bealer, K., & Madden, T. L. (2009). BLAST+: architecture and applications. BMC Bioinformatics, 10, 421. 10.1186/1471-2105-10-421

Caporaso, J. G., Lauber, C. L., Walters, W. A., Berg-Lyons, D., Lozupone, C. A., Turnbaugh, P. J., Fierer, N., & Knight, R. (2011). Global patterns of 16S rRNA diversity at a depth of millions of sequences per sample. Proceedings of the National Academy of Sciences, 108(supplement_1), 4516–4522. 10.1073/pnas.1000080107

Carr, A., Diener, C., Baliga, N. S., & Gibbons, S. M. (2019). Use and abuse of correlation analyses in microbial ecology. The ISME Journal, 13(11), 2647–2655. 10.1038/s41396-019-0459-z

Chao, A. (1984). Nonparametric estimation of the number of classes in a population. Scandinavian Journal of Statistics, 11(4), 265–270.

Chao, A., & Lee, S.-M. (1992). Estimating the number of classes via sample coverage. Journal of the American Statistical Association, 87(417), 210–217. 10.1080/01621459.1992.10475194

Chiu, C.-H., & Chao, A. (2016). Estimating and comparing microbial diversity in the presence of sequencing errors. PeerJ, 4, e1634. 10.7717/peerj.1634

Csárdi, G., & Nepusz, T. (2006). The igraph software package for complex network research. InterJournal, Complex Systems, 1695.

Curtis, T. P., Sloan, W. T., & Scannell, J. W. (2002). Estimating prokaryotic diversity and its limits. Proceedings of the National Academy of Sciences, 99(16), 10494–10499. 10.1073/pnas.142680199

Di Tommaso, P., Chatzou, M., Floden, E. W., Prieto Barja, P., Palumbo, E., & Notredame, C. (2017). Nextflow enables reproducible computational workflows. Nature Biotechnology, 35(4), 316–319. 10.1038/nbt.3820

Duvallet, C., Gibbons, S. M., Gurry, T., Irizarry, R. A., & Alm, E. J. (2017). Meta-analysis of gut microbiome studies identifies disease-specific and shared responses. Nature Communications, 8(1), 1784. 10.1038/s41467-017-01973-8

Fediajevaite, J., Priestley, V., Arnold, R., & Savolainen, V. (2021). Meta-analysis shows that environmental DNA outperforms traditional surveys, but warrants better reporting standards. Ecology and Evolution, 11(9), 4803–4815. 10.1002/ece3.7382

Ficetola, G. F., Pansu, J., Bonin, A., Coissac, E., Giguet-Covex, C., De Barba, M., Gielly, L., Lopes, C. M., Boyer, F., Pompanon, F., Rayé, G., & Taberlet, P. (2015). Replication levels, false presences and the estimation of the presence/absence from eDNA metabarcoding data. Molecular Ecology Resources, 15(3), 543–556. 10.1111/1755-0998.12338

Friedman, J., & Alm, E. J. (2012). Inferring correlation networks from genomic survey data. PLoS Computational Biology, 8(9), e1002687. 10.1371/journal.pcbi.1002687

Gilbert, J. A., Jansson, J. K., & Knight, R. (2014). The Earth Microbiome project: Successes and aspirations. BMC Biology, 12(1), 69. 10.1186/s12915-014-0069-1

Gilbert, J. A., Peixoto, R. S., Scholz, A. H., Dominguez Bello, M. G., Korsten, L., Berg, G., Singh, B., Boetius, A., Wang, F., Greening, C., Wrighton, K., Bordenstein, S., Jansson, J. K., Lennon, J. T., Souza, V., Thomas, T., Cowan, D., Crowther, T. W., Nguyen, N., … Redford, K. (2025). Launching the IUCN Microbial Conservation Specialist Group as a global safeguard for microbial biodiversity. Nature Microbiology, 10(10), 2359–2360. 10.1038/s41564-025-02113-5

Glassman, S. I., & Martiny, J. B. H. (2018). Broadscale ecological patterns are robust to use of exact sequence variants versus operational taxonomic units. mSphere, 3(4), e00148–18. 10.1128/mSphere.00148-18

Gloor, G. B., Macklaim, J. M., Pawlowsky-Glahn, V., & Egozcue, J. J. (2017). Microbiome datasets are compositional: And this is not optional. Frontiers in Microbiology, 8, 2224. 10.3389/fmicb.2017.02224

Gower, J. C. (1975). Generalized Procrustes analysis. Psychometrika, 40(1), 33–51. 10.1007/BF02291478

Haas, B. J., Gevers, D., Earl, A. M., Feldgarden, M., Ward, D. V., Giannoukos, G., Ciulla, D., Tabbaa, D., Highlander, S. K., Sodergren, E., Methé, B., DeSantis, T. Z., Petrosino, J. F., Knight, R., & Birren, B. W. (2011). Chimeric 16S rRNA sequence formation and detection in Sanger and 454-pyrosequenced PCR amplicons. Genome Research, 21(3), 494–504. 10.1101/gr.112730.110

Herlemann, D. P. R., Labrenz, M., Jürgens, K., Bertilsson, S., Waniek, J. J., & Andersson, A. F. (2011). Transitions in bacterial communities along the 2000 km salinity gradient of the Baltic Sea. The ISME Journal, 5(10), 1571–1579. 10.1038/ismej.2011.41

Hill, M. O. (1973). Diversity and evenness: A unifying notation and its consequences. Ecology, 54(2), 427–432. 10.2307/1934352

Hong, S., Bunge, J., Leslin, C., Jeon, S., & Epstein, S. S. (2009). Polymerase chain reaction primers miss half of rRNA microbial diversity. The ISME Journal, 3(12), 1365–1373. 10.1038/ismej.2009.89

Human Microbiome Project Consortium. (2012). Structure, function and diversity of the healthy human microbiome. Nature, 486(7402), 207–214. 10.1038/nature11234

Janda, J. M., & Abbott, S. L. (2007). 16S rRNA gene sequencing for bacterial identification in the diagnostic laboratory: Pluses, perils, and pitfalls. Journal of Clinical Microbiology, 45(9), 2761–2764. 10.1128/JCM.01228-07

Johnson, J. S., Spakowicz, D. J., Hong, B.-Y., Petersen, L. M., Demkowicz, P., Chen, L., Leopold, S. R., Hanson, B. M., Agresta, H. O., Gerstein, M., Sodergren, E., & Weinstock, G. M. (2019). Evaluation of 16S rRNA gene sequencing for species and strain-level microbiome analysis. Nature Communications, 10(1), 5029. 10.1038/s41467-019-13036-1

Jost, L. (2006). Entropy and diversity. Oikos, 113(2), 363–375. 10.1111/j.2006.0030-1299.14714.x

Killick, R., & Eckley, I. A. (2014). changepoint: An R package for changepoint analysis. Journal of Statistical Software, 58(3), 1–19. 10.18637/jss.v058.i03

Killick, R., Fearnhead, P., & Eckley, I. A. (2012). Optimal detection of changepoints with a linear computational cost. Journal of the American Statistical Association, 107(500), 1590–1598. 10.1080/01621459.2012.737745

Klindworth, A., Pruesse, E., Schweer, T., Peplies, J., Quast, C., Horn, M., & Glöckner, F. O. (2013). Evaluation of general 16S ribosomal RNA gene PCR primers for classical and next-generation sequencing-based diversity studies. Nucleic Acids Research, 41(1), e1. 10.1093/nar/gks808

Koo, T. K., & Li, M. Y. (2016). A guideline of selecting and reporting intraclass correlation coefficients for reliability research. Journal of Chiropractic Medicine, 15(2), 155–163. 10.1016/j.jcm.2016.02.012

Kurtz, Z. D., Müller, C. L., Miraldi, E. R., Littman, D. R., Blaser, M. J., & Bonneau, R. A. (2015). Sparse and compositionally robust inference of microbial ecological networks. PLoS Computational Biology, 11(5), e1004226. 10.1371/journal.pcbi.1004226

Kurtzer, G. M., Sochat, V., & Bauer, M. W. (2017). Singularity: Scientific containers for mobility of compute. PLOS ONE, 12(5), e0177459. 10.1371/journal.pone.0177459

Lane, D. J. (1991). 16S/23S rRNA sequencing. In E. Stackebrandt & M. Goodfellow (Eds.), Nucleic acid techniques in bacterial systematics (pp. 115–175). John Wiley & Sons.

Laursen, M. F., Dalgaard, M. D., & Bahl, M. I. (2017). Genomic GC-content affects the accuracy of 16S rRNA gene sequencing based microbial profiling due to PCR bias. Frontiers in Microbiology, 8, 1934. 10.3389/fmicb.2017.01934

Lee, H. B., Jeong, D. H., Cho, B. C., & Park, J. S. (2023). Comparative analyses of eight primer sets commonly used to target the bacterial 16S rRNA gene for marine metabarcoding-based studies. Frontiers in Marine Science, 10, 1199116. 10.3389/fmars.2023.1199116

Li, Z., Chen, Y., Sun, Y., McArthur, K., Carnes, M., Liu, T., Mueller, N. T., Page, G. P., Rossman, L., Smirnova, E., White, J. D., Kress, A. M., & Debelius, J. W. (2025). In harmony? A scoping review of methods to combine multiple 16S amplicon data sets. bioRxiv. 10.1101/2025.02.11.637740

Ling, W., Lu, J., Zhao, N., Lulla, A., Plantinga, A. M., Fu, W., Zhang, A., Liu, H., Song, H., Li, Z., Chen, J., Randolph, T. W., Koay, W. L. A., White, J. R., Launer, L. J., Fodor, A. A., Meyer, K. A., & Wu, M. C. (2022). Batch effects removal for microbiome data via conditional quantile regression. Nature Communications, 13(1), 5418. 10.1038/s41467-022-33071-9

Liu, C., Cui, Y., Li, X., & Yao, M. (2021). microeco: An R package for data mining in microbial community ecology. FEMS Microbiology Ecology, 97(2), fiaa255. 10.1093/femsec/fiaa255

Lloyd, K. G., Steen, A. D., Ladau, J., Yin, J., & Crosby, L. (2018). Phylogenetically novel uncultured microbial cells dominate Earth microbiomes. mSystems, 3(5), e00055–18. 10.1128/mSystems.00055-18

López-Aladid, R., Fernández-Barat, L., Alcaraz-Serrano, V., Bueno-Freire, L., Vázquez, N., Pastor-Ibáñez, R., Palomeque, A., Oscanoa, P., & Torres, A. (2023). Determining the most accurate 16S rRNA hypervariable region for taxonomic identification from respiratory samples. Scientific Reports, 13(1), 3974. 10.1038/s41598-023-30764-z

Ma, S., Shungin, D., Mallick, H., Schirmer, M., Nguyen, L. H., Kolde, R., Franzosa, E., Vlamakis, H., Xavier, R., & Huttenhower, C. (2022). Population structure discovery in meta-analyzed microbial communities and inflammatory bowel disease using MMUPHin. Genome Biology, 23(1), 208. 10.1186/s13059-022-02753-4

MacKenzie, D. I., Nichols, J. D., Lachman, G. B., Droege, S., Royle, J. A., & Langtimm, C. A. (2002). Estimating site occupancy rates when detection probabilities are less than one. Ecology, 83(8), 2248–2255. 10.1890/0012-9658(2002)083[2248:ESORWD]2.0.CO;2

Mantel, N. (1967). The detection of disease clustering and a generalized regression approach. Cancer Research, 27(2), 209–220.

Martin, M. (2011). Cutadapt removes adapter sequences from high-throughput sequencing reads. EMBnet.journal, 17(1), 10–12. 10.14806/ej.17.1.200

Martinez-Porchas, M., Villalpando-Canchola, E., Ortiz Suarez, L. E., & Vargas-Albores, F. (2017). How conserved are the conserved 16S-rRNA regions? PeerJ, 5, e3036. 10.7717/peerj.3036

McArdle, B. H., & Anderson, M. J. (2001). Fitting multivariate models to community data: A comment on distance-based redundancy analysis. Ecology, 82(1), 290–297. 10.1890/0012-9658(2001)082[0290:FMMTCD]2.0.CO;2

McMurdie, P. J., & Holmes, S. (2013). phyloseq: An R package for reproducible interactive analysis and graphics of microbiome census data. PLoS ONE, 8(4), e61217. 10.1371/journal.pone.0061217

Metz, S., Huber, P., Mateus-Barros, E., Junger, P. C., de Melo, M., Bagatini, I. L., Izaguirre, I., dos Reis, M. C., Llames, M. E., Accattatis, V., Quiroga, M. V., Devercelli, M., Schiaffino, M. R., Niño-García, J. P., Bastidas Navarro, M., Modenutti, B., Vieira, H., Saraceno, M., Sabio y García, C. A., Pereira, E., González-Revello, A., Piccini, C., Unrein, F., Alonso, C., & Sarmento, H. (2022). A georeferenced rRNA amplicon database of aquatic microbiomes from South America. Scientific Data, 9, 565. 10.1038/s41597-022-01665-z

Minot, S. S., Garb, B., Roldan, A., Tang, A. S., Oskotsky, T. T., Rosenthal, C., Hoffman, N. G., Sirota, M., & Golob, J. L. (2023). MaLiAmPi enables generalizable and taxonomy-independent microbiome features from technically diverse 16S-based microbiome studies. Cell Reports Methods, 3(11), 100639. 10.1016/j.crmeth.2023.100639

Nearing, J. T., Douglas, G. M., Comeau, A. M., & Langille, M. G. I. (2018). Denoising the denoisers: An independent evaluation of microbiome sequence error-correction approaches. PeerJ, 6, e5364. 10.7717/peerj.5364

North, B. V., Curtis, D., & Sham, P. C. (2002). A note on the calculation of empirical P values from Monte Carlo procedures. American Journal of Human Genetics, 71(2), 439–441. 10.1086/341527

Oksanen, J., Simpson, G. L., Blanchet, F. G., Kindt, R., Legendre, P., Minchin, P. R., O’Hara, R. B., Solymos, P., Stevens, M. H. H., Szoecs, E., & Wagner, H. (2025). vegan: Community ecology package (Version 2.7-5) [Computer software]. https://CRAN.R-project.org/package=vegan

Parada, A. E., Needham, D. M., & Fuhrman, J. A. (2016). Every base matters: Assessing small subunit rRNA primers for marine microbiomes with mock communities, time series and global field samples. Environmental Microbiology, 18(5), 1403–1414. 10.1111/1462-2920.13023

Peres-Neto, P. R., & Jackson, D. A. (2001). How well do multivariate data sets match? The advantages of a Procrustean superimposition approach over the Mantel test. Oecologia, 129(2), 169–178. 10.1007/s004420100720

Pinto, A. J., & Raskin, L. (2012). PCR biases distort bacterial and archaeal community structure in pyrosequencing datasets. PLoS ONE, 7(8), e43093. 10.1371/journal.pone.0043093

Pollock, J., Glendinning, L., Wisedchanwet, T., & Watson, M. (2018). The madness of microbiome: Attempting to find consensus “best practice” for 16S microbiome studies. Applied and Environmental Microbiology, 84(7), e02627–17. 10.1128/AEM.02627-17

Potapov, V., & Ong, J. L. (2017). Examining sources of error in PCR by single-molecule sequencing. PLOS ONE, 12(1), e0169774. 10.1371/journal.pone.0169774

Quast, C., Pruesse, E., Yilmaz, P., Gerken, J., Schweer, T., Yarza, P., Peplies, J., & Glöckner, F. O. (2013). The SILVA ribosomal RNA gene database project: Improved data processing and web-based tools. Nucleic Acids Research, 41(D1), D590–D596. 10.1093/nar/gks1219

R Core Team. (2023). R: A language and environment for statistical computing. R Foundation for Statistical Computing. https://www.R-project.org/

Ramirez, K. S., Knight, C. G., de Hollander, M., Brearley, F. Q., Constantinides, B., Cotton, A., Creer, S., Crowther, T. W., Davison, J., Delgado-Baquerizo, M., Dorrepaal, E., Elliott, D. R., Fox, G., Griffiths, R. I., Hale, C., Hartman, K., Houlden, A., Jones, D. L., Krab, E. J., … de Vries, F. T. (2018). Detecting macroecological patterns in bacterial communities across independent studies of global soils. Nature Microbiology, 3(2), 189–196. 10.1038/s41564-017-0062-x

Rideout, J. R., He, Y., Navas-Molina, J. A., Walters, W. A., Ursell, L. K., Gibbons, S. M., Chase, J., McDonald, D., Gonzalez, A., Robbins-Pianka, A., Clemente, J. C., Gilbert, J. A., Huse, S. M., Zhou, H.-W., Knight, R., & Caporaso, J. G. (2014). Subsampled open-reference clustering creates consistent, comprehensive OTU definitions and scales to billions of sequences. PeerJ, 2, e545. 10.7717/peerj.545

Rognes, T., Flouri, T., Nichols, B., Quince, C., & Mahé, F. (2016). VSEARCH: A versatile open source tool for metagenomics. PeerJ, 4, e2584. 10.7717/peerj.2584

Schirmer, M., D’Amore, R., Ijaz, U. Z., Hall, N., & Quince, C. (2016). Illumina error profiles: Resolving fine-scale variation in metagenomic sequencing data. BMC Bioinformatics, 17(1), 125. 10.1186/s12859-016-0976-y

Schirmer, M., Ijaz, U. Z., D’Amore, R., Hall, N., Sloan, W. T., & Quince, C. (2015). Insight into biases and sequencing errors for amplicon sequencing with the Illumina MiSeq platform. Nucleic Acids Research, 43(6), e37. 10.1093/nar/gku1341

Schloss, P. D., Gevers, D., & Westcott, S. L. (2011). Reducing the effects of PCR amplification and sequencing artifacts on 16S rRNA-based studies. PLoS ONE, 6(12), e27310. 10.1371/journal.pone.0027310

Shoemaker, W. R., Locey, K. J., & Lennon, J. T. (2017). A macroecological theory of microbial biodiversity. Nature Ecology & Evolution, 1(5), 0107. 10.1038/s41559-017-0107

Stackebrandt, E., & Goebel, B. M. (1994). Taxonomic note: A place for DNA-DNA reassociation and 16S rRNA sequence analysis in the present species definition in bacteriology. International Journal of Systematic Bacteriology, 44(4), 846–849. 10.1099/00207713-44-4-846

Thompson, L. R., Sanders, J. G., McDonald, D., Amir, A., Ladau, J., Locey, K. J., Prill, R. J., Tripathi, A., Gibbons, S. M., Ackermann, G., Navas-Molina, J. A., Janssen, S., Kopylova, E., Vázquez-Baeza, Y., González, A., Morton, J. T., Mirarab, S., Xu, Z. Z., Jiang, L., … Knight, R. (2017). A communal catalogue reveals Earth’s multiscale microbial diversity. Nature, 551(7681), 457–463. 10.1038/nature24621

Tremblay, J., Singh, K., Fern, A., Kirton, E. S., He, S., Woyke, T., Lee, J., Chen, F., Dangl, J. L., & Tringe, S. G. (2015). Primer and platform effects on 16S rRNA tag sequencing. Frontiers in Microbiology, 6, 771. 10.3389/fmicb.2015.00771

Vadillo Gonzalez, S., Jongen, R., Thomas, T., Marzinelli, E. M., & Gribben, P. E. (2026). Seagrass–microbe interactions: A systematic review of current research trends and mapping of the core microbiome. Biological Reviews, 101(3), 1334–1357. 10.1002/brv.70126

Van de Peer, Y., Chapelle, S., & De Wachter, R. (1996). A quantitative map of nucleotide substitution rates in bacterial rRNA. Nucleic Acids Research, 24(17), 3381–3391. 10.1093/nar/24.17.3381

Wang, Y., & Lê Cao, K.-A. (2023). PLSDA-batch: A multivariate framework to correct for batch effects in microbiome data. Briefings in Bioinformatics, 24(2), bbac622. 10.1093/bib/bbac622

Wright, E. S. (2015). DECIPHER: Harnessing local sequence context to improve protein multiple sequence alignment. BMC Bioinformatics, 16(1), 322. 10.1186/s12859-015-0749-z

Wright, E. S. (2016). Using DECIPHER v2.0 to analyze big biological sequence data in R. The R Journal, 8(1), 352–359. 10.32614/RJ-2016-025

