## Supplementary Information for "SeCAT (Sequence Consensus Amplicon Trimming): a simulation-informed Nextflow pipeline for cross-study harmonisation of 16S rRNA metabarcoding datasets"

*Contents: Supplementary Tables S1–S4, Supplementary Figures S1–S4 and Supplementary Data S1–S2 (supplied as separate PDF files).*

**Supplementary Table S1.** The thirteen validation datasets. Coordinate mapping, window support and inclusion columns are from the SeCAT paper run; source-study titles, geography and sample environments are from the ENA/NCBI BioProject records (accessed August 2026). Sequencing platform is from the ENA read-run records of the amplicon runs used.

| Study ID (manifest) | BioProject | Source study (ENA/NCBI title, abbreviated) | Geographic origin | Sample environment (ENA/NCBI record) | Primer set | Modal window (SILVA 138.2 columns) | ASVs mapped | Window support | Mapping quality | Samples retained | In consensus | In roster | Sequencing platform |
|---|---|---|---|---|---|---|---|---|---|---|---|---|---|
| PRJEB22038 | PRJEB22038 | SO243 cruise (sea-surface microlayer) | Peruvian upwelling, SE Pacific | Sea-surface microlayer / water column | 341F–805R | 6428–23454 | 814 | 1.00 | CONVERGED | 19 | Yes | Yes | Illumina MiSeq |
| PRJEB25224 | PRJEB25224 | Malaspina picoplankton, surface 16S | Global tropical/subtropical surface ocean | Surface water picoplankton | 515F(-Y)–926R | 13862–27666 | 1407 | 0.99 | CONVERGED | — | Yes | No | Illumina MiSeq |
| PRJEB27168 | PRJEB27168 | Galapagos bacterioplankton under contrasting conditions | Galapagos Archipelago, Ecuador | Coastal water column (CO2 vent, upwelling, reference) | 341F–805R | 6428–23444 | 103 | 1.00 | CONVERGED | 5 | Yes | Yes | Illumina MiSeq |
| PRJEB36282 | PRJEB36282 | Tara Oceans 16S amplicon, prokaryote fractions | Global ocean | Open-ocean water column | 515F(-Y)–926R | 13862–27684 | 3273 | 1.00 | CONVERGED | 59 | Yes | Yes | Illumina HiSeq 2500 |
| PRJEB42499 | PRJEB42499 | PS113 Atlantic Microbiome (RV Polarstern) | Atlantic Ocean transect | Water column | 515F–806R | 13862–23567 | 2782 | 1.00 | CONVERGED | 121 | Yes | Yes | Illumina MiSeq |
| PRJEB50983 | PRJEB50983 | Atlantic pro-/eukaryotic communities: primer choice matters | Atlantic Ocean, 51°S–47°N | Water column | 515F(-Y)–926R | 13862–27728 | 5145 | 1.00 | CONVERGED | 321 | Yes | Yes | Illumina MiSeq |
| PRJEB52328 | PRJEB52328 | Tara Pacific 16S V4V5, island-associated plankton | Pacific Ocean (Tara Pacific) | Island/reef-associated water column | 515F(-Y)–926R | 13862–27859 | 4988 | 0.99 | CONVERGED | 56 | Yes | Yes | Illumina HiSeq 2500 |
| PRJEB90443 | PRJEB90443 | Environmental disturbance of coastal microbial communities | France and Chile, 26 coastal sites | Coastal waters (lagoons, fjords, estuaries, harbours) | 515F(-Y)–926R | 13862–27675 | 1191 | 1.00 | CONVERGED | 15 | Yes | Yes | Illumina MiSeq |
| PRJNA263621 | PRJNA263621 | Size-fractioned community transcription, OMZ nitrogen metabolism | Eastern Tropical North Pacific OMZ | Water column (metatranscriptome project) | 515F–806R | 6238–37980 | 3104 | 0.24 | UNSTABLE | — | No | No | Illumina MiSeq |
| PRJNA664813 | PRJNA664813 | Alfonso Basin water samples | Bay of La Paz, Gulf of California, Mexico | Water column (oxygen-depleted basin) | 341F–805R | 6428–23448 | 623 | 1.00 | CONVERGED | 18 | Yes | Yes | Illumina MiSeq |
| PRJNA734169 | PRJNA73416 | MG_SAObay | San Antonio Bay, | Sponge- | 515F– | 13862– | 665 | 1.00 | CONVERGE | 31 | Yes | Yes | Illumina |

| Study ID (manifest) | BioProject | Source study (ENA/NCBI title, abbreviated) | Geographic origin | Sample environment (ENA/NCBI record) | Primer set | Modal window (SILVA 138.2 columns) | ASVs mapped | Window support | Mapping quality | Samples retained | In consensus | In roster | Sequencing platform |
|---|---|---|---|---|---|---|---|---|---|---|---|---|---|
| | 9 | (Hymeniacidon perlevis microbiome) | North Patagonia, Argentina | associated (intertidal) | 806R | 23444 | | | D | | | | MiniSeq |
| PRJNA780024 | PRJNA780024 | Oxygen minimum zone of the Mexican Pacific | Mexican Pacific OMZ (Mazatlán) | Seawater (OMZ) | 341F–805R | 6428–23449 | 363 | 1.00 | CONVERGED | 10 | Yes | Yes | Illumina MiSeq |
| PRJNA903199 | PRJNA903199 | Microbial communities of invasive kelp Undaria pinnatifida | Patagonia, Argentina | Kelp-associated and surrounding water | 515F–806R | 13862–23479 | 944 | 1.00 | CONVERGED | 27 | Yes | Yes | Illumina MiSeq |

*Sources: mapping and inclusion columns from the SeCAT paper run (Table 3; supporting/provenance manifest); titles, geography and sample environment from ENA/NCBI BioProject records (accessed August 2026). Platform column: instrument model of the amplicon runs in the ENA read-run records (accessed October 2026); PRJEB52328 also holds NovaSeq 6000 runs, but all 56 of its runs used here were sequenced on the HiSeq 2500. Sample counts are retained samples after merging; the two studies retained for consensus determination but not rostered carry no sample count.*

**Supplementary Table S2.** Compute environment and execution timeline of the demonstration run. The pipeline ran in two phases on the JASMIN LOTUS2 SLURM cluster in published container images; per-module statistics are from the Nextflow task logs of the Phase 1 (voluminous_pauling, total wall time 3 h 39 m) and Phase 2 (intergalactic_moriondo, 1 h 30 m) paper runs.

| Phase | Module (Nextflow process) | Tasks | Status | Wall time (per task) | CPU | Peak memory | Notes |
|---|---|---|---|---|---|---|---|
| 1 | CLEAN_DATA | 1 | COMPLETED | 32.3 s | 86.5% | 761 MB | manifest QC and staging |
| 1 | STUDY_MAPPING | 13 | COMPLETED | median 2 m 7 s (1 m 13 s – 6 m 58 s) | 53–89% | 25.0–28.9 GB | one task per study; memory dominated by the SILVA 138.2 alignment |
| 1 | COLLECT_MAPS | 1 | COMPLETED | 0.1 s | 33.7% | 3.2 MB | |
| 1 | PREPARE_SIMS | 1 | COMPLETED | 22 m 13 s | 97.9% | 2.8 GB | consensus determination and simulation set-up |
| 1 | RUN_SIMULATION | 1,200 (100 × 12 converged studies) | COMPLETED | median 2 m 13 s (36 s – 12 m 27 s) | 72–98% | 1.0–3.9 GB | independent SLURM tasks; fully parallel |
| 1 | ANALYSE_REAL | 12 | COMPLETED | median 1 m 49 s (31 s – 8 m 48 s) | 95–99% | 0.5–1.6 GB | one task per converged study |
| 1 | AGGREGATE | 1 | COMPLETED | 3 m 52 s | 95.5% | 902 MB | detector consensus and verdicts |
| 1 | GENERATE_VERDICTS | 1 | COMPLETED | 1.6 s | 84.8% | 140 MB | |
| 1 | GENERATE_REPORT | 1 | COMPLETED | 16 m 24 s | 96.0% | 6.7 GB | per-study reports and roster pause |
| 2 | STANDARDIZE:TRIM_SEQUENCES | 1 | COMPLETED | 24.6 s | 69.8% | 899 MB | |
| 2 | STANDARDIZE:MERGE_DATASETS | 1 | COMPLETED | 30.9 s | 73.0% | 1.0 GB | |
| 2 | STANDARDIZE:VALIDATE | 1 | COMPLETED | 1 h 16 m 30 s | 67.3% | 4.2 GB | full five-tier validation incl. SPIEC-EASI networks |

*Compute environment: JASMIN LOTUS2 SLURM cluster (Singularity 4.5.0; Nextflow 26.04.3); Phase 1 ran the released secat v5.0.6 container, Phase 2 the secat v5.0.5 image (identical dependency versions; the v5.0.6 image was not yet published at run time). Phase 1 statistics from the Nextflow task log of the Phase 1 paper run (voluminous_pauling, 2026-09-17; total wall time 3 h 39 m; all 1,231 tasks executed, none cached). Phase 2 from the standardize run that produced the released dataset (intergalactic_moriondo, 2026-09-03; total wall time 1 h 30 m including scheduler queueing; all tasks executed, none cached).*

**Supplementary Table S3.** Full signal-preservation validation of the harmonised dataset. (a) Sequence staging before and after trimming to the consensus window; (b) signal-preservation metrics at three taxonomic levels (expanded from Table 4); (c) co-occurrence network detail underlying the network rows.

##### (a) Sequence staging

| Sequence staging (Tier 0) | Before trimming | After trimming to consensus |
|---|---|---|
| Sequences (rostered ASVs) | 20,891 | 12,667 |
| Length coefficient of variation | 0.147 | 0.019 |
| Median sequence length (bp) | 372 | 249 |
| Mean GC content | 0.515 | 0.514 |
| Trim success | — | 100% |

##### (b) Signal preservation

| Metric | ASV | Genus | Family | Criterion |
|---|---|---|---|---|
| Beta-diversity (Mantel r) | 0.790 | 1.000 | 1.000 | > 0.85 |
| Procrustes $m^2$ | 0.522 | <0.001 | <0.001 | < 0.10 |
| PROTEST correlation | 0.692 | 1.000 | 1.000 | > 0.95 |
| PERMANOVA $\Delta R^2$ (StudyID) | −0.039 | −0.001 | +0.0002 | ≤ 0.05 |
| Dispersion change | 1.9% | 0.1% | <0.1% | < 10% |
| Alpha ICC (Shannon) | 0.927 | 0.999 | 1.000 | > 0.90 |
| Rank abundance (Spearman ρ) | 0.943 | 0.995 | 0.998 | > 0.90 |
| Abundance concordance (ρ) | n/a | 0.974 | 0.988 | > 0.90 |
| Network vertex / edge change | −0.6% / −15.9% | −1.2% / −7.9% | +1.3% / +1.6% | < 20% |
| Hub node retention | 0.23 (5/22) | 0.89 (8/9) | 1.00 (8/8) | ≥ 0.80 |
| Cross-study sharing / Jaccard | 0.211 | 0.207 | 0.335 | ≥ baseline |

##### (c) Network detail

| Network detail (SPIEC-EASI, prevalence-filtered) | ASV | Genus | Family |
|---|---|---|---|
| Vertices, before → after | 333 → 331 | 86 → 85 | 78 → 79 |
| Edges, before → after | 1734 → 1459 | 139 → 128 | 122 → 124 |
| Mappable pre-harmonisation hubs retained as hubs | 5/22 | 8/9 | 8/8 |
| Vertex-degree Spearman ρ (features in both networks) | 0.28 (n = 219) | — | — |

*All values from the SeCAT paper run outputs (final_outputs: 4_verification/validation_summary.csv and the GraphML co-occurrence networks; Zenodo outputs v3). ASV-rank hub retention and the vertex-degree correlation are computed from the GraphML networks through asv_mapping_final.tsv.*

**Supplementary Table S4.** Full per-study, per-taxonomic-level degradation verdict table (13 studies × 6 levels, 78 tests). Detector thresholds are SILVA 138.2 alignment columns; the verdict class applies the manuscript rule to the twelve eligible studies (31 pass, 38 caution, 3 fail); the excluded study is marked.

| Study | Primer | Level | Threshold_Required | Threshold_Observed_Changepoint | Threshold_Observed_Cutoff | Threshold_Observed_NullModel | Consensus_Status | Confirmed_Threshold_BP | N_Methods_Triggered | N_Methods_Agree | Verdict_class |
|---|---|---|---|---|---|---|---|---|---|---|---|
| PRJEB22038 | 341F_805R | Phylum | 7444 | 5000 | — | — | WARNING_SINGLE | — | 1 | 1 | caution |
| PRJEB22038 | 341F_805R | Class | 7444 | 1200 | — | — | WARNING_SINGLE | — | 1 | 1 | caution |
| PRJEB22038 | 341F_805R | Order | 7444 | 9100 | — | — | WARNING_SINGLE | — | 1 | 1 | pass |
| PRJEB22038 | 341F_805R | Family | 7444 | 9400 | — | — | WARNING_SINGLE | — | 1 | 1 | pass |
| PRJEB22038 | 341F_805R | Genus | 7444 | 4900 | — | — | WARNING_SINGLE | — | 1 | 1 | caution |
| PRJEB22038 | 341F_805R | ASV | 7444 | 8600 | — | — | WARNING_SINGLE | — | 1 | 1 | pass |
| PRJEB25224 | 515F-Y_926R | Phylum | 4222 | 1600 | — | — | WARNING_SINGLE | — | 1 | 1 | caution |
| PRJEB25224 | 515F-Y_926R | Class | 4222 | 5600 | — | — | WARNING_SINGLE | — | 1 | 1 | pass |
| PRJEB25224 | 515F-Y_926R | Order | 4222 | 4100 | — | — | WARNING_SINGLE | — | 1 | 1 | caution |
| PRJEB25224 | 515F-Y_926R | Family | 4222 | 1900 | — | — | WARNING_SINGLE | — | 1 | 1 | caution |
| PRJEB25224 | 515F-Y_926R | Genus | 4222 | 4400 | — | 700 | WARNING_SINGLE | — | 2 | 1 | caution |
| PRJEB25224 | 515F-Y_926R | ASV | 4222 | 1900 | — | 1500 | CONFIRMED | 1700 | 2 | 2 | fail |
| PRJEB27168 | 341F_805R | Phylum | 7434 | 7700 | — | — | WARNING_SINGLE | — | 1 | 1 | pass |
| PRJEB27168 | 341F_805R | Class | 7434 | 600 | — | — | WARNING_SINGLE | — | 1 | 1 | caution |
| PRJEB27168 | 341F_805R | Order | 7434 | 2200 | — | — | WARNING_SINGLE | — | 1 | 1 | caution |
| PRJEB27168 | 341F_805R | Family | 7434 | 8700 | — | — | WARNING_SINGLE | — | 1 | 1 | pass |
| PRJEB27168 | 341F_805R | Genus | 7434 | 5300 | — | — | WARNING_SINGLE | — | 1 | 1 | caution |
| PRJEB27168 | 341F_805R | ASV | 7434 | 600 | — | — | WARNING_SINGLE | — | 1 | 1 | caution |
| PRJEB36282 | 515F-Y_926R | Phylum | 4240 | 300 | — | — | WARNING_SINGLE | — | 1 | 1 | caution |
| PRJEB36282 | 515F-Y_926R | Class | 4240 | 3500 | — | — | WARNING_SINGLE | — | 1 | 1 | caution |
| PRJEB36282 | 515F-Y_926R | Order | 4240 | 6100 | — | — | WARNING_SINGLE | — | 1 | 1 | pass |
| PRJEB36282 | 515F-Y_926R | Family | 4240 | 3700 | — | — | WARNING_SINGLE | — | 1 | 1 | caution |
| PRJEB36282 | 515F-Y_926R | Genus | 4240 | 500 | — | — | WARNING_SINGLE | — | 1 | 1 | caution |
| PRJEB36282 | 515F-Y_926R | ASV | 4240 | 200 | — | 800 | WARNING_SINGLE | — | 2 | 1 | caution |
| PRJEB42499 | 515F_806R | Phylum | 123 | 1600 | — | — | WARNING_SINGLE | — | 1 | 1 | pass |
| PRJEB42499 | 515F_806R | Class | 123 | 1600 | — | — | WARNING_SINGLE | — | 1 | 1 | pass |
| PRJEB42499 | 515F_806R | Order | 123 | 1800 | — | — | WARNING_SINGLE | — | 1 | 1 | pass |
| PRJEB42499 | 515F_806R | Family | 123 | 1300 | — | — | WARNING_SINGLE | — | 1 | 1 | pass |

| Study | Primer | Level | Threshold_Required | Threshold_Observed_Changepoint | Threshold_Observed_Cutoff | Threshold_Observed_NullModel | Consensus_Status | Confirmed_Threshold_BP | N_Methods_Triggered | N_Methods_Agree | Verdict_class |
|---|---|---|---|---|---|---|---|---|---|---|---|
| PRJEB42499 | 515F_806R | Genus | 123 | 200 | — | — | WARNING_SINGLE | — | 1 | 1 | pass |
| PRJEB42499 | 515F_806R | ASV | 123 | 1600 | — | — | WARNING_SINGLE | — | 1 | 1 | pass |
| PRJEB50983 | 515F-Y_926R | Phylum | 4284 | 1500 | — | — | WARNING_SINGLE | — | 1 | 1 | caution |
| PRJEB50983 | 515F-Y_926R | Class | 4284 | 2400 | — | — | WARNING_SINGLE | — | 1 | 1 | caution |
| PRJEB50983 | 515F-Y_926R | Order | 4284 | 5700 | — | — | WARNING_SINGLE | — | 1 | 1 | pass |
| PRJEB50983 | 515F-Y_926R | Family | 4284 | 2000 | — | — | WARNING_SINGLE | — | 1 | 1 | caution |
| PRJEB50983 | 515F-Y_926R | Genus | 4284 | 6200 | — | — | WARNING_SINGLE | — | 1 | 1 | pass |
| PRJEB50983 | 515F-Y_926R | ASV | 4284 | 3400 | — | — | WARNING_SINGLE | — | 1 | 1 | caution |
| PRJEB52328 | 515F-Y_926R | Phylum | 4415 | 300 | — | 1500 | WARNING_SINGLE | — | 2 | 1 | caution |
| PRJEB52328 | 515F-Y_926R | Class | 4415 | 1600 | — | 1500 | CONFIRMED | 1550 | 2 | 2 | fail |
| PRJEB52328 | 515F-Y_926R | Order | 4415 | 6500 | — | — | WARNING_SINGLE | — | 1 | 1 | pass |
| PRJEB52328 | 515F-Y_926R | Family | 4415 | 600 | — | — | WARNING_SINGLE | — | 1 | 1 | caution |
| PRJEB52328 | 515F-Y_926R | Genus | 4415 | 200 | — | 1300 | WARNING_SINGLE | — | 2 | 1 | caution |
| PRJEB52328 | 515F-Y_926R | ASV | 4415 | 2600 | — | 1300 | WARNING_SINGLE | — | 2 | 1 | caution |
| PRJEB90443 | 515F-Y_926R | Phylum | 4231 | 3400 | — | — | WARNING_SINGLE | — | 1 | 1 | caution |
| PRJEB90443 | 515F-Y_926R | Class | 4231 | 5600 | — | — | WARNING_SINGLE | — | 1 | 1 | pass |
| PRJEB90443 | 515F-Y_926R | Order | 4231 | 4700 | — | — | WARNING_SINGLE | — | 1 | 1 | pass |
| PRJEB90443 | 515F-Y_926R | Family | 4231 | 3800 | — | — | WARNING_SINGLE | — | 1 | 1 | caution |
| PRJEB90443 | 515F-Y_926R | Genus | 4231 | 5500 | — | 2200 | WARNING_SINGLE | — | 2 | 1 | caution |
| PRJEB90443 | 515F-Y_926R | ASV | 4231 | 2400 | — | 5600 | WARNING_SINGLE | — | 2 | 1 | caution |
| PRJNA263621 | 515F_806R | Phylum | 22160 | 600 | 24200 | 100 | CONFIRMED | 350 | 3 | 2 | excluded (UNSTABLE) |
| PRJNA263621 | 515F_806R | Class | 22160 | 11400 | 24200 | 100 | WARNING_SINGLE | — | 3 | 1 | excluded (UNSTABLE) |
| PRJNA263621 | 515F_806R | Order | 22160 | 6300 | 24200 | 100 | WARNING_SINGLE | — | 3 | 1 | excluded (UNSTABLE) |
| PRJNA263621 | 515F_806R | Family | 22160 | 11200 | 24200 | 100 | WARNING_SINGLE | — | 3 | 1 | excluded (UNSTABLE) |
| PRJNA263621 | 515F_806R | Genus | 22160 | 17200 | 24200 | 100 | WARNING_SINGLE | — | 3 | 1 | excluded (UNSTABLE) |
| PRJNA263621 | 515F_806R | ASV | 22160 | 5800 | 24200 | 100 | HARD_FAIL | 21200 | 4 | 1 | excluded (UNSTABLE) |
| PRJNA664813 | 341F_805R | Phylum | 7438 | 1300 | — | — | WARNING_SINGLE | — | 1 | 1 | caution |

| Study | Primer | Level | Threshold_Required | Threshold_Observed_Changepoint | Threshold_Observed_Cutoff | Threshold_Observed_NullModel | Consensus_Status | Confirmed_Threshold_BP | N_Methods_Triggered | N_Methods_Agree | Verdict_class |
|---|---|---|---|---|---|---|---|---|---|---|---|
| PRJNA664813 | 341F_805R | Class | 7438 | 8500 | — | — | WARNING_SINGLE | — | 1 | 1 | pass |
| PRJNA664813 | 341F_805R | Order | 7438 | 5900 | — | — | WARNING_SINGLE | — | 1 | 1 | caution |
| PRJNA664813 | 341F_805R | Family | 7438 | 1600 | — | — | WARNING_SINGLE | — | 1 | 1 | caution |
| PRJNA664813 | 341F_805R | Genus | 7438 | 5500 | — | 1400 | WARNING_SINGLE | — | 2 | 1 | caution |
| PRJNA664813 | 341F_805R | ASV | 7438 | 6300 | — | 1300 | WARNING_SINGLE | — | 2 | 1 | caution |
| PRJNA734169 | 515F_806R | Phylum | 0 | 600 | — | — | WARNING_SINGLE | — | 1 | 1 | pass |
| PRJNA734169 | 515F_806R | Class | 0 | 2000 | — | — | WARNING_SINGLE | — | 1 | 1 | pass |
| PRJNA734169 | 515F_806R | Order | 0 | 1900 | — | — | WARNING_SINGLE | — | 1 | 1 | pass |
| PRJNA734169 | 515F_806R | Family | 0 | 2000 | — | — | WARNING_SINGLE | — | 1 | 1 | pass |
| PRJNA734169 | 515F_806R | Genus | 0 | 700 | — | — | WARNING_SINGLE | — | 1 | 1 | pass |
| PRJNA734169 | 515F_806R | ASV | 0 | 1400 | — | — | WARNING_SINGLE | — | 1 | 1 | pass |
| PRJNA780024 | 341F_805R | Phylum | 7439 | 2700 | — | 2200 | CONFIRMED | 2450 | 2 | 2 | fail |
| PRJNA780024 | 341F_805R | Class | 7439 | 1300 | — | 2200 | WARNING_SINGLE | — | 2 | 1 | caution |
| PRJNA780024 | 341F_805R | Order | 7439 | 7900 | — | 2200 | WARNING_SINGLE | — | 2 | 1 | caution |
| PRJNA780024 | 341F_805R | Family | 7439 | 7000 | — | 2200 | WARNING_SINGLE | — | 2 | 1 | caution |
| PRJNA780024 | 341F_805R | Genus | 7439 | 200 | — | 2200 | WARNING_SINGLE | — | 2 | 1 | caution |
| PRJNA780024 | 341F_805R | ASV | 7439 | 9300 | — | 1300 | WARNING_SINGLE | — | 2 | 1 | caution |
| PRJNA903199 | 515F_806R | Phylum | 35 | 900 | — | — | WARNING_SINGLE | — | 1 | 1 | pass |
| PRJNA903199 | 515F_806R | Class | 35 | 1900 | — | — | WARNING_SINGLE | — | 1 | 1 | pass |
| PRJNA903199 | 515F_806R | Order | 35 | 1200 | — | — | WARNING_SINGLE | — | 1 | 1 | pass |
| PRJNA903199 | 515F_806R | Family | 35 | 1900 | — | — | WARNING_SINGLE | — | 1 | 1 | pass |
| PRJNA903199 | 515F_806R | Genus | 35 | 200 | — | — | WARNING_SINGLE | — | 1 | 1 | pass |
| PRJNA903199 | 515F_806R | ASV | 35 | 2000 | — | — | WARNING_SINGLE | — | 1 | 1 | pass |

*Full per-study, per-rank verdict table from the SeCAT paper run (3_verdicts/master_verdict_table.csv). Verdict class derived by the manuscript rule: pass = no detector threshold at or before the required trim; fail = consensus CONFIRMED at or before the required trim; caution = uncorroborated signal before the required trim. PRJNA263621 excluded (UNSTABLE mapping). Eligible tests: 31 pass / 38 caution / 3 fail.*

**Supplementary Figure S1.** Alpha-diversity preservation through harmonisation. Per-sample diversity before versus after trimming to the consensus window and collapse to MetaASVs (top) and Bland–Altman agreement (bottom) at ASV, genus and family level. Shannon ICC: 0.927 (ASV), 0.999 (genus), 1.000 (family); full index-level values in Table S3.

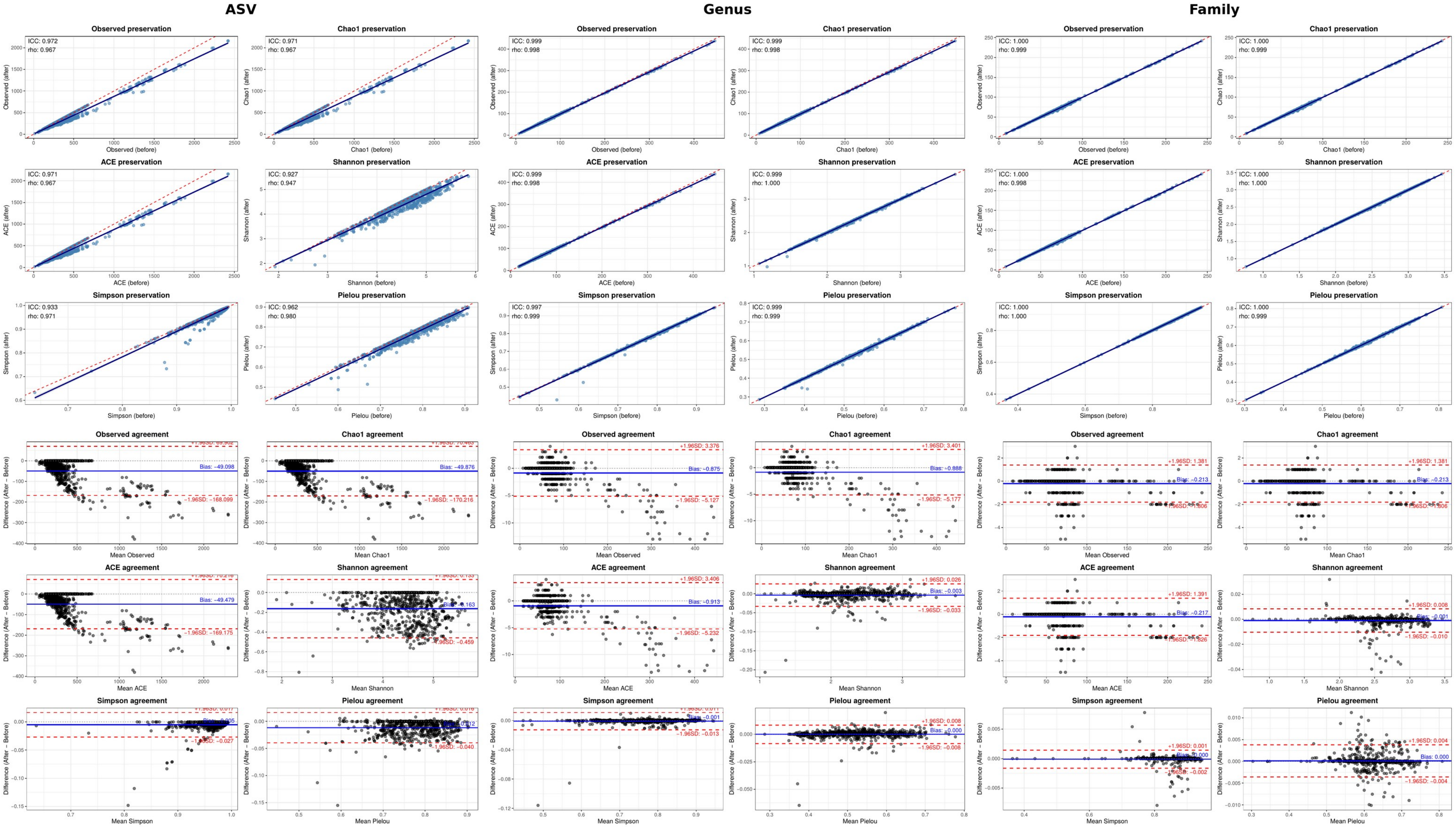

**Supplementary Figure S2.** Ordination geometry before and after harmonisation. Bray–Curtis PCoA of the pre- (top) and post-harmonisation (middle) datasets and Procrustes superimposition of the two ordinations (bottom) at ASV, genus and family level (Procrustes m² = 0.522, <0.001, <0.001; PROTEST r = 0.692, 1.000, 1.000; 999 permutations).

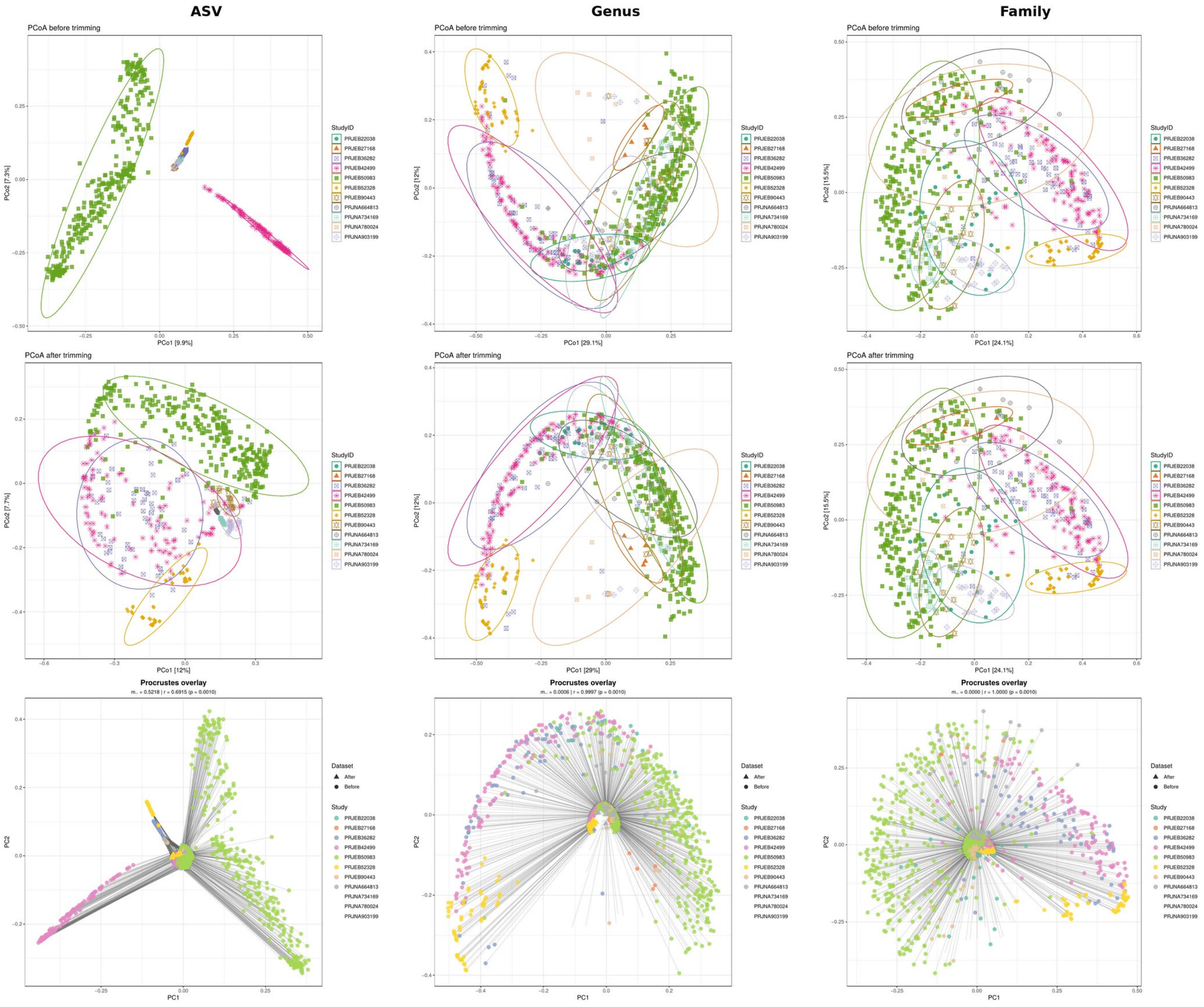

**Supplementary Figure S3.** Community structure diagnostics. Within-group dispersion (distance to centroid, top), per-sample rank-abundance concordance (middle) and per-taxon abundance concordance (bottom) at ASV, genus and family level. Abundance concordance is not computed at ASV level because features do not match across studies before harmonisation.

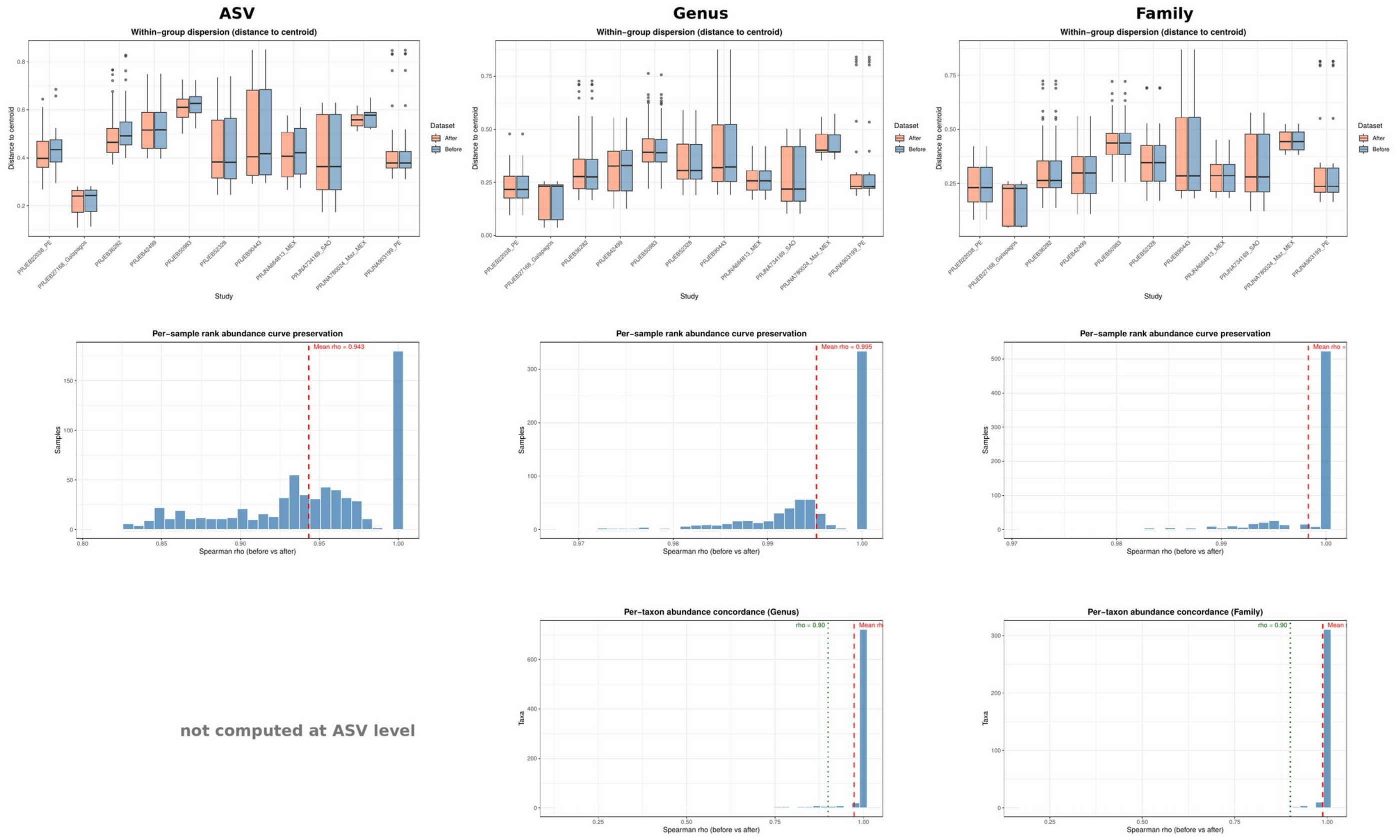

**Supplementary Figure S4.** Taxonomic composition before and after harmonisation at ASV, genus and family level.

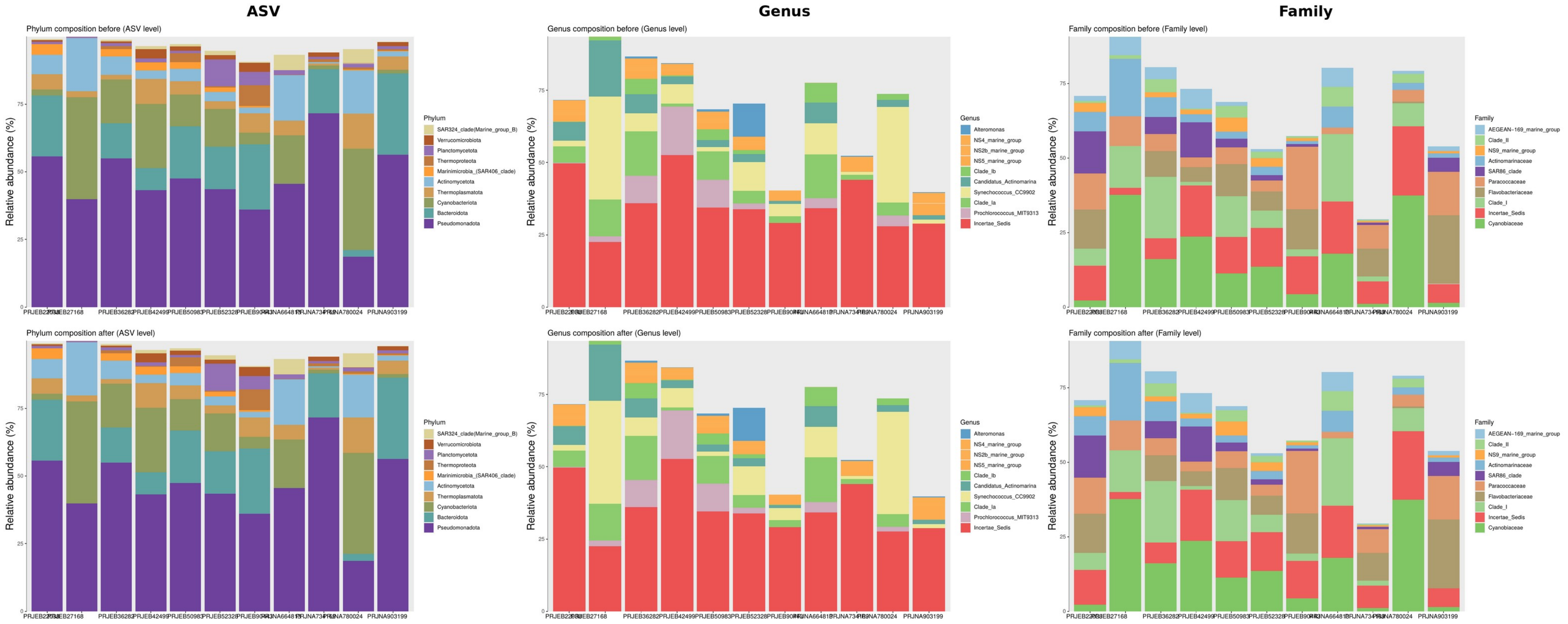

**Supplementary Data S1.** Per-study degradation diagnostics at ASV level (separate file: SeCAT_Supplementary_Data_S1_per-study_ASV_reports.pdf). One page per study, ordered by primer set, for the twelve analysed datasets (the UNSTABLE study receives no report by design). Each page reports, at ASV level: (A) simulated Bray–Curtis dissimilarity to the untrimmed baseline against trim step, with the detected degradation threshold marked; (B) taxon retention against trim step; (C) the taxa showing the largest change in relative abundance; and (D) the most abundant prevalence-filtered taxa. The page header gives the detector methods triggered, the observed threshold and the consensus verdict. Pages are extracted unaltered from the per-study diagnostic reports written by GENERATE_REPORT during the demonstration run; the full reports covering all six taxonomic levels are included in the archived run outputs (DOI 10.5281/zenodo.22871720).

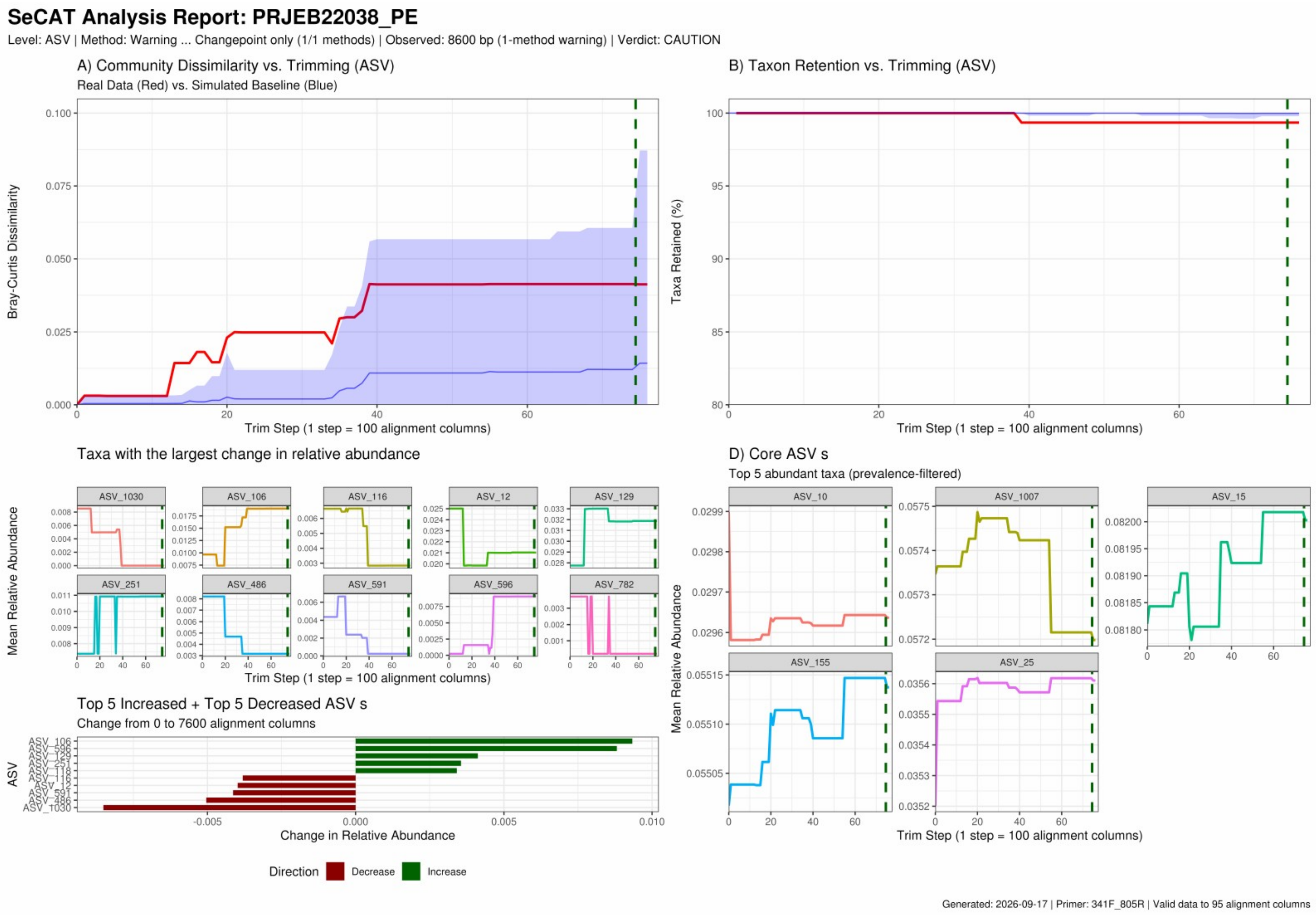


*Supplementary Data S1, page 1 of 12 — PRJEB22038_341F_805R*

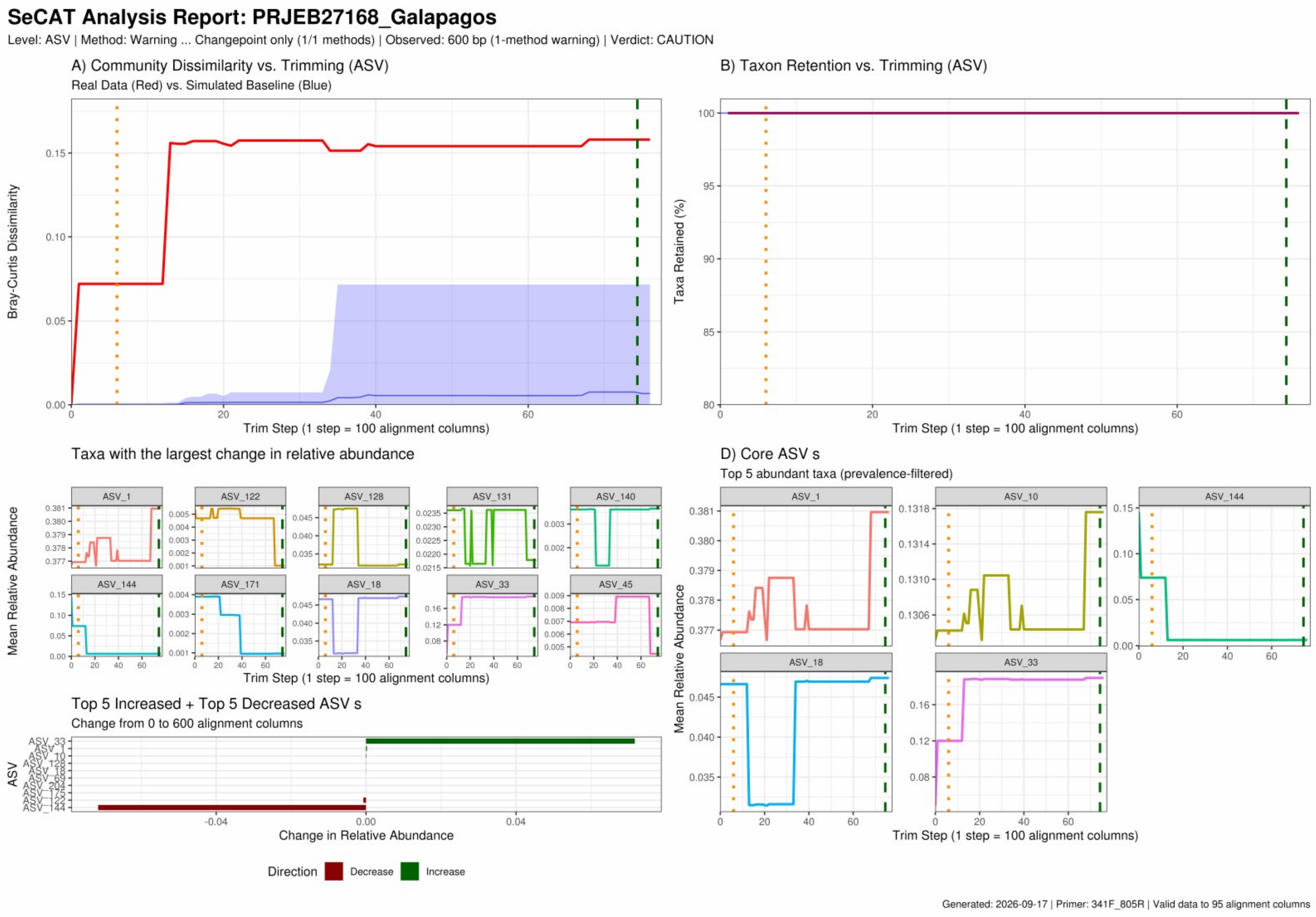


*Supplementary Data S1, page 2 of 12 — PRJEB27168_341F_805R*

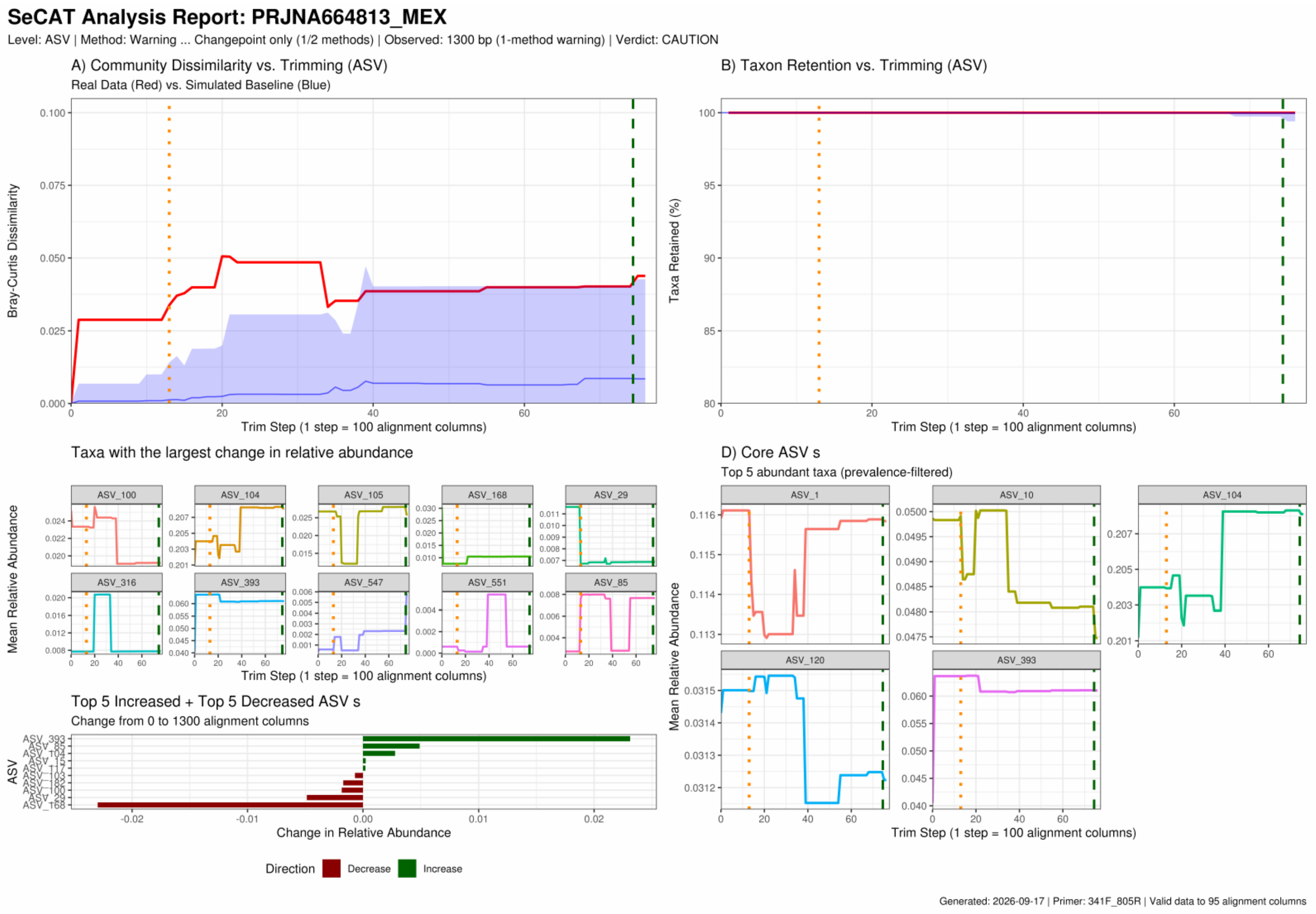


*Supplementary Data S1, page 3 of 12 — PRJNA664813_341F_805R*

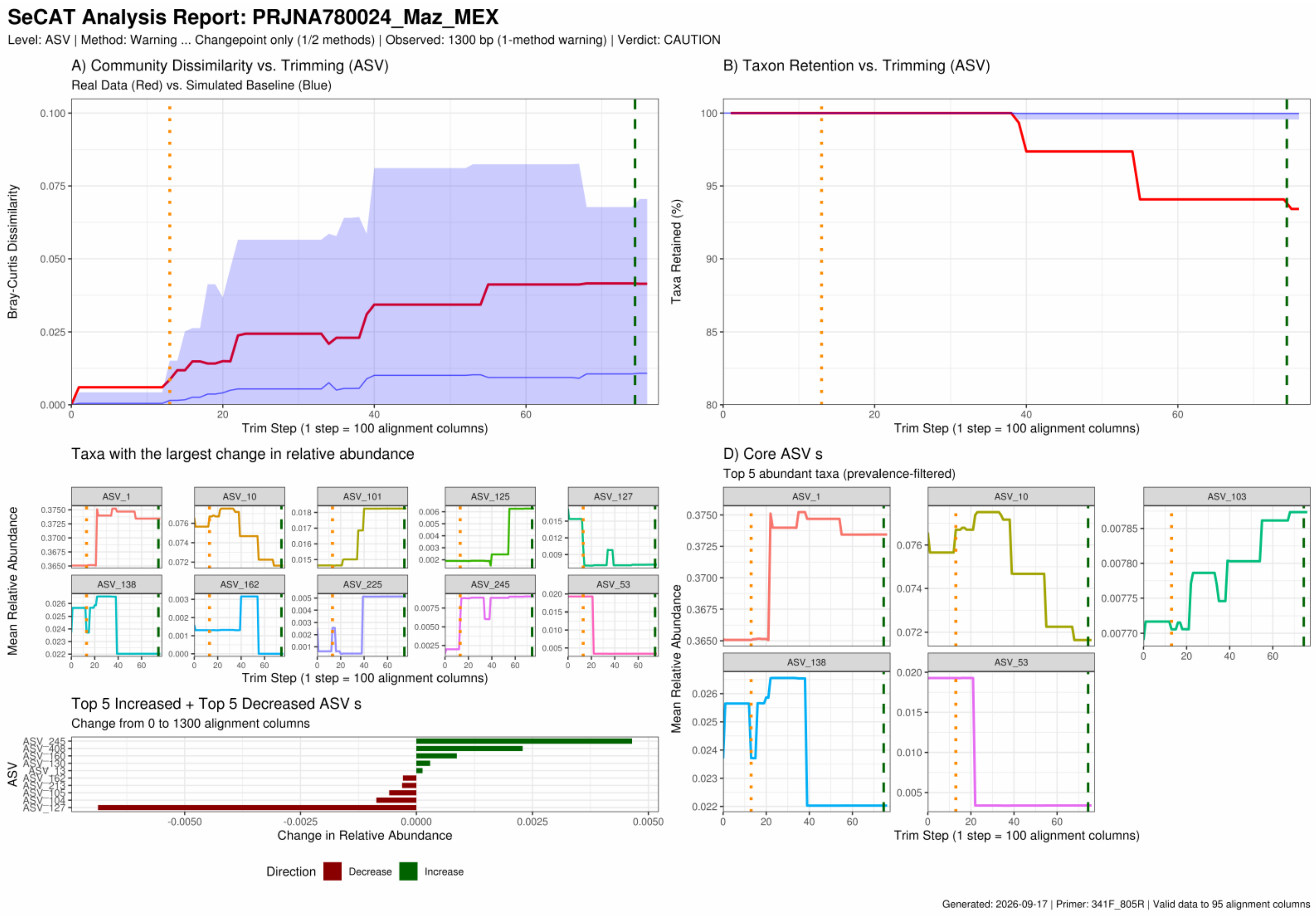


*Supplementary Data S1, page 4 of 12 — PRJNA780024_341F_805R*

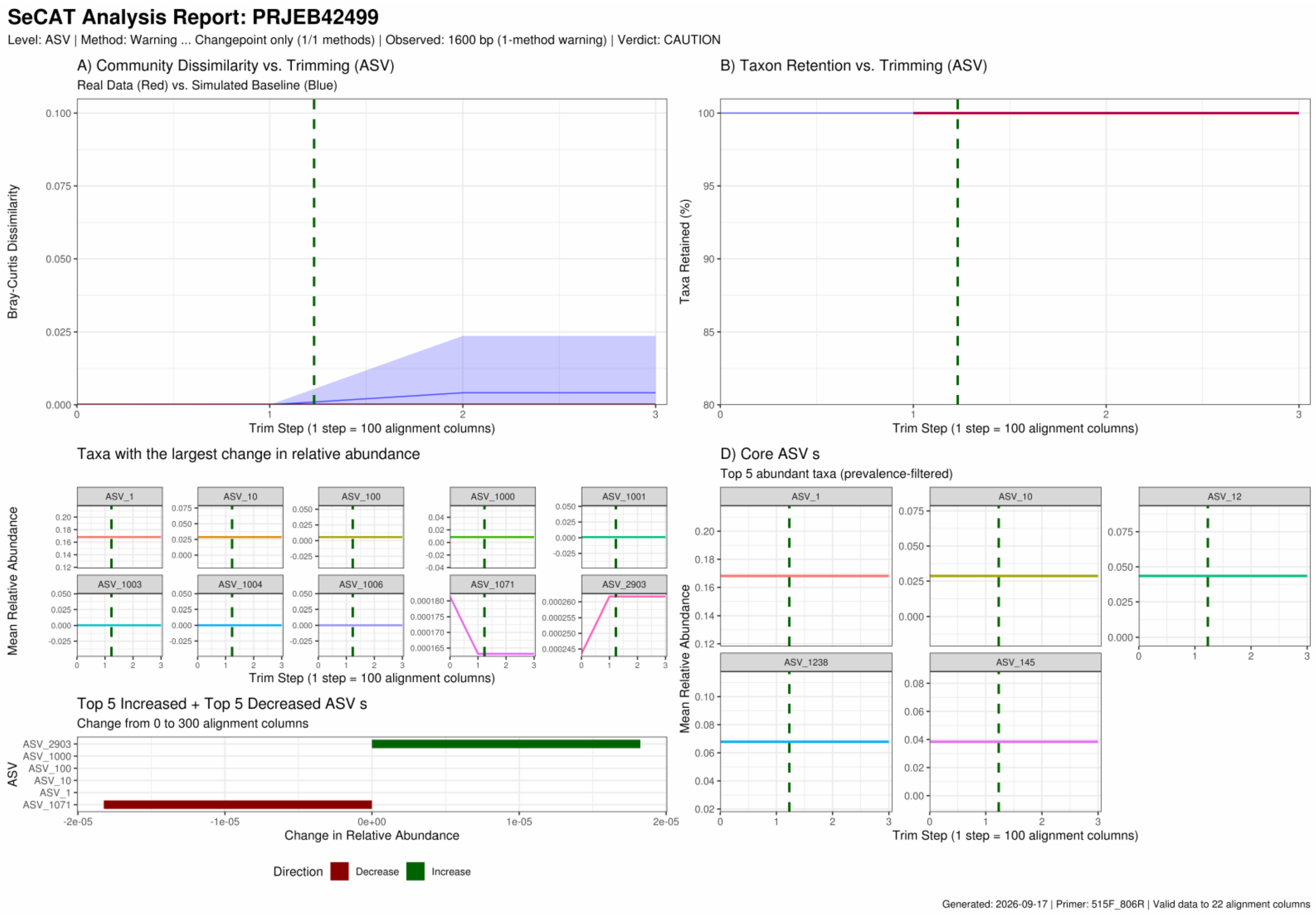


*Supplementary Data S1, page 5 of 12 — PRJEB42499_515F_806R*

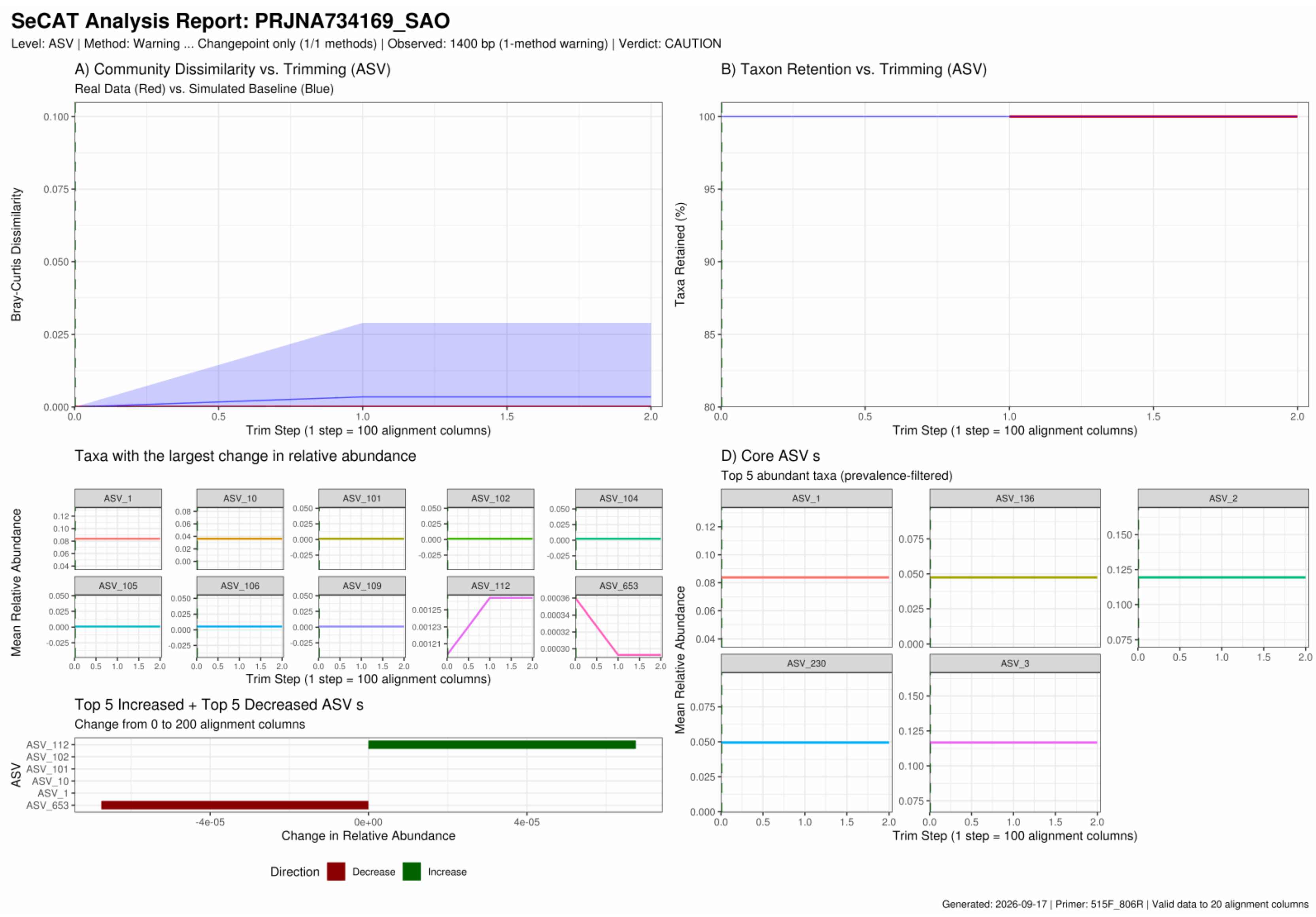


*Supplementary Data S1, page 6 of 12 — PRJNA734169_515F_806R*

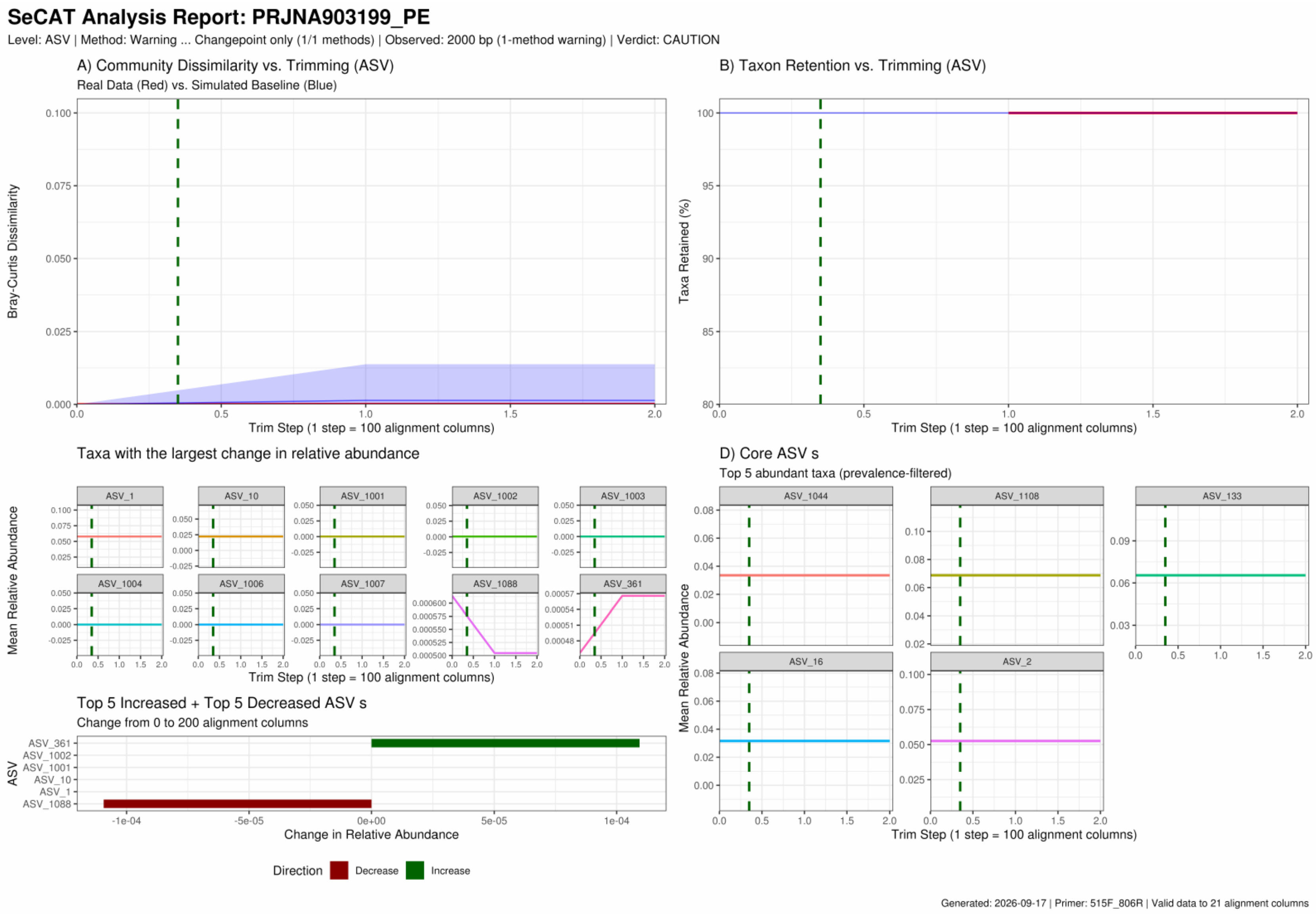


*Supplementary Data S1, page 7 of 12 — PRJNA903199_515F_806R*

#### SeCAT Analysis Report: PRJEB25224

Level: ASV | Method: Consensus (2 methods agree) | Observed: 1700 alignment columns | Verdict: FAIL

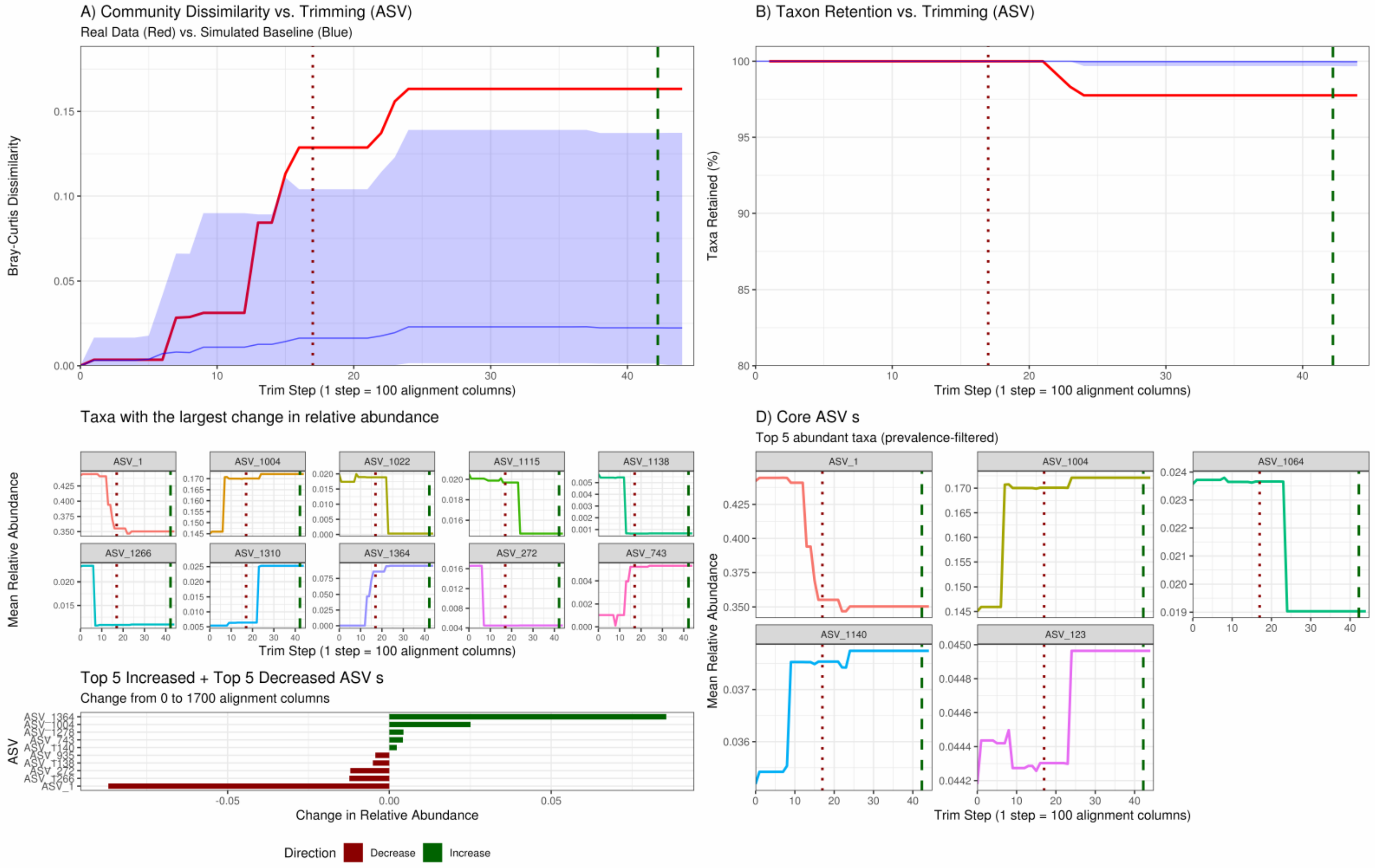


Generated: 2026-09-17 | Primer: 515F-Y_926R | Valid data to 63 alignment columns

### SeCAT Analysis Report: PRJEB36282

Level: ASV | Method: Warning ... Changepoint only (1/2 methods) | Observed: 200 bp (1-method warning) | Verdict: CAUTION

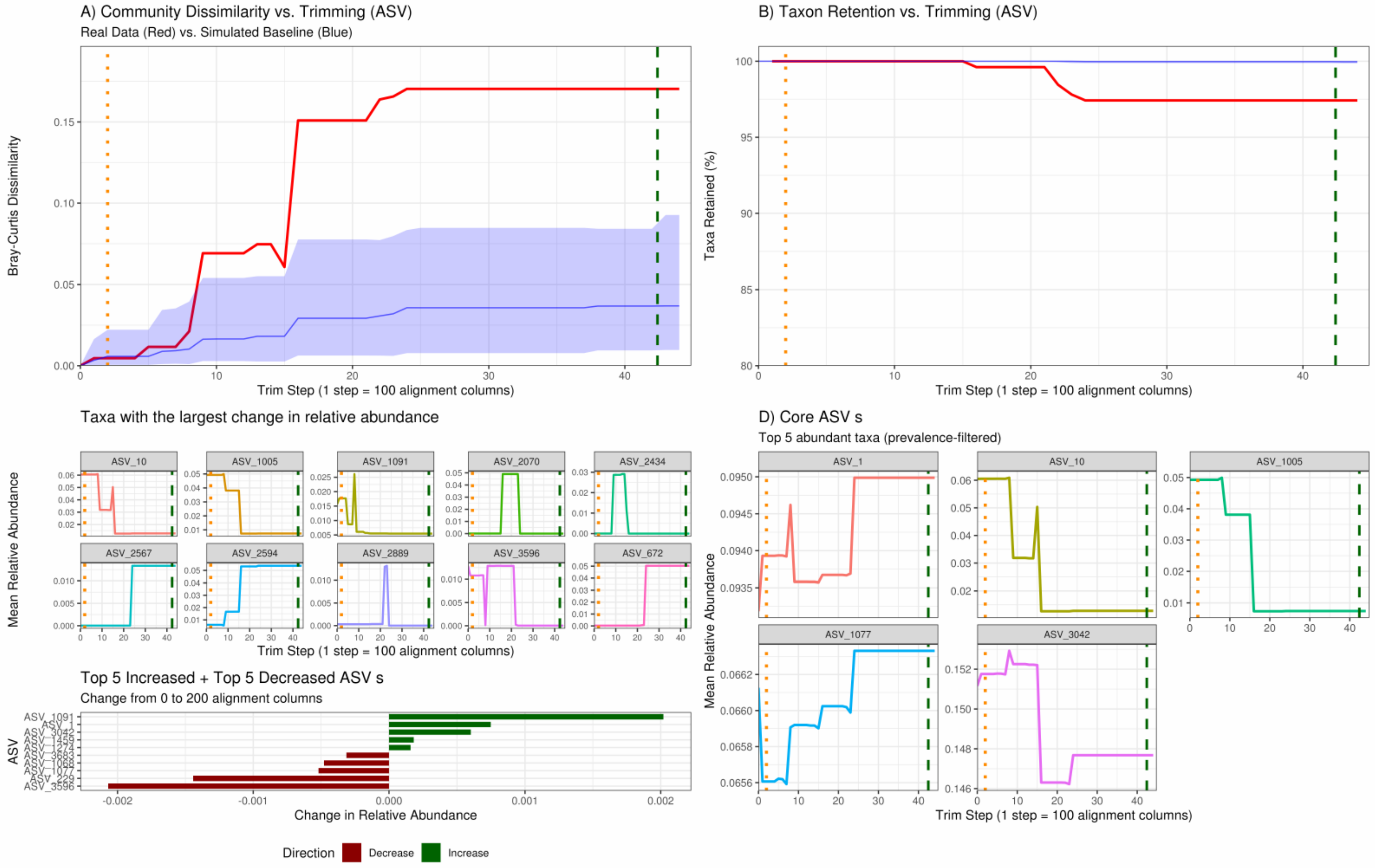


Generated: 2026-09-17 | Primer: 515F-Y_926R | Valid data to 63 alignment columns

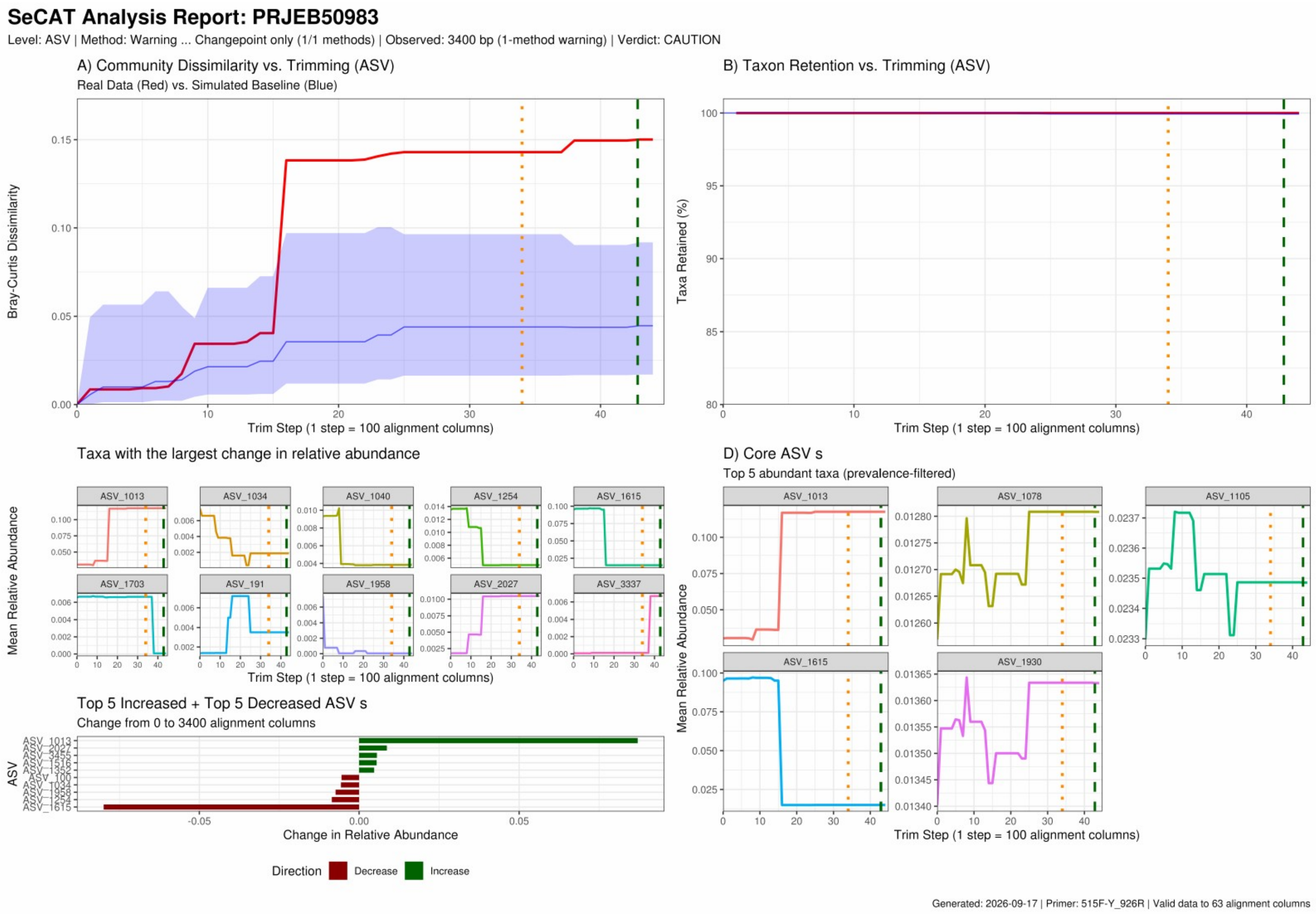


*Supplementary Data S1, page 10 of 12 — PRJEB50983_515F-Y_926R*

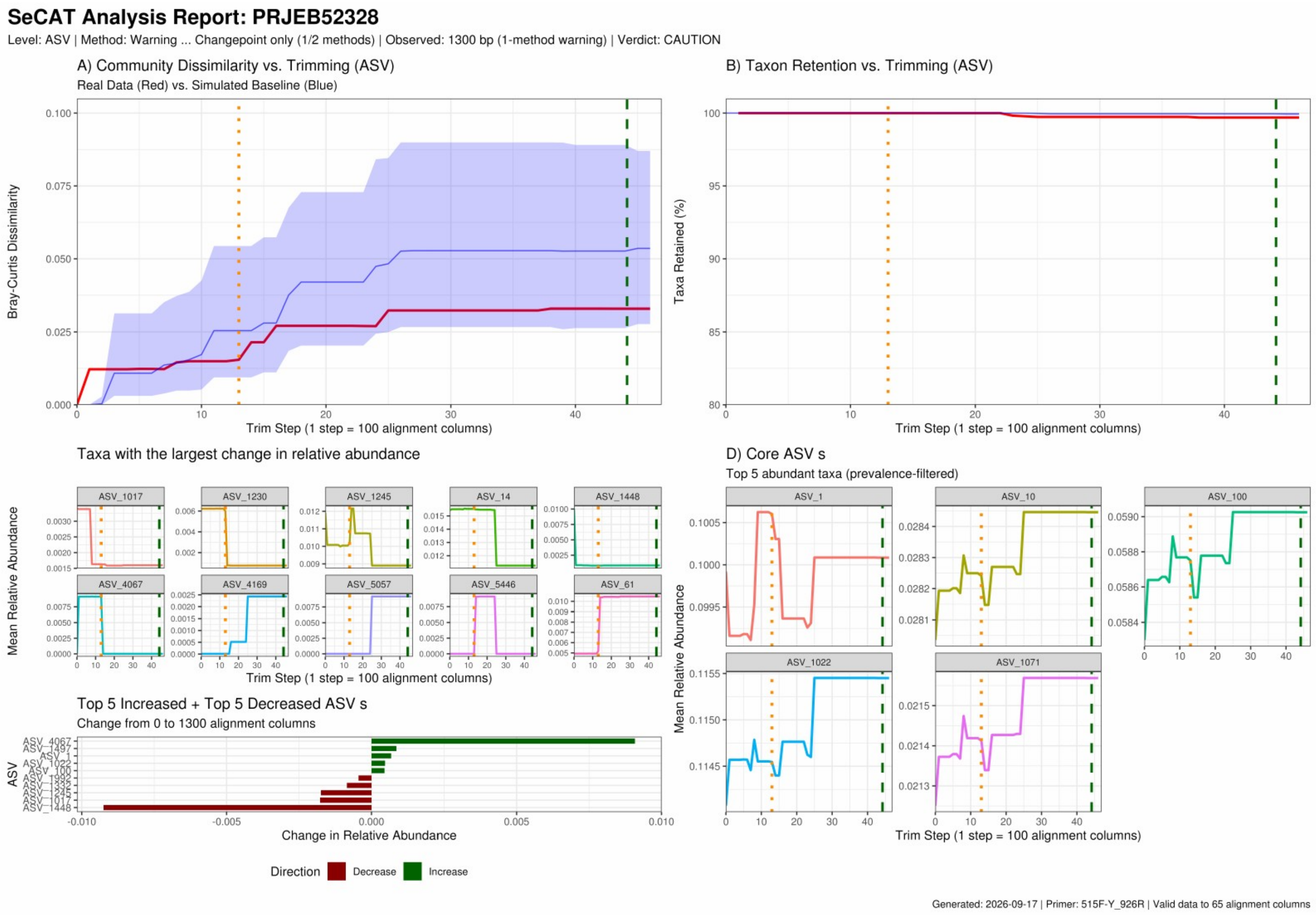
SeCAT Analysis Report: PRJEB52328
Level: ASV | Method: Warning ... Changepoint only (1/2 methods) | Observed: 1300 bp (1-method warning) | Verdict: CAUTION
A) Community Dissimilarity vs. Trimming (ASV)
Real Data (Red) vs. Simulated Baseline (Blue)
Bray-Curtis Dissimilarity
Trim Step (1 step = 100 alignment columns)
B) Taxon Retention vs. Trimming (ASV)
Taxa Retained (%)
Trim Step (1 step = 100 alignment columns)
Taxa with the largest change in relative abundance
ASV_1017
ASV_1230
ASV_1245
ASV_14
ASV_1448
ASV_4067
ASV_4169
ASV_5057
ASV_5446
ASV_61
Mean Relative Abundance
Trim Step (1 step = 100 alignment columns)
D) Core ASV s
Top 5 abundant taxa (prevalence-filtered)
ASV_1
ASV_10
ASV_100
ASV_1022
ASV_1071
Mean Relative Abundance
Trim Step (1 step = 100 alignment columns)
Top 5 Increased + Top 5 Decreased ASV s
Change from 0 to 1300 alignment columns
ASV
Change in Relative Abundance
Direction
Decrease
Increase
Generated: 2026-09-17 | Primer: 515F-Y_926R | Valid data to 65 alignment columns

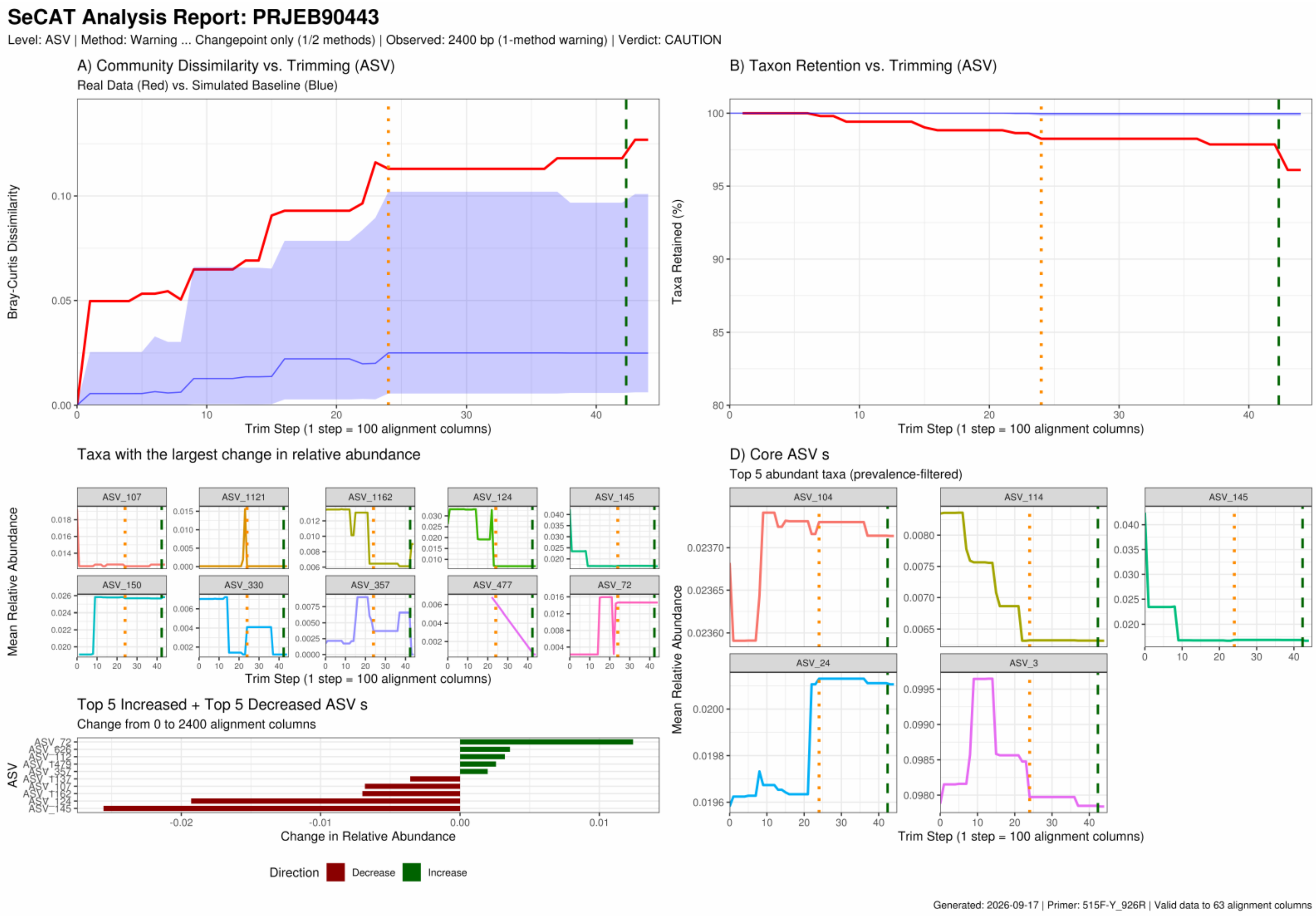
SeCAT Analysis Report: PRJEB90443
Level: ASV | Method: Warning ... Changepoint only (1/2 methods) | Observed: 2400 bp (1-method warning) | Verdict: CAUTION
A) Community Dissimilarity vs. Trimming (ASV)
Real Data (Red) vs. Simulated Baseline (Blue)
Bray-Curtis Dissimilarity
Trim Step (1 step = 100 alignment columns)
B) Taxon Retention vs. Trimming (ASV)
Taxa Retained (%)
Trim Step (1 step = 100 alignment columns)
Taxa with the largest change in relative abundance
Mean Relative Abundance
ASV_107
ASV_1121
ASV_1162
ASV_124
ASV_145
ASV_150
ASV_330
ASV_357
ASV_477
ASV_72
Trim Step (1 step = 100 alignment columns)
D) Core ASV s
Top 5 abundant taxa (prevalence-filtered)
ASV_104
ASV_114
ASV_145
ASV_24
ASV_3
Mean Relative Abundance
Trim Step (1 step = 100 alignment columns)
Top 5 Increased + Top 5 Decreased ASV s
Change from 0 to 2400 alignment columns
ASV
Change in Relative Abundance
Direction
Decrease
Increase
Generated: 2026-09-17 | Primer: 515F-Y_926R | Valid data to 63 alignment columns

**Supplementary Data S2.** Master summary report of the demonstration run (separate file: SeCAT_Supplementary_Data_S2_master_summary_report.pdf). Seven pages summarising all thirteen mapped studies together: overall amplicon alignment and safe trimming regions at each of the six taxonomic levels, with each study's mapped window drawn on the SILVA alignment frame and segments coloured by trimming safety relative to the consensus region, followed by a cross-study summary of detector performance. Reproduced unaltered from the run outputs.

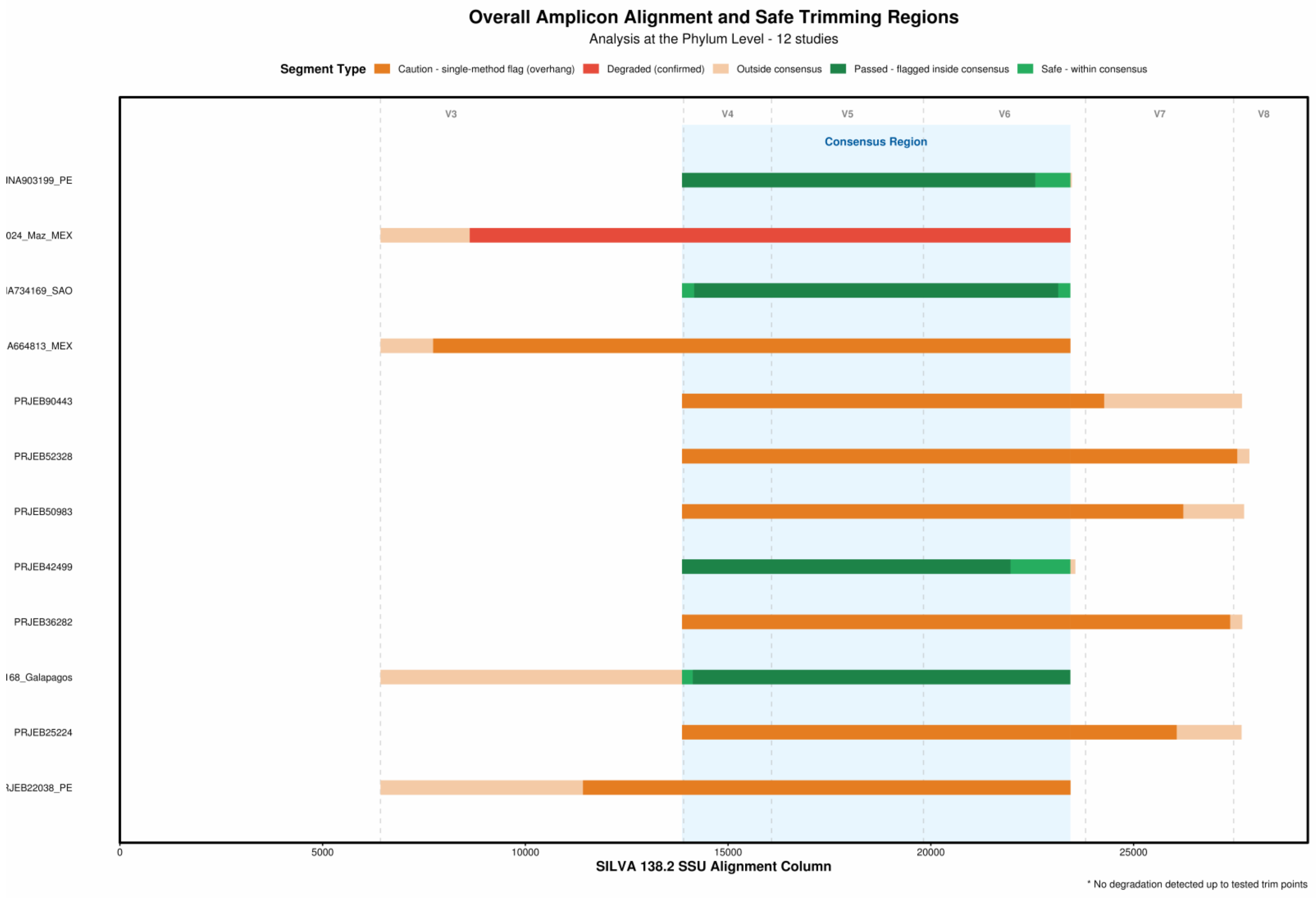


*Supplementary Data S2, page 1 of 7*

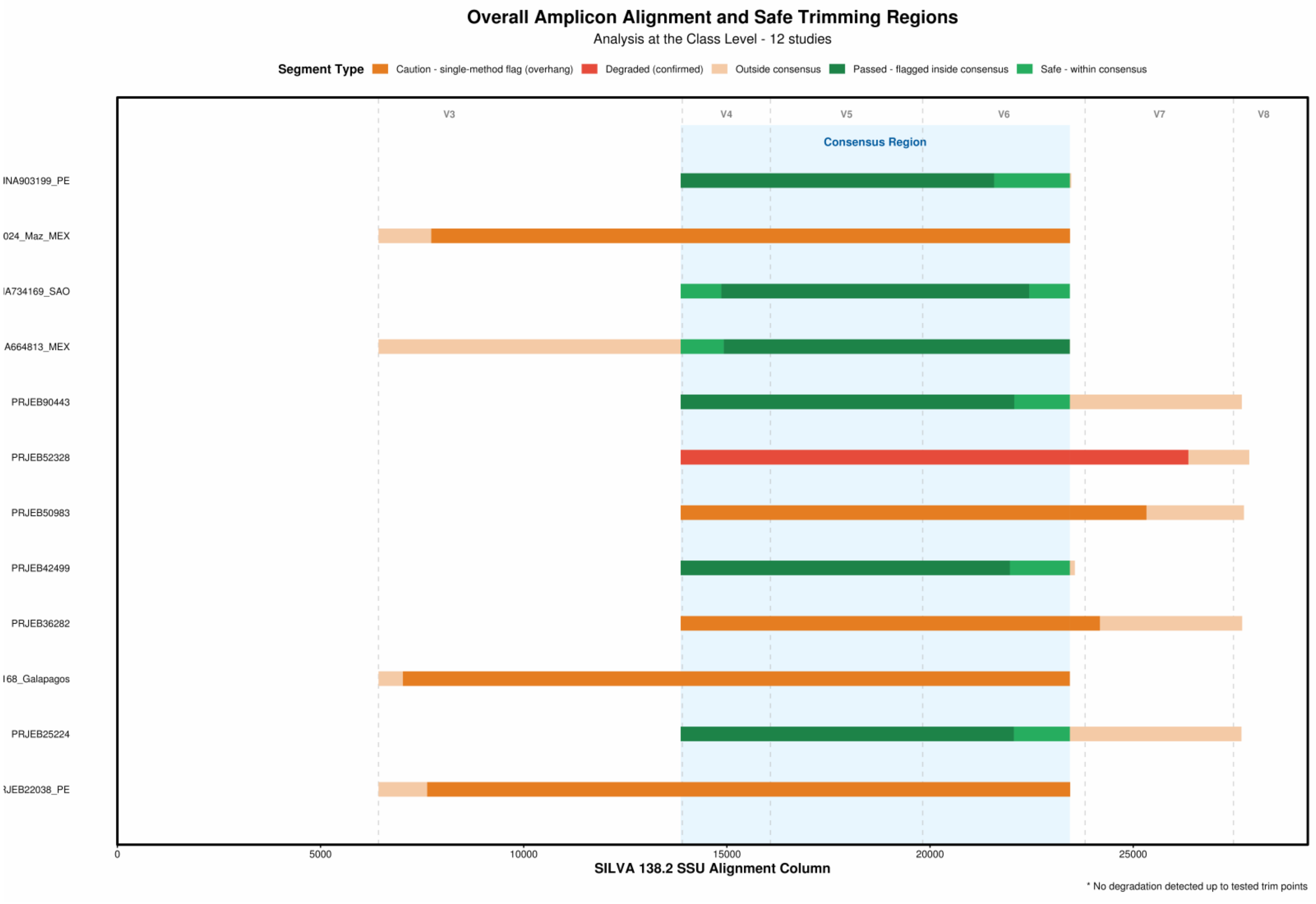

Overall Amplicon Alignment and Safe Trimming Regions
Analysis at the Class Level - 12 studies
Segment Type
Caution - single-method flag (overhang)
Degraded (confirmed)
Outside consensus
Passed - flagged inside consensus
Safe - within consensus
V3
V4
V5
V6
V7
V8
Consensus Region
INA903199_PE
024_Maz_MEX
IA734169_SAO
A664813_MEX
PRJEB90443
PRJEB52328
PRJEB50983
PRJEB42499
PRJEB36282
168_Galapagos
PRJEB25224
JEB22038_PE
0
5000
10000
15000
20000
25000
SILVA 138.2 SSU Alignment Column
* No degradation detected up to tested trim points

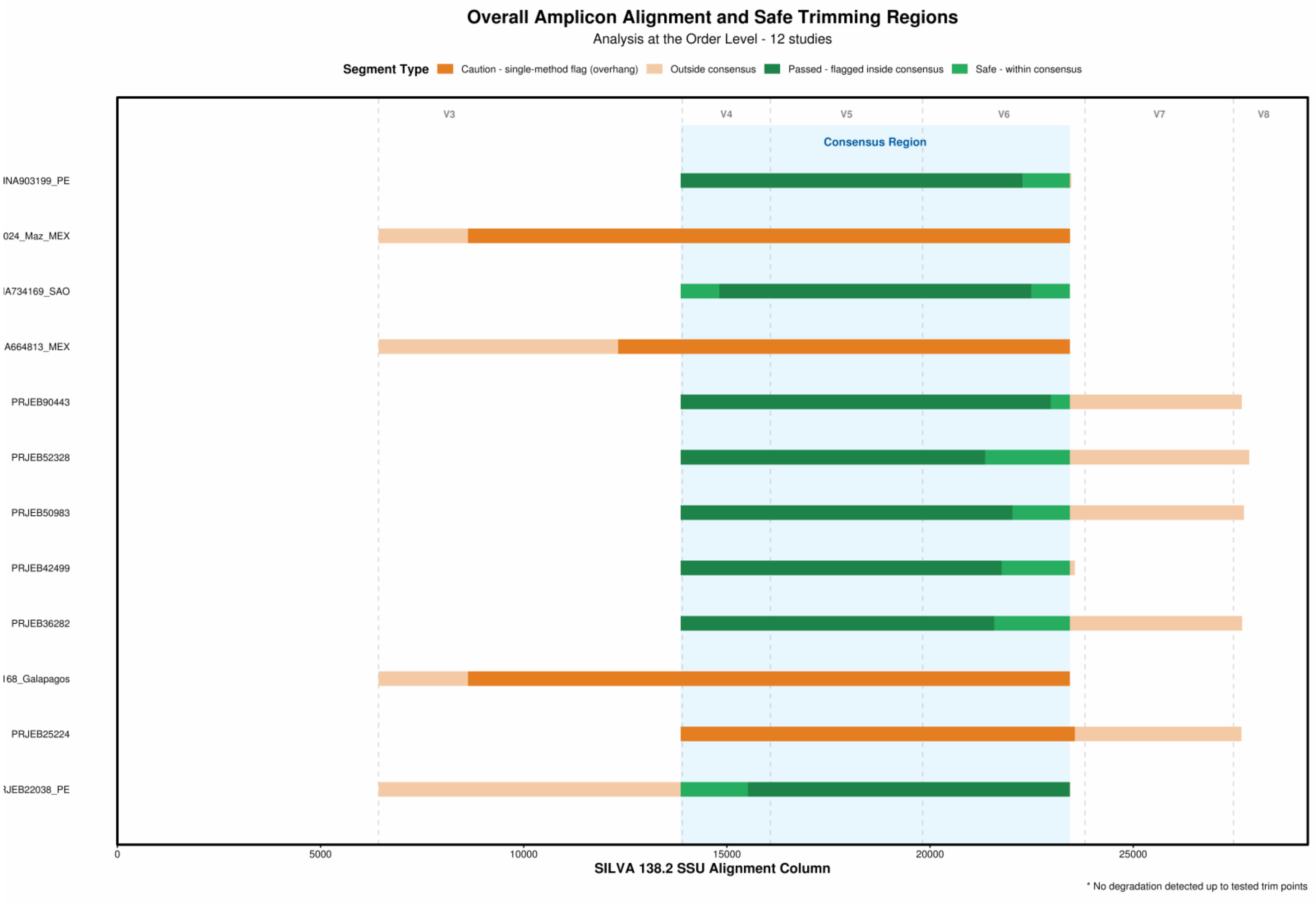
Overall Amplicon Alignment and Safe Trimming Regions
Analysis at the Order Level - 12 studies
Segment Type
Caution - single-method flag (overhang)
Outside consensus
Passed - flagged inside consensus
Safe - within consensus
V3
V4
V5
V6
V7
V8
Consensus Region
INA903199_PE
024_Maz_MEX
IA734169_SAO
A664813_MEX
PRJEB90443
PRJEB52328
PRJEB50983
PRJEB42499
PRJEB36282
168_Galapagos
PRJEB25224
RJEB22038_PE
0
5000
10000
15000
20000
25000
SILVA 138.2 SSU Alignment Column
* No degradation detected up to tested trim points

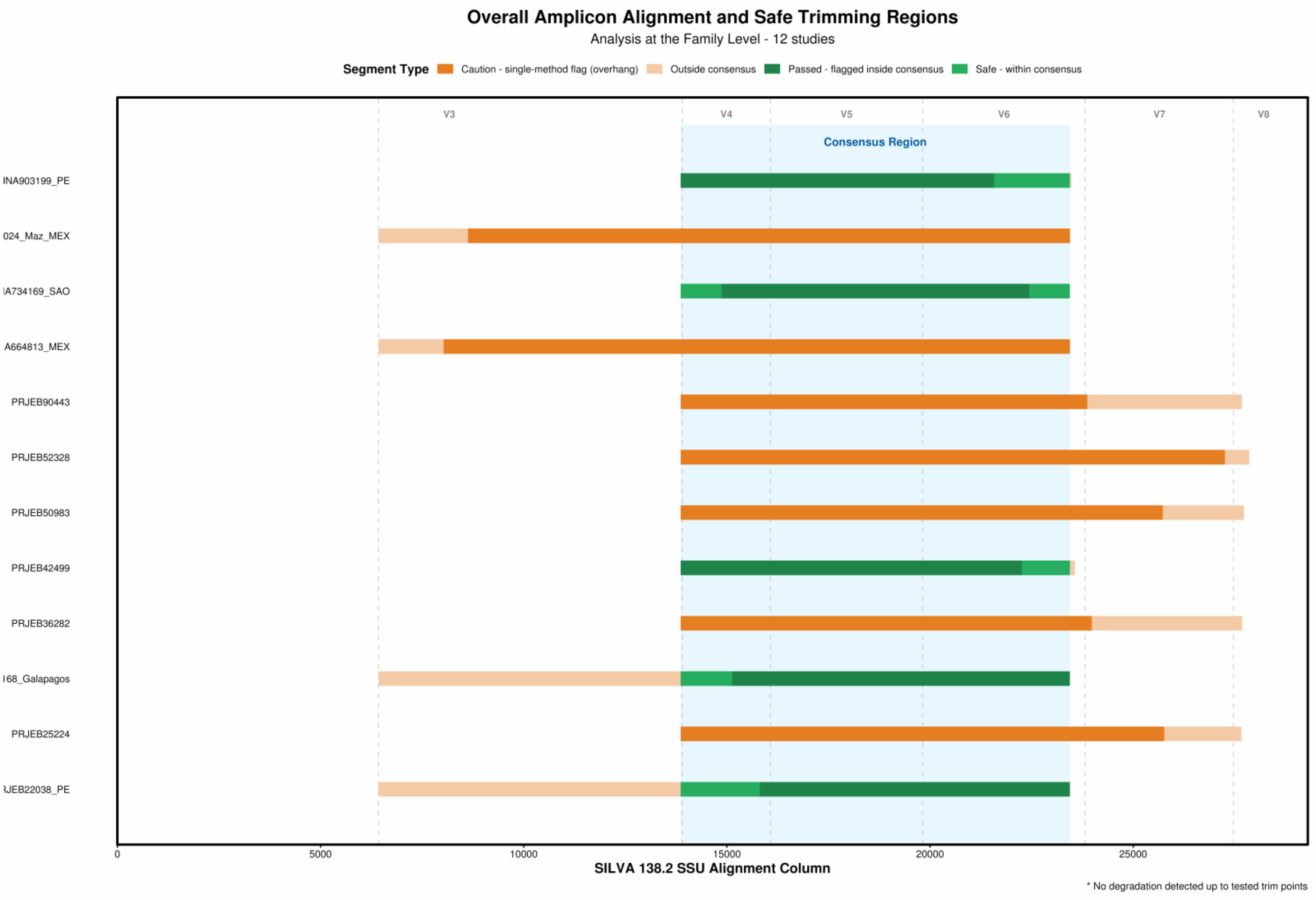

Overall Amplicon Alignment and Safe Trimming Regions
Analysis at the Family Level - 12 studies
Segment Type
Caution - single-method flag (overhang)
Outside consensus
Passed - flagged inside consensus
Safe - within consensus
V3
V4
V5
V6
V7
V8
Consensus Region
INA903199_PE
024_Maz_MEX
IA734169_SAO
A664813_MEX
PRJEB90443
PRJEB52328
PRJEB50983
PRJEB42499
PRJEB36282
168_Galapagos
PRJEB25224
JEB22038_PE
0
5000
10000
15000
20000
25000
SILVA 138.2 SSU Alignment Column
* No degradation detected up to tested trim points

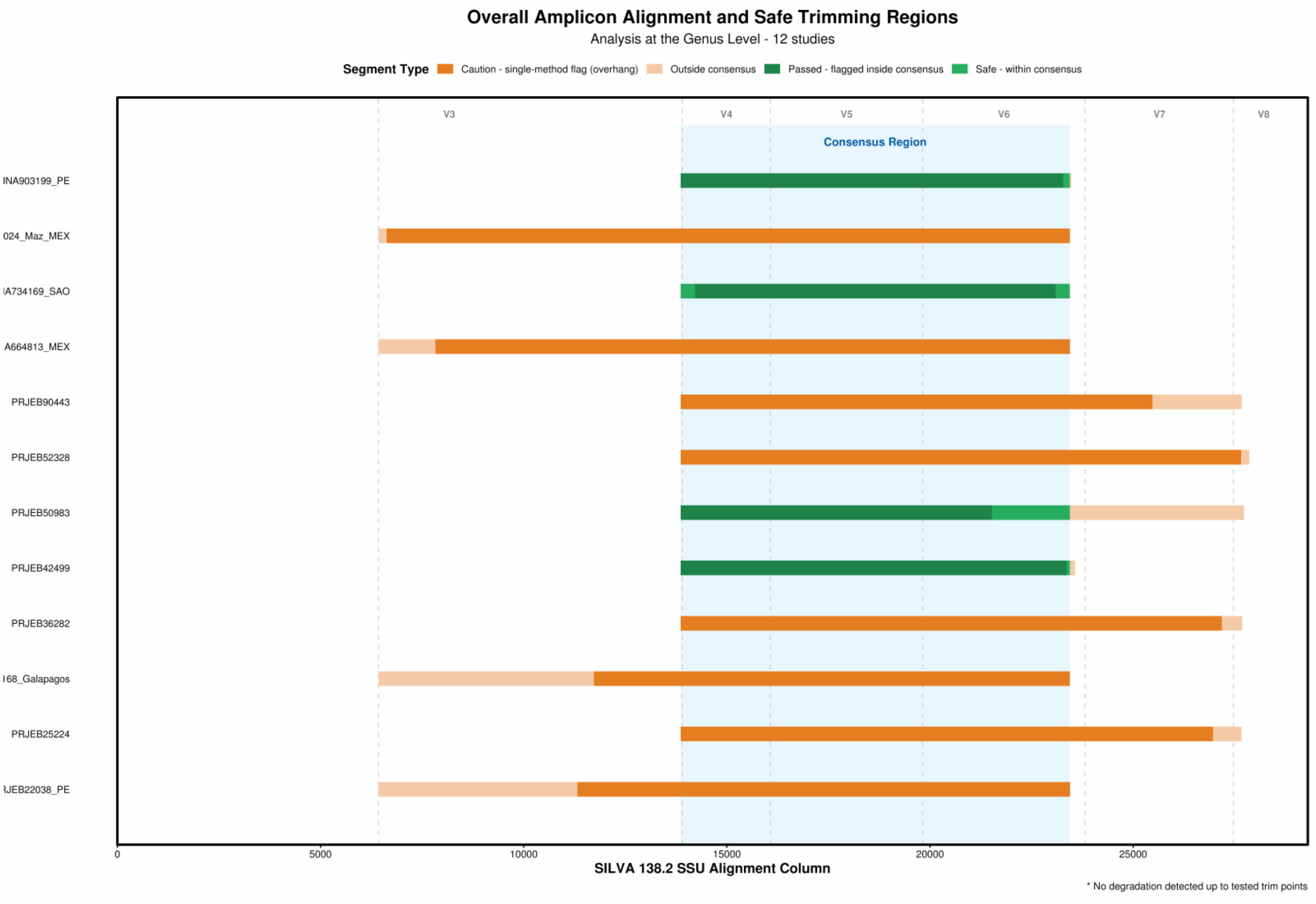
Overall Amplicon Alignment and Safe Trimming Regions
Analysis at the Genus Level - 12 studies
Segment Type
Caution - single-method flag (overhang)
Outside consensus
Passed - flagged inside consensus
Safe - within consensus
V3
V4
V5
V6
V7
V8
Consensus Region
INA903199_PE
024_Maz_MEX
IA734169_SAO
A664813_MEX
PRJEB90443
PRJEB52328
PRJEB50983
PRJEB42499
PRJEB36282
168_Galapagos
PRJEB25224
JEB22038_PE
0
5000
10000
15000
20000
25000
SILVA 138.2 SSU Alignment Column
* No degradation detected up to tested trim points

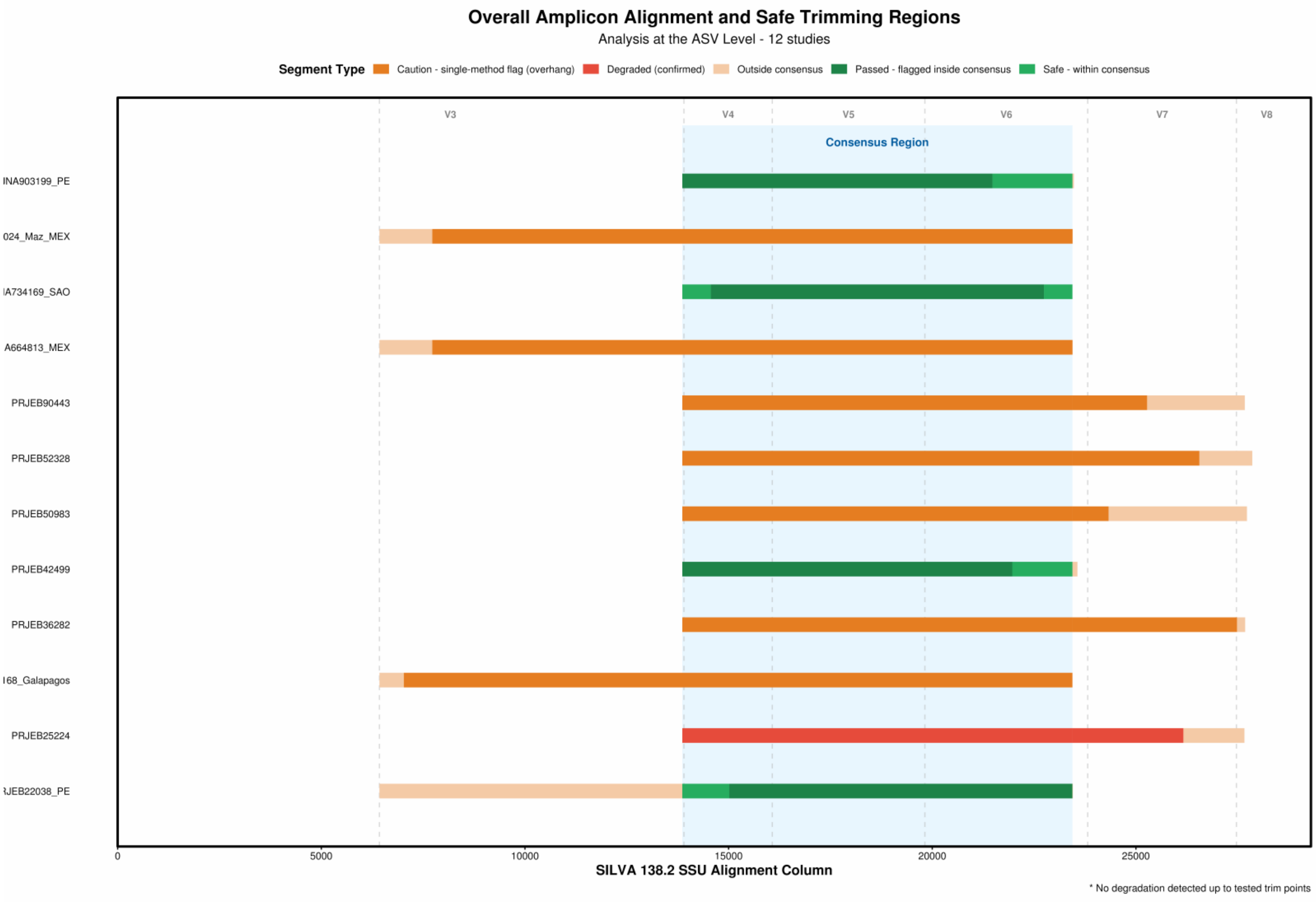
Overall Amplicon Alignment and Safe Trimming Regions
Analysis at the ASV Level - 12 studies
Segment Type
Caution - single-method flag (overhang)
Degraded (confirmed)
Outside consensus
Passed - flagged inside consensus
Safe - within consensus
V3
V4
V5
V6
V7
V8
Consensus Region
INA903199_PE
024_Maz_MEX
IA734169_SAO
A664813_MEX
PRJEB90443
PRJEB52328
PRJEB50983
PRJEB42499
PRJEB36282
I68_Galapagos
PRJEB25224
RJEB22038_PE
0
5000
10000
15000
20000
25000
SILVA 138.2 SSU Alignment Column
* No degradation detected up to tested trim points

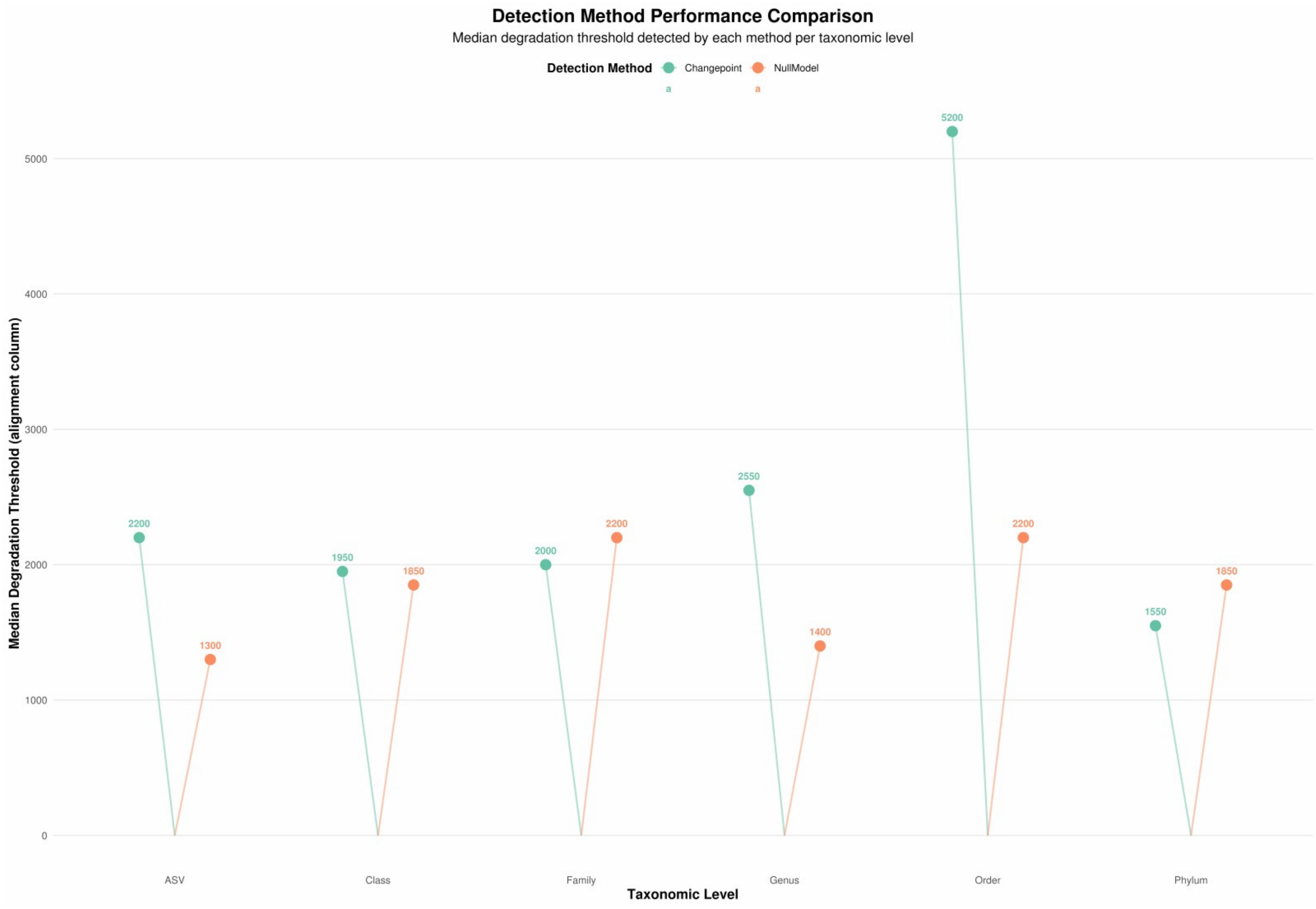




*Note on the reports in Supplementary Data S1 and S2. Both files are the unedited output of the pipeline's reporting module for the final run. Page headers and axis labels carry per-run dataset identifiers (for example PRJEB22038_PE), which correspond to the plain project accessions used in the text.*
