## Supplementary Note S1 for "SeCAT (Sequence Consensus Amplicon Trimming): a simulation-informed Nextflow pipeline for cross-study harmonisation of 16S rRNA metabarcoding datasets"

### Supplementary Note S1. Parameter calibration of the SeCAT degradation-detection ensemble.

SeCAT's statistical defaults are grounded in a systematic false-positive-rate (FPR) and sensitivity calibration study: FPR was measured under a no-signal null (simulated communities trimmed with no genuine degradation) and sensitivity under controlled-magnitude spike-ins (community shifts of known Bray-Curtis size), targeting Type I error control at alpha = 0.05 on the seagrass 16S meta-analysis benchmark set. Three detectors operate in combination, each with a distinct sensitivity/specificity profile that the defaults were tuned against; the conservative >= 2-of-3 voting ensemble is the load-bearing mechanism for FPR control.

The detector calibration was carried out with a distance cutoff of 0.15 and three consecutive null-model steps, and the sensitivity figures in Table S1 were obtained under that configuration. Under it the distance cutoff gave FPR ≈ 0.006 at a cutoff of 0.15 and FPR < 0.05 across the 0.10–0.30 range, which includes the shipped default of 0.20. The calibration reports no false-positive rate for two consecutive steps (its null-model result is FPR = 0 at p ≤ 0.01 and min_consecutive ≥ 3), and the shipped two-step setting is therefore more permissive than the calibrated three, with false-positive control in the demonstration run (voluminous_pauling, 17 September 2026, SeCAT v5.0.6) resting on the ≥ 2-of-3 ensemble agreement described below.

**Table S1. Calibrated behaviour of the three degradation detectors and the voting ensemble, from the calibration at a distance cutoff of 0.15 and three consecutive null-model steps.**

| Detector | Parameter (shipped default) | Calibrated behaviour (FPR / sensitivity) | Role in the ensemble |
|---|---|---|---|
| Null model | null_model_p_threshold = 0.05; null_model_min_consecutive = 2 | FPR ~ 0 across the parameter grid (FPR = 0 at p <= 0.01 and min_consecutive >= 3); near-zero standalone sensitivity | High-specificity confirmatory signal - almost never fires falsely |
| Distance cutoff | distance_cutoff_threshold = 0.20 | FPR ~ 0.006 at 0.15 and remains < 0.05 across the 0.10-0.30 range; fires at lag 0 (at the true signal); sensitivity approximately linear with spike size (~80% at delta = 0.3, ~5% at delta = 0.1) | Primary detector - well-calibrated workhorse of the ensemble |
| Changepoint (PELT) | Penalty selected for each study and taxonomic level by bootstrap cross-validation over a scan range of 0.4–2.4 (× log n); changepoint_penalty_multiplier is a legacy parameter that does not affect verdicts | In the calibration (fixed penalty multiplier): FPR ~ 0.70 at low multipliers; drops below 0.05 only near multiplier ~ 500, where sensitivity falls to ~15% for spike delta = 0.20 | Sensitivity-driven confirmatory signal - kept deliberately sensitive; relies on ensemble agreement for FPR control |
| Ensemble (>= 2 of 3 agree) | tolerance_bp = 500 (agreement window) | Load-bearing FPR-control mechanism: no single detector is FPR < 0.05 standalone, but the vote exploits the detectors' complementary error modes | Final KEEP / EXCLUDE gate; hard quality gates (bc_ceiling = 0.20, retention_floor = 0.70) override the vote when breached |

### Changepoint penalty selection

The changepoint (PELT) penalty is selected for each study and taxonomic level by bootstrap cross-validation over a scan range of 0.4–2.4 (× log n), in steps of 0.1 with 50 bootstrap resamples, and changepoint_penalty_multiplier is a legacy parameter that does not affect verdicts; in the demonstration run the selected penalties ranged from 0.4 to 2.3 (× log n) across the 72 study × level tests. The calibration, which varied a fixed penalty multiplier, showed that a penalty strict enough to control PELT's false-positive rate on its own (a multiplier of about 500) retains only ~15% sensitivity for moderate community shifts (delta Bray-Curtis = 0.20), which would defeat the detector's purpose as an early-warning signal, and the detector is therefore kept sensitive at the cost of standalone specificity, with the >= 2-methods-agree rule supplying the FPR control that the penalty alone cannot. In v5.0.6 the changepoint location is taken from bootstrap resamples that do not preserve trim order and is therefore unreliable; a corrected detector (ordered resampling with a final fit on the ordered series) is planned for v5.0.7.

**Table S2. Parameters most sensitive to the calibration, with shipped defaults.**

| Parameter | Shipped default | What it controls | Sensitivity |
|---|---|---|---|
| null_model_p_threshold | 0.05 | Alpha for the empirical null-model significance test | High - directly determines KEEP/EXCLUDE rate |
| null_model_min_consecutive | 2 | Consecutive significant trim steps required before a verdict triggers | Moderate - suppresses single-point false positives |
| changepoint_penalty_method | MANUAL | PELT penalty type | Low - MANUAL gives direct control |
| changepoint_scan_min / changepoint_scan_max / changepoint_bootstrap_n | 0.4 / 2.4 / 50 | Bootstrap cross-validation scan from which the PELT penalty (× log n) is selected for each study and taxonomic level; changepoint_penalty_multiplier is a legacy parameter that does not affect verdicts | Raising the range makes detection more conservative and lowering it flags smaller shifts; ensemble supplies FPR control |
| distance_cutoff_threshold | 0.20 | Bray-Curtis cutoff for the secondary distance-based verdict | High - stricter cutoffs flag more studies |
| num_simulations | 100 | Null-distribution replicates per study | Moderate - fewer replicates give noisier p-values |
| bc_ceiling | 0.20 | Hard Bray-Curtis ceiling that forces exclusion regardless of the vote | High - absolute safeguard against gradual drift |
| retention_floor | 0.70 | Minimum fraction of sequences retained after trimming | Moderate - guards against over-trimming |
| sim_pcr_gc_bias / sim_pcr_cycles / sim_error_rate | 0.65 / 25 / 0.003 | PCR- and error-model realism of the simulation null | Moderate - calibrated against typical Illumina MiSeq profiles |

### When deviation from the defaults is defensible

- Under-powered datasets (< 5 studies, < 30 samples per study): the null model's near-zero standalone sensitivity becomes a liability when the ensemble is degraded by missing votes -

consider relaxing null_model_p_threshold to 0.10 (null_model_min_consecutive is already 2 in the shipped defaults); the distance cutoff remains the safety net.

- Strongly heterogeneous primer sets (e.g. V1-V3 + V4 + V6-V8 in one analysis): reducing the study count entering the consensus is usually more defensible than relaxing the detectors, but the shipped distance_cutoff_threshold of 0.20 admits roughly 10-15 more studies in the benchmark set than the calibrated 0.15 without crossing FPR = 0.05.
- Publication-quality runs: increase num_simulations from 100 to 500 to tighten the null distribution; this does not change the changepoint penalty, which is selected from the data by bootstrap cross-validation rather than from the simulations.

Any deviation must be (i) recorded in the methods with its rationale, ideally with a sensitivity analysis re-run at the published default, and (ii) accompanied by the run provenance file output/final_outputs/provenance/params_used.yaml, which records every effective parameter for the run.

### Provenance

**Provenance of the values in this note.**

| Item | Source |
| --- | --- |
| Detector FPR / sensitivity figures (Table S1) | Manuscripts/SeCAT/0005-calibration-grounded-in-data.patch (calibration of 27 May 2026, run at a distance cutoff of 0.15 and three consecutive null-model steps; effective settings from Manuscripts/SeCAT/SeCAT_Methods_ratification.md) |
| Sensitive-parameter table + deviation guidance (Table S2) | Manuscripts/SeCAT/0004-parameter-calibration-section.patch |
| Shipped default values | SeCAT params.yaml (github.com/DETerrey/SeCAT, v5.0.6) |
| Effective parameters and selected changepoint penalties of the demonstration run (voluminous_pauling, 17 September 2026, SeCAT v5.0.6) | final_outputs/supporting/provenance/ params_used.yaml and final_outputs/3_verdicts/verdict_data_all_levels.csv in Thesis/Chapter2_SeCAT/manuscript/secat_paper_v506 .tar.gz |
| Hard-gate constants (bc_ceiling, retention_floor, tolerance_bp) | params_used.yaml of the demonstration run (bc_ceiling_threshold, retention_floor_threshold, verdict_consensus_tolerance_bp) |
