## Supplementary Data S2 for "SeCAT (Sequence Consensus Amplicon Trimming): a simulation-informed Nextflow pipeline for cross-study harmonisation of 16S rRNA metabarcoding datasets"

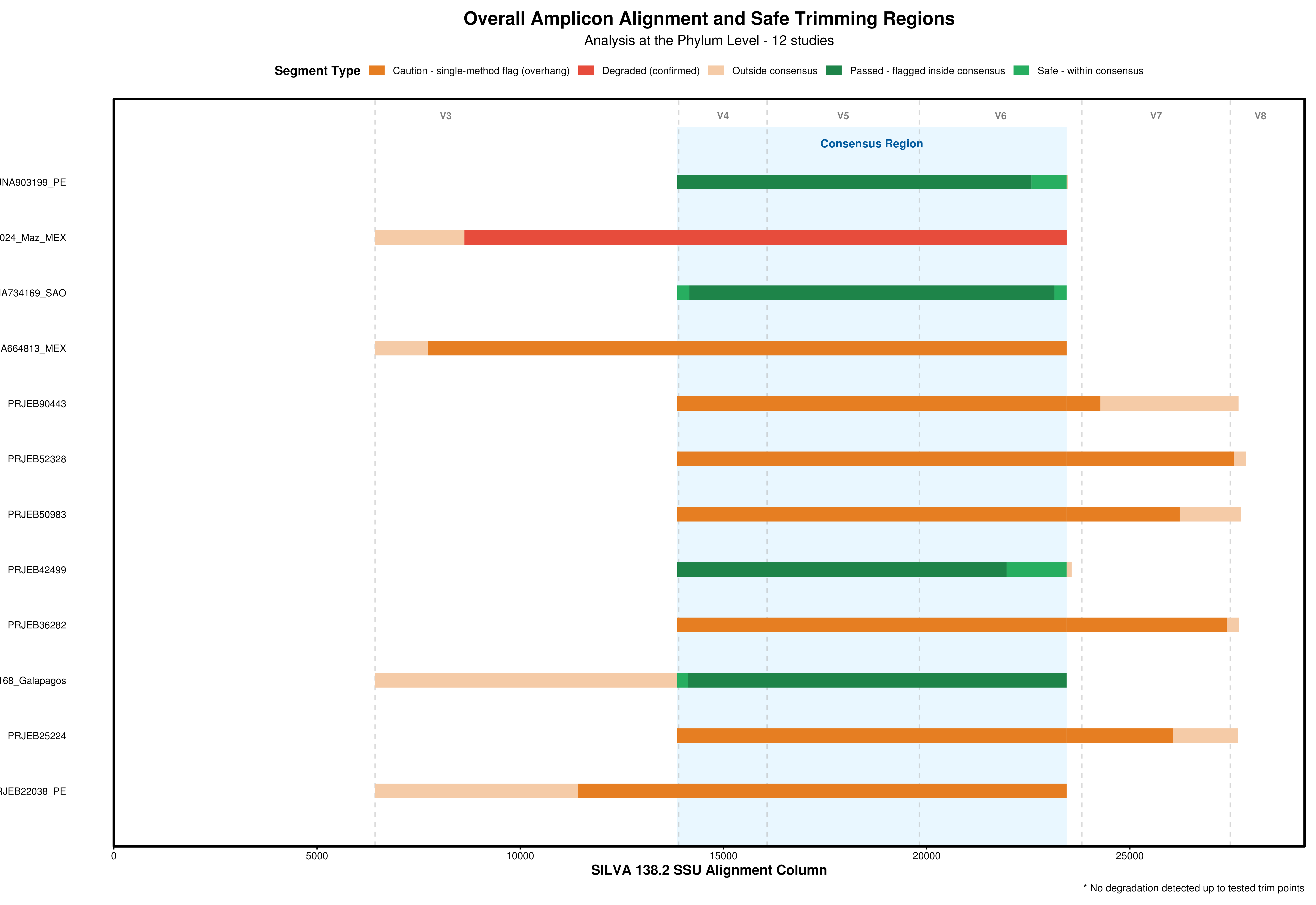

Overall Amplicon Alignment and Safe Trimming Regions
Analysis at the Phylum Level - 12 studies
Segment Type
Caution - single-method flag (overhang)
Degraded (confirmed)
Outside consensus
Passed - flagged inside consensus
Safe - within consensus
V3
V4
V5
V6
V7
V8
Consensus Region
NA903199_PE
024_Maz_MEX
A734169_SAO
A664813_MEX
PRJEB90443
PRJEB52328
PRJEB50983
PRJEB42499
PRJEB36282
168_Galapagos
PRJEB25224
RJEB22038_PE
0
5000
10000
15000
20000
25000
SILVA 138.2 SSU Alignment Column
* No degradation detected up to tested trim points

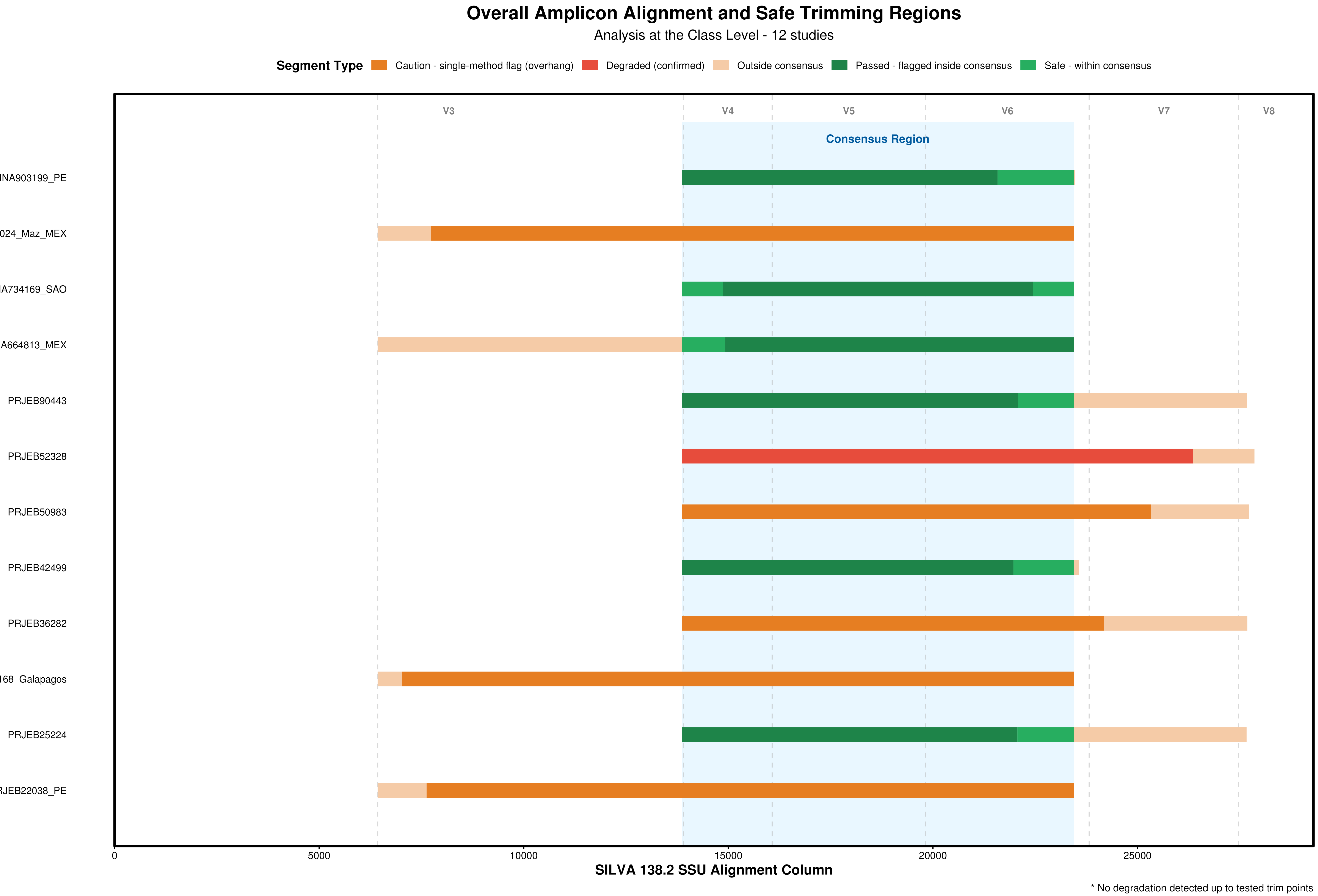
Overall Amplicon Alignment and Safe Trimming Regions
Analysis at the Class Level - 12 studies
Segment Type
Caution - single-method flag (overhang)
Degraded (confirmed)
Outside consensus
Passed - flagged inside consensus
Safe - within consensus
V3
V4
V5
V6
V7
V8
Consensus Region
JNA903199_PE
024_Maz_MEX
IA734169_SAO
A664813_MEX
PRJEB90443
PRJEB52328
PRJEB50983
PRJEB42499
PRJEB36282
168_Galapagos
PRJEB25224
RJEB22038_PE
0
5000
10000
15000
20000
25000
SILVA 138.2 SSU Alignment Column
* No degradation detected up to tested trim points

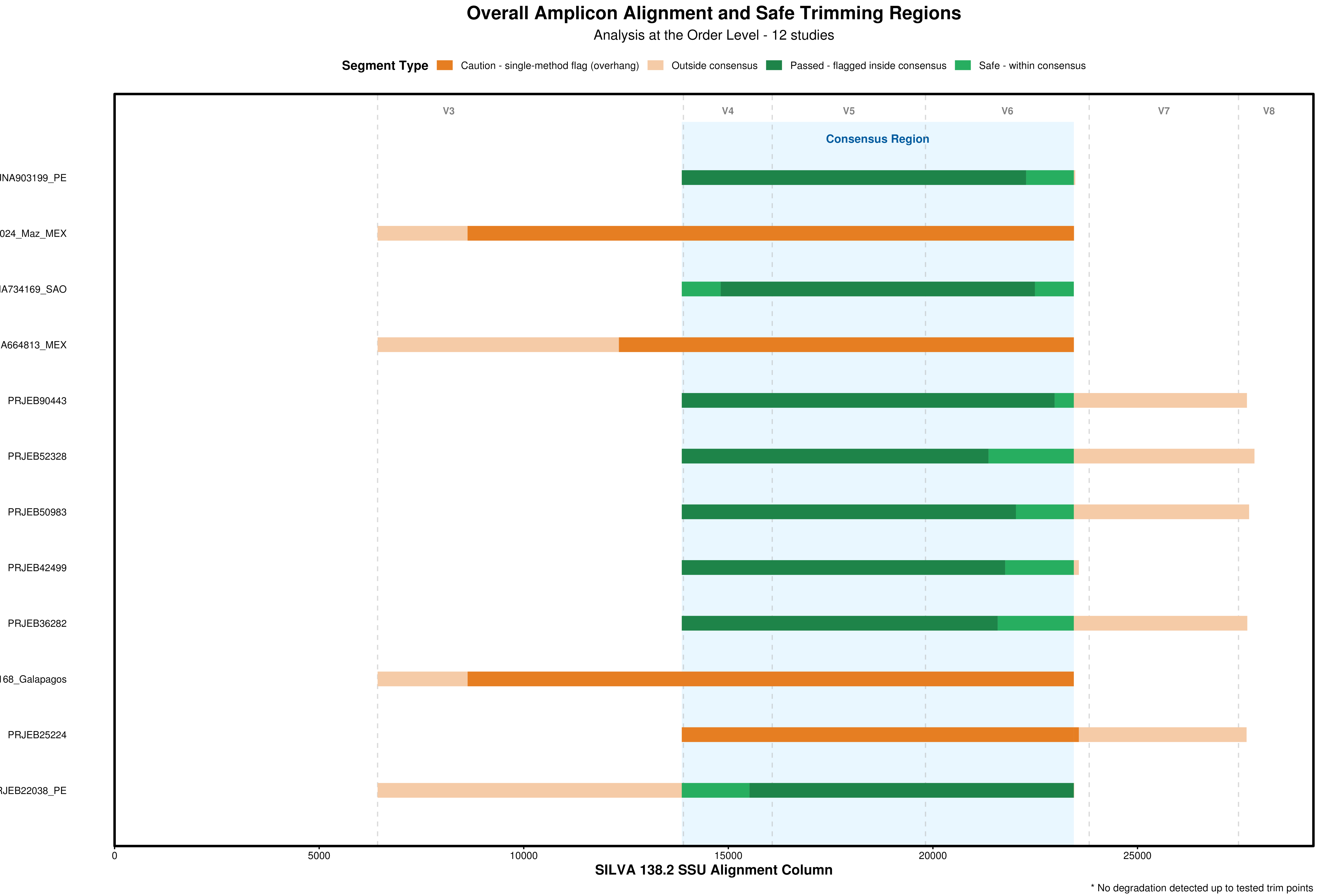
Overall Amplicon Alignment and Safe Trimming Regions
Analysis at the Order Level - 12 studies
Segment Type
Caution - single-method flag (overhang)
Outside consensus
Passed - flagged inside consensus
Safe - within consensus
V3
V4
V5
V6
V7
V8
Consensus Region
NA903199_PE
024_Maz_MEX
A734169_SAO
A664813_MEX
PRJEB90443
PRJEB52328
PRJEB50983
PRJEB42499
PRJEB36282
168_Galapagos
PRJEB25224
RJEB22038_PE
0
5000
10000
15000
20000
25000
SILVA 138.2 SSU Alignment Column
* No degradation detected up to tested trim points

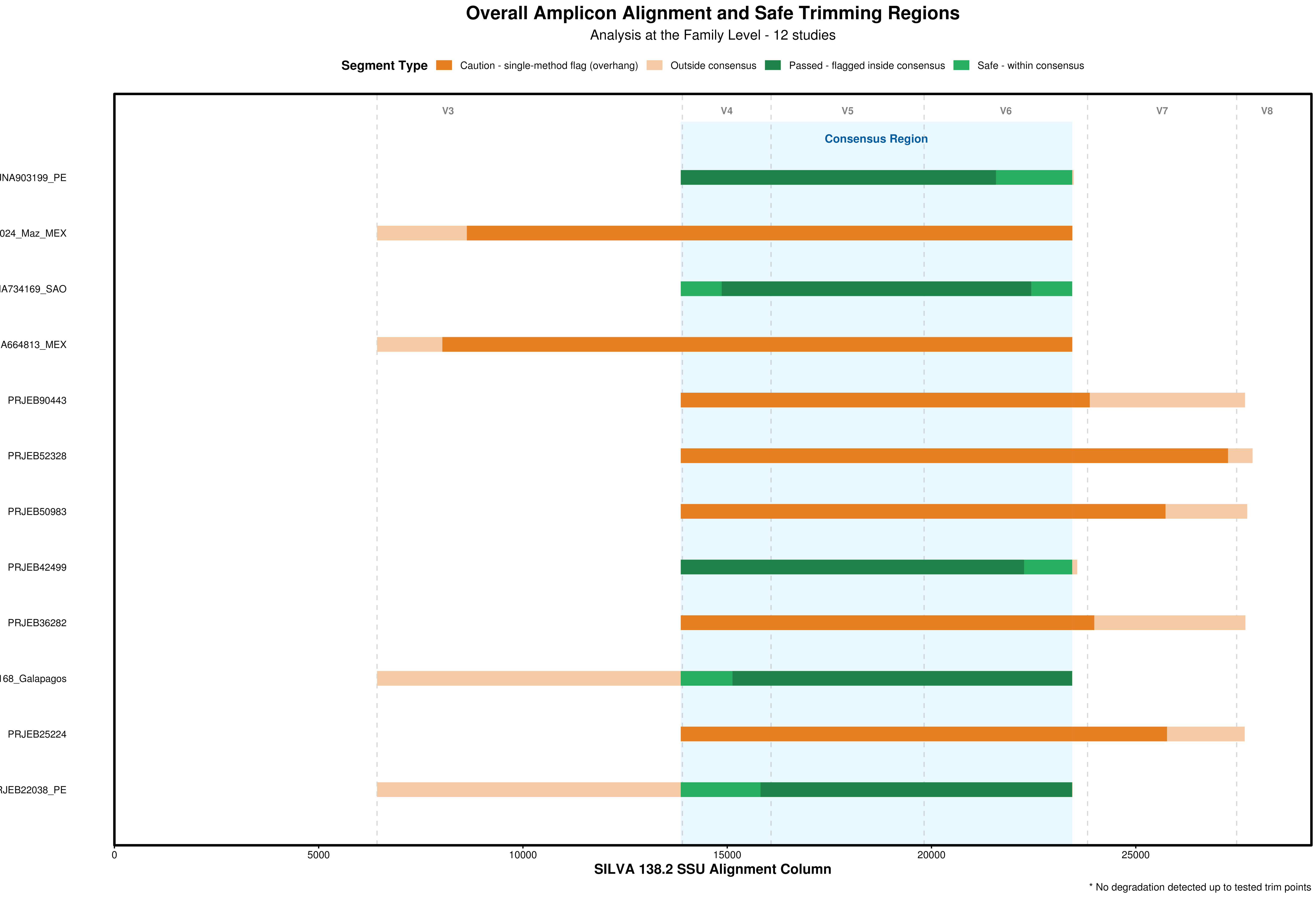
Overall Amplicon Alignment and Safe Trimming Regions
Analysis at the Family Level - 12 studies
Segment Type
Caution - single-method flag (overhang)
Outside consensus
Passed - flagged inside consensus
Safe - within consensus
V3
V4
V5
V6
V7
V8
Consensus Region
NA903199_PE
024_Maz_MEX
A734169_SAO
A664813_MEX
PRJEB90443
PRJEB52328
PRJEB50983
PRJEB42499
PRJEB36282
68_Galapagos
PRJEB25224
JEB22038_PE
0
5000
10000
15000
20000
25000
SILVA 138.2 SSU Alignment Column
* No degradation detected up to tested trim points

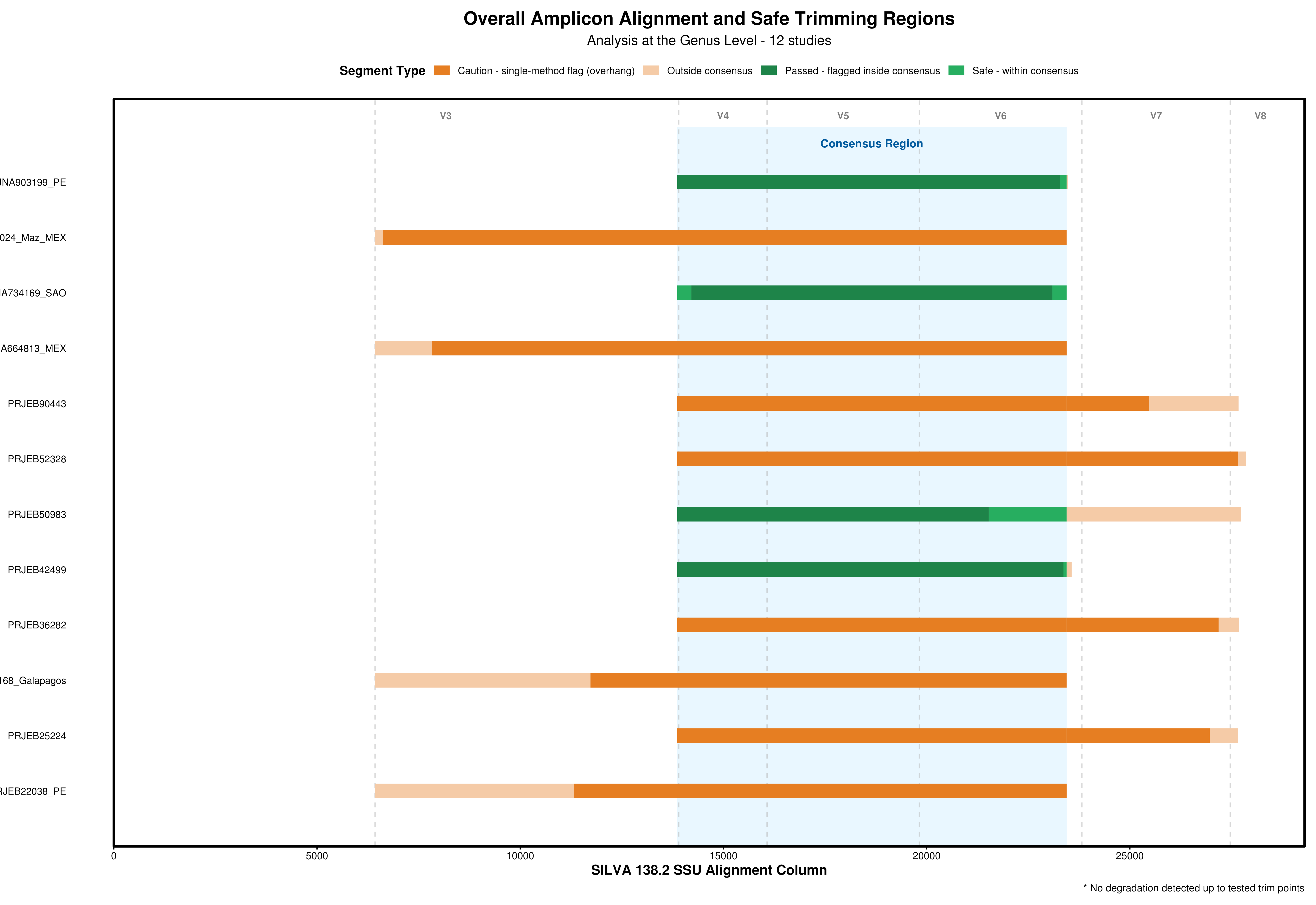
Overall Amplicon Alignment and Safe Trimming Regions
Analysis at the Genus Level - 12 studies
Segment Type
Caution - single-method flag (overhang)
Outside consensus
Passed - flagged inside consensus
Safe - within consensus
V3
V4
V5
V6
V7
V8
Consensus Region
NA903199_PE
024_Maz_MEX
IA734169_SAO
A664813_MEX
PRJEB90443
PRJEB52328
PRJEB50983
PRJEB42499
PRJEB36282
168_Galapagos
PRJEB25224
RJEB22038_PE
0
5000
10000
15000
20000
25000
SILVA 138.2 SSU Alignment Column
* No degradation detected up to tested trim points

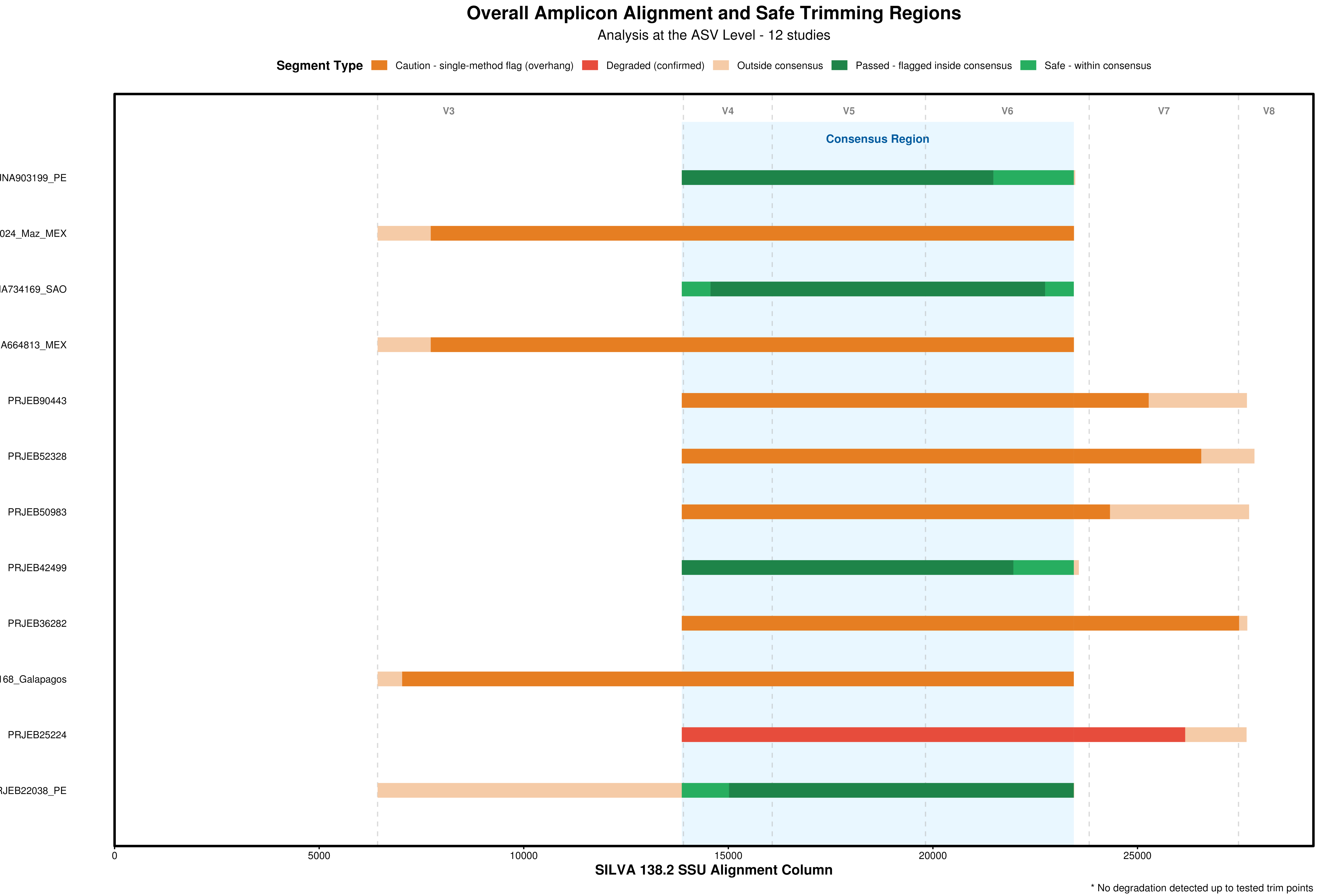

Overall Amplicon Alignment and Safe Trimming Regions
Analysis at the ASV Level - 12 studies
Segment Type
Caution - single-method flag (overhang)
Degraded (confirmed)
Outside consensus
Passed - flagged inside consensus
Safe - within consensus
V3
V4
V5
V6
V7
V8
Consensus Region
JNA903199_PE
024_Maz_MEX
JA734169_SAO
A664813_MEX
PRJEB90443
PRJEB52328
PRJEB50983
PRJEB42499
PRJEB36282
168_Galapagos
PRJEB25224
RJEB22038_PE
0
5000
10000
15000
20000
25000
SILVA 138.2 SSU Alignment Column
* No degradation detected up to tested trim points

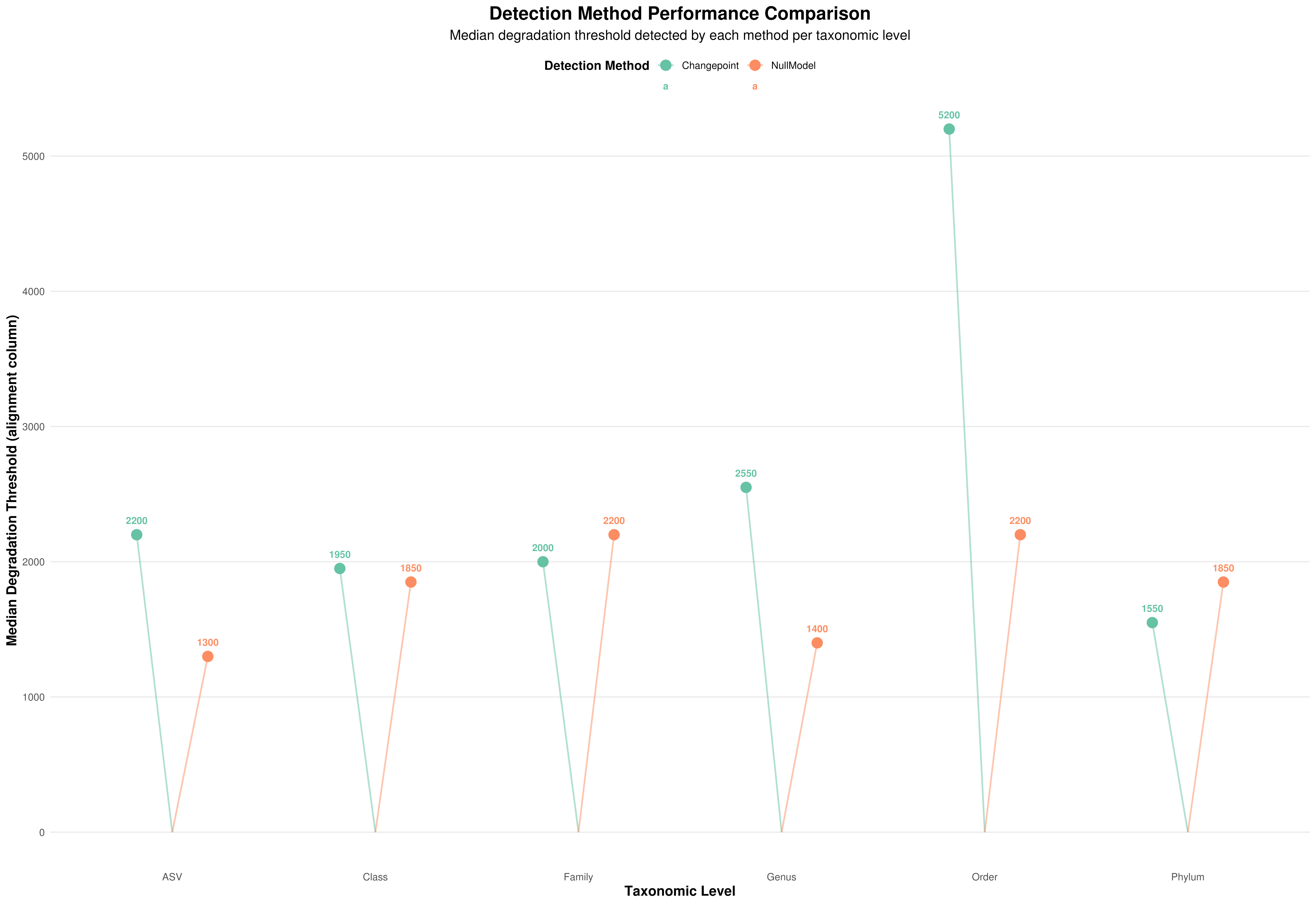

Detection Method Performance Comparison
Median degradation threshold detected by each method per taxonomic level
Detection Method
Changepoint
NullModel
a
a
5200
2550
2200
1300
1950
1850
2000
2200
1400
2200
1550
1850
0
1000
2000
3000
4000
5000
Median Degradation Threshold (alignment column)
ASV
Class
Family
Genus
Order
Phylum
Taxonomic Level
