## Supplementary Data S1 for "SeCAT (Sequence Consensus Amplicon Trimming): a simulation-informed Nextflow pipeline for cross-study harmonisation of 16S rRNA metabarcoding datasets"

### SeCAT Analysis Report: PRJEB22038_PE

Level: ASV | Method: Warning ... Changepoint only (1/1 methods) | Observed: 8600 bp (1-method warning) | Verdict: CAUTION

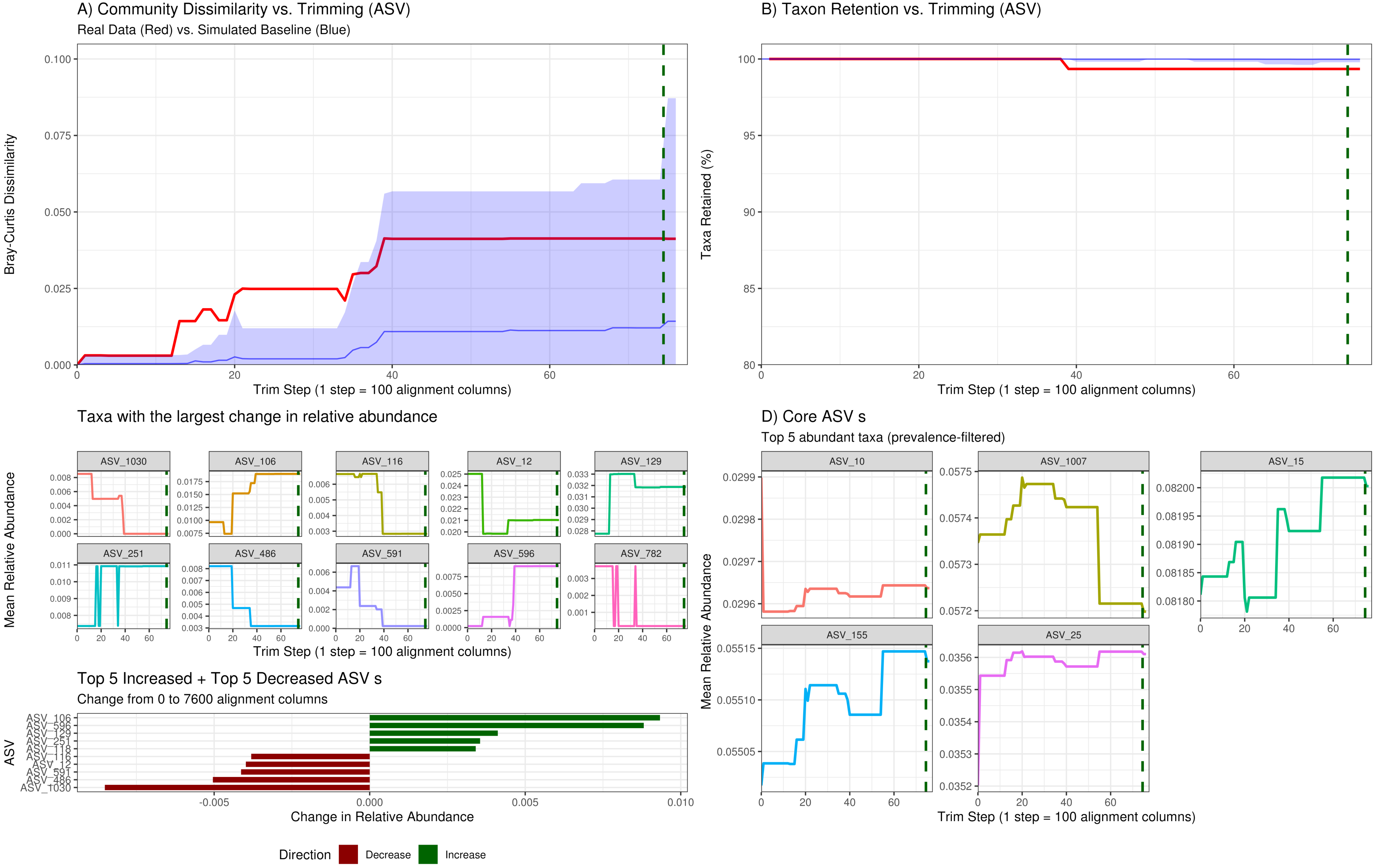

### SeCAT Analysis Report: PRJEB27168_Galapagos

Level: ASV | Method: Warning ... Changepoint only (1/1 methods) | Observed: 600 bp (1-method warning) | Verdict: CAUTION

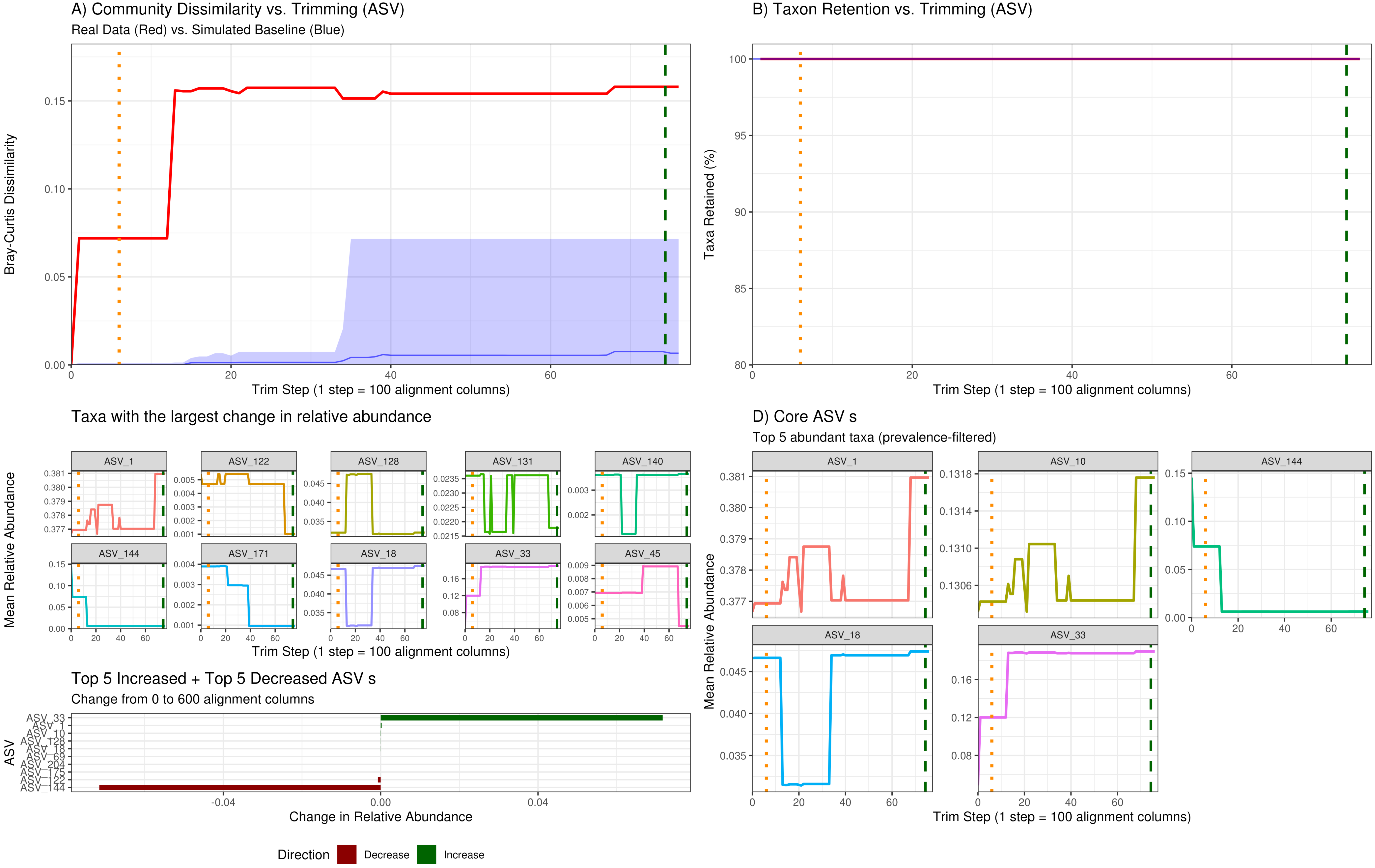


Generated: 2026-09-17 | Primer: 341F_805R | Valid data to 95 alignment columns

### SeCAT Analysis Report: PRJNA664813_MEX

Level: ASV | Method: Warning ... Changepoint only (1/2 methods) | Observed: 1300 bp (1-method warning) | Verdict: CAUTION

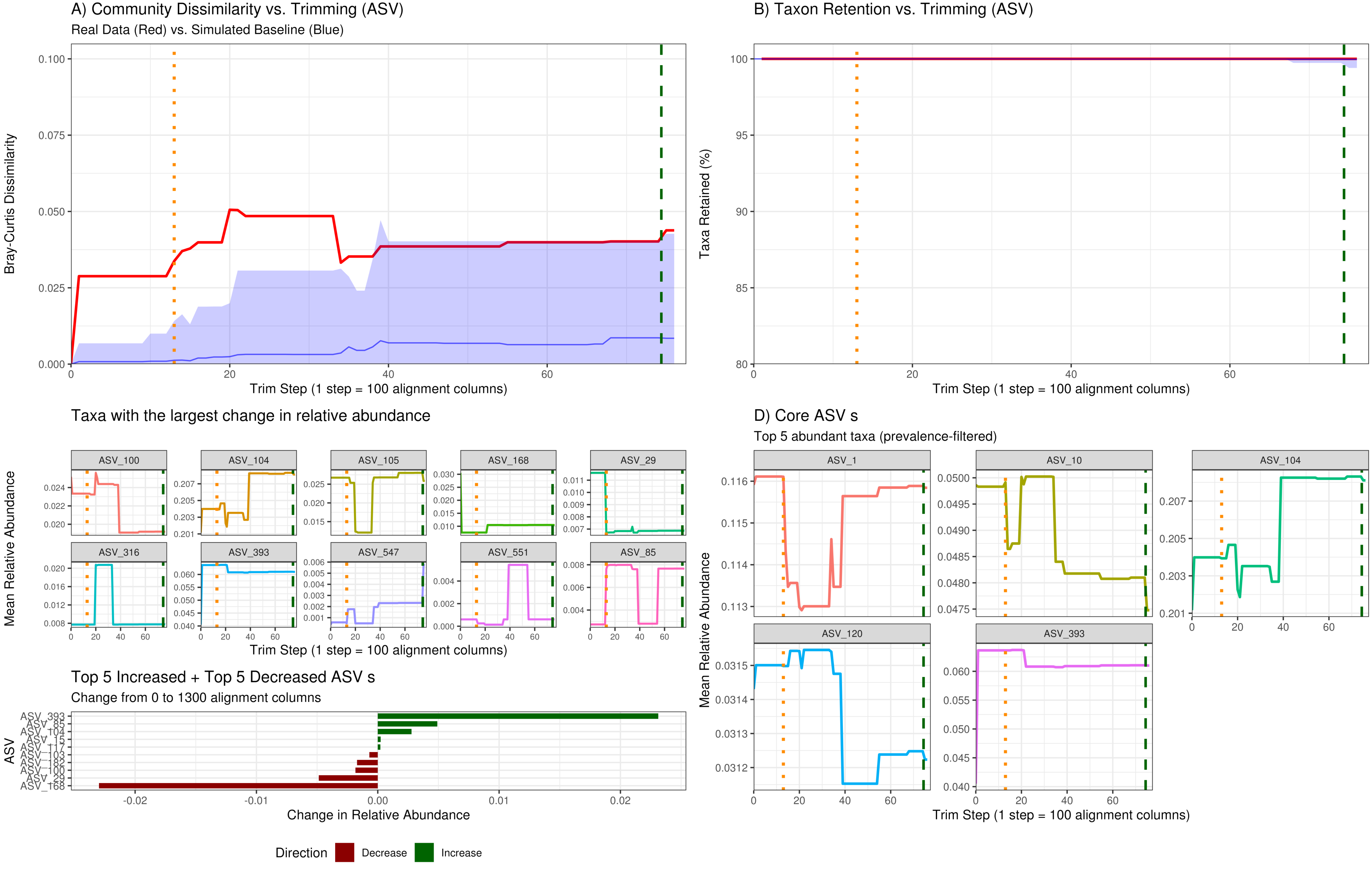


Generated: 2026-09-17 | Primer: 341F_805R | Valid data to 95 alignment columns

### SeCAT Analysis Report: PRJNA780024_Maz_MEX

Level: ASV | Method: Warning ... Changepoint only (1/2 methods) | Observed: 1300 bp (1-method warning) | Verdict: CAUTION

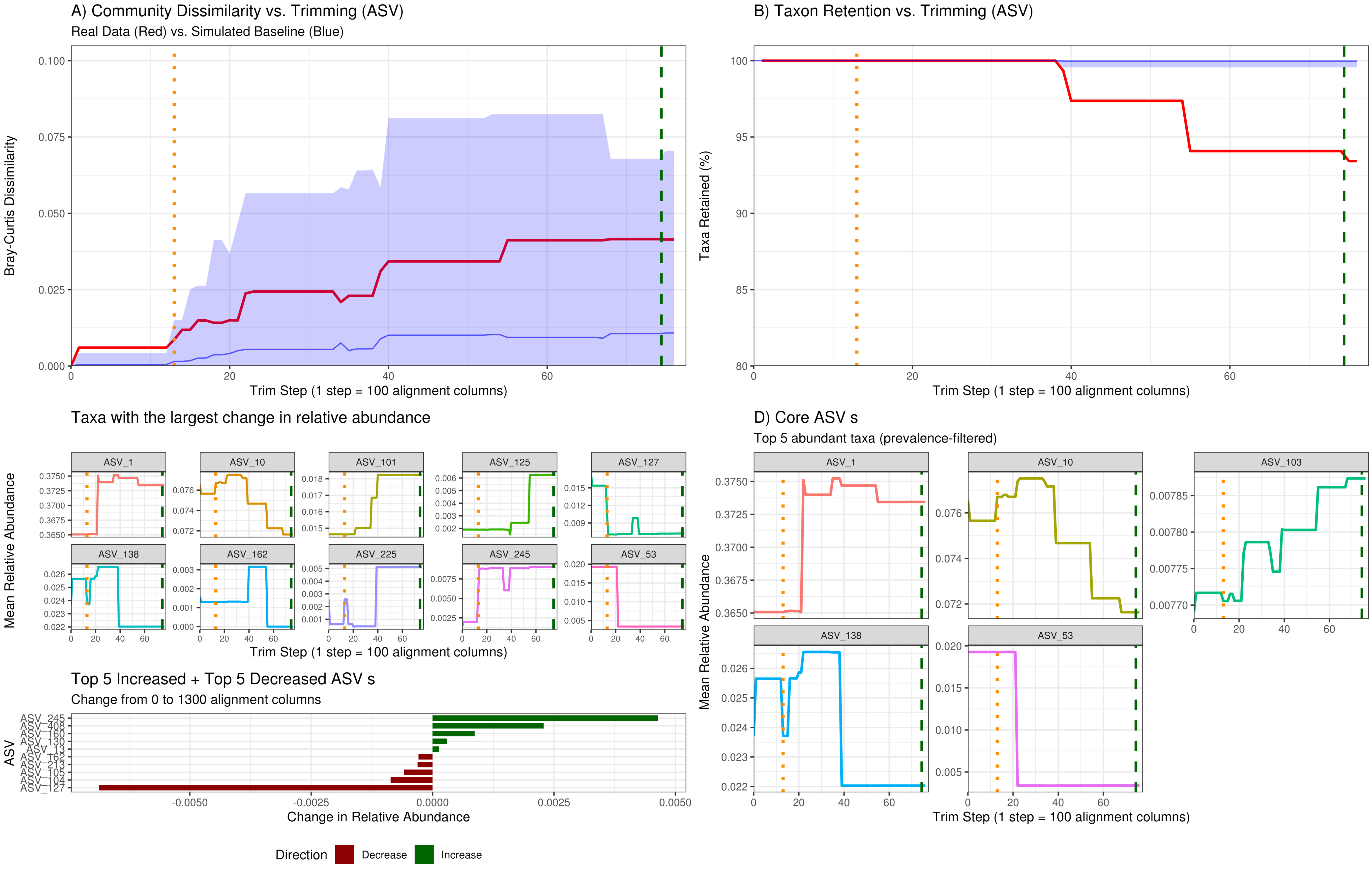


Generated: 2026-09-17 | Primer: 341F_805R | Valid data to 95 alignment columns

### SeCAT Analysis Report: PRJEB25224

Level: ASV | Method: Consensus (2 methods agree) | Observed: 1700 alignment columns | Verdict: FAIL

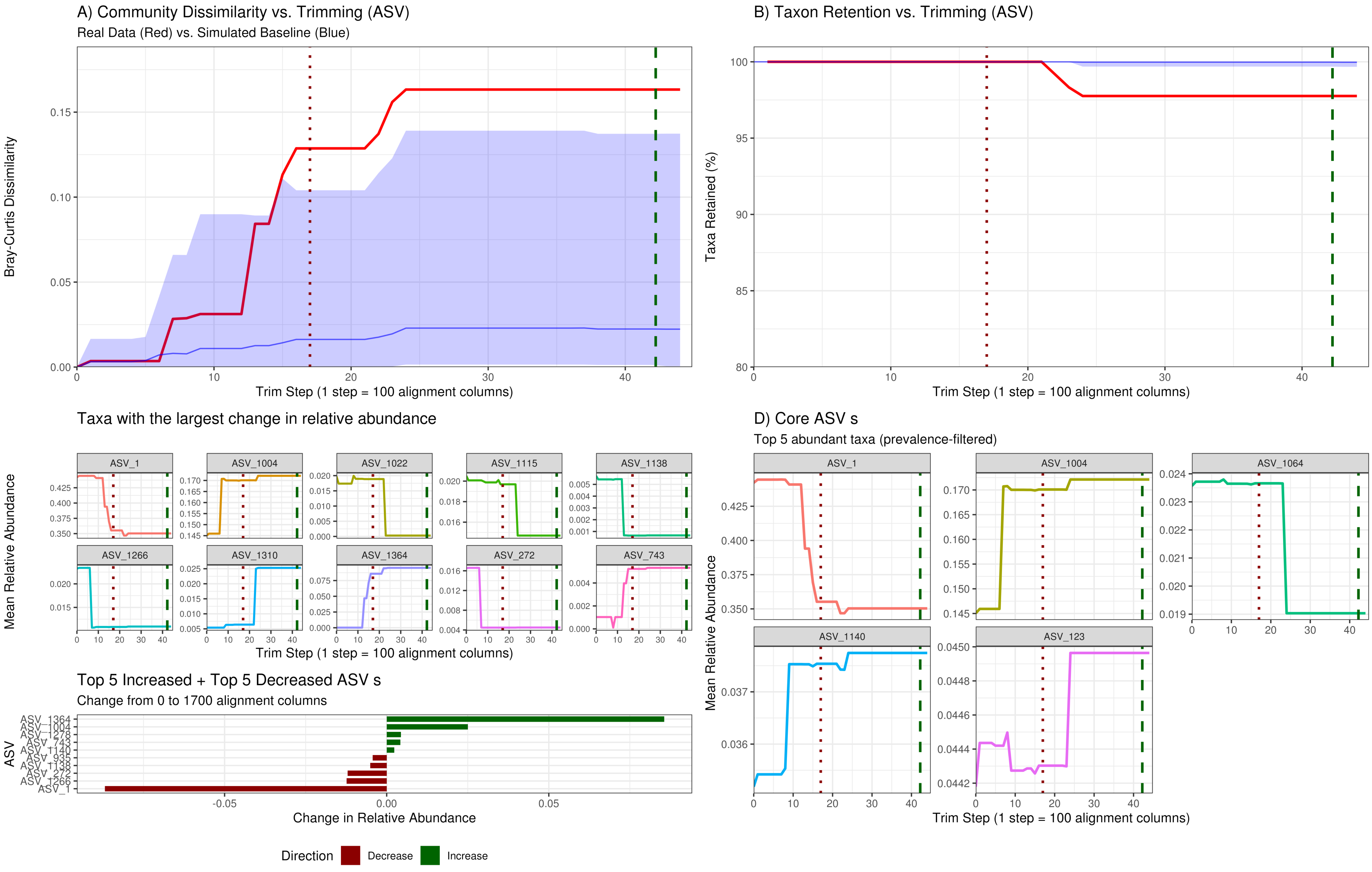


Generated: 2026-09-17 | Primer: 515F-Y_926R | Valid data to 63 alignment columns

### SeCAT Analysis Report: PRJEB36282

Level: ASV | Method: Warning ... Changepoint only (1/2 methods) | Observed: 200 bp (1-method warning) | Verdict: CAUTION

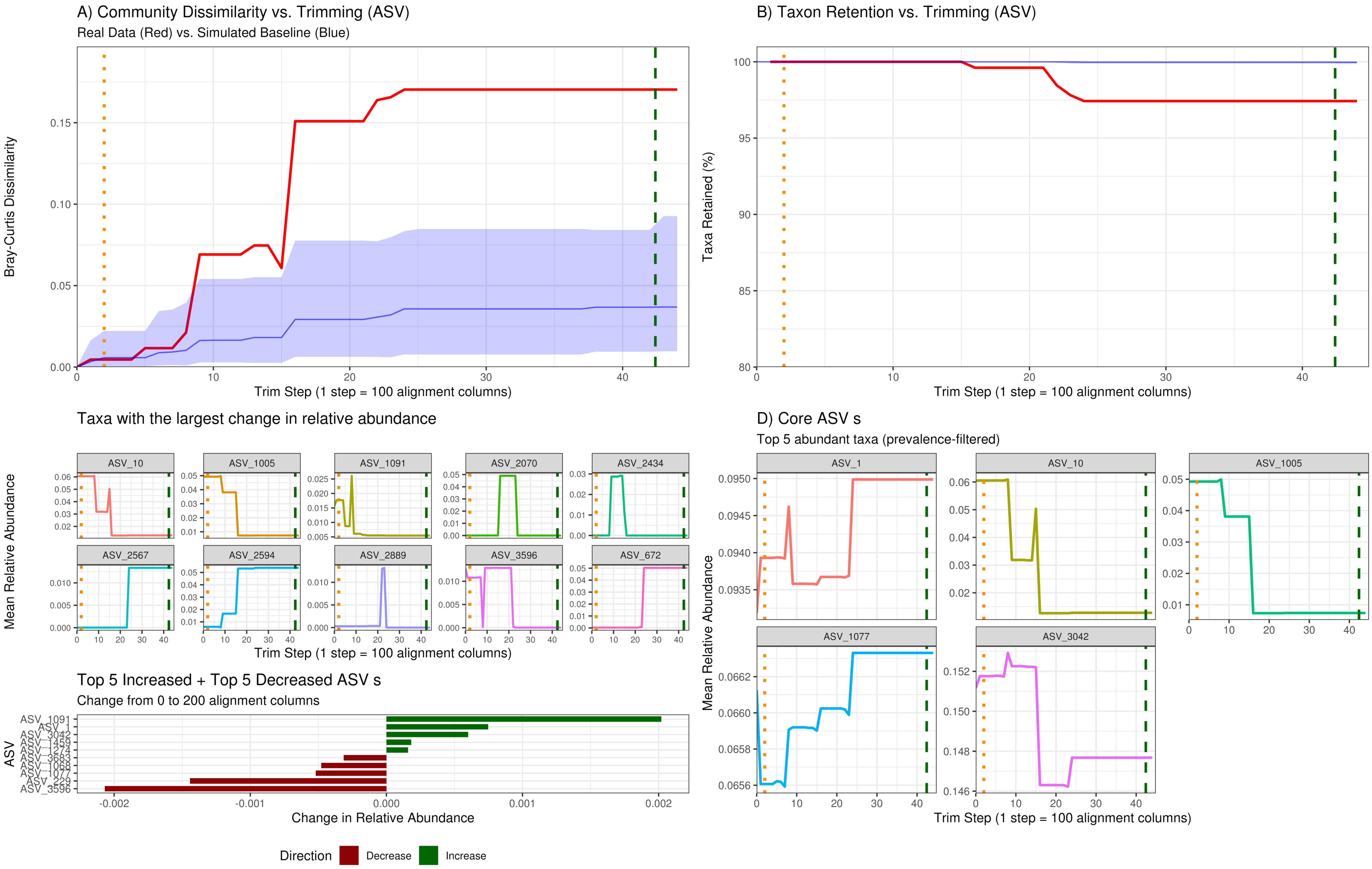


Generated: 2026-09-17 | Primer: 515F-Y_926R | Valid data to 63 alignment columns

### SeCAT Analysis Report: PRJEB50983

Level: ASV | Method: Warning ... Changepoint only (1/1 methods) | Observed: 3400 bp (1-method warning) | Verdict: CAUTION

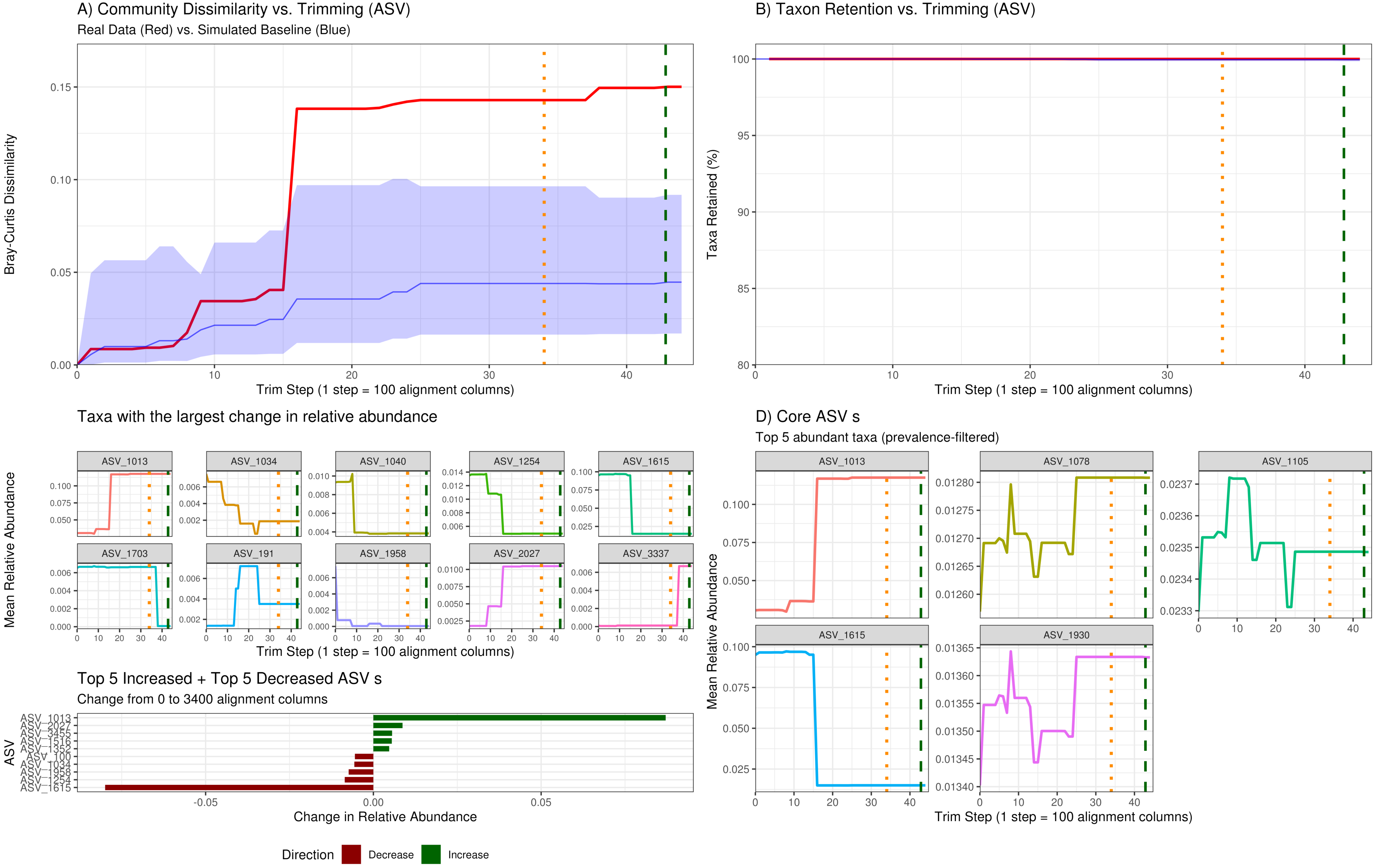


Generated: 2026-09-17 | Primer: 515F-Y_926R | Valid data to 63 alignment columns

### SeCAT Analysis Report: PRJEB52328

Level: ASV | Method: Warning ... Changepoint only (1/2 methods) | Observed: 1300 bp (1-method warning) | Verdict: CAUTION

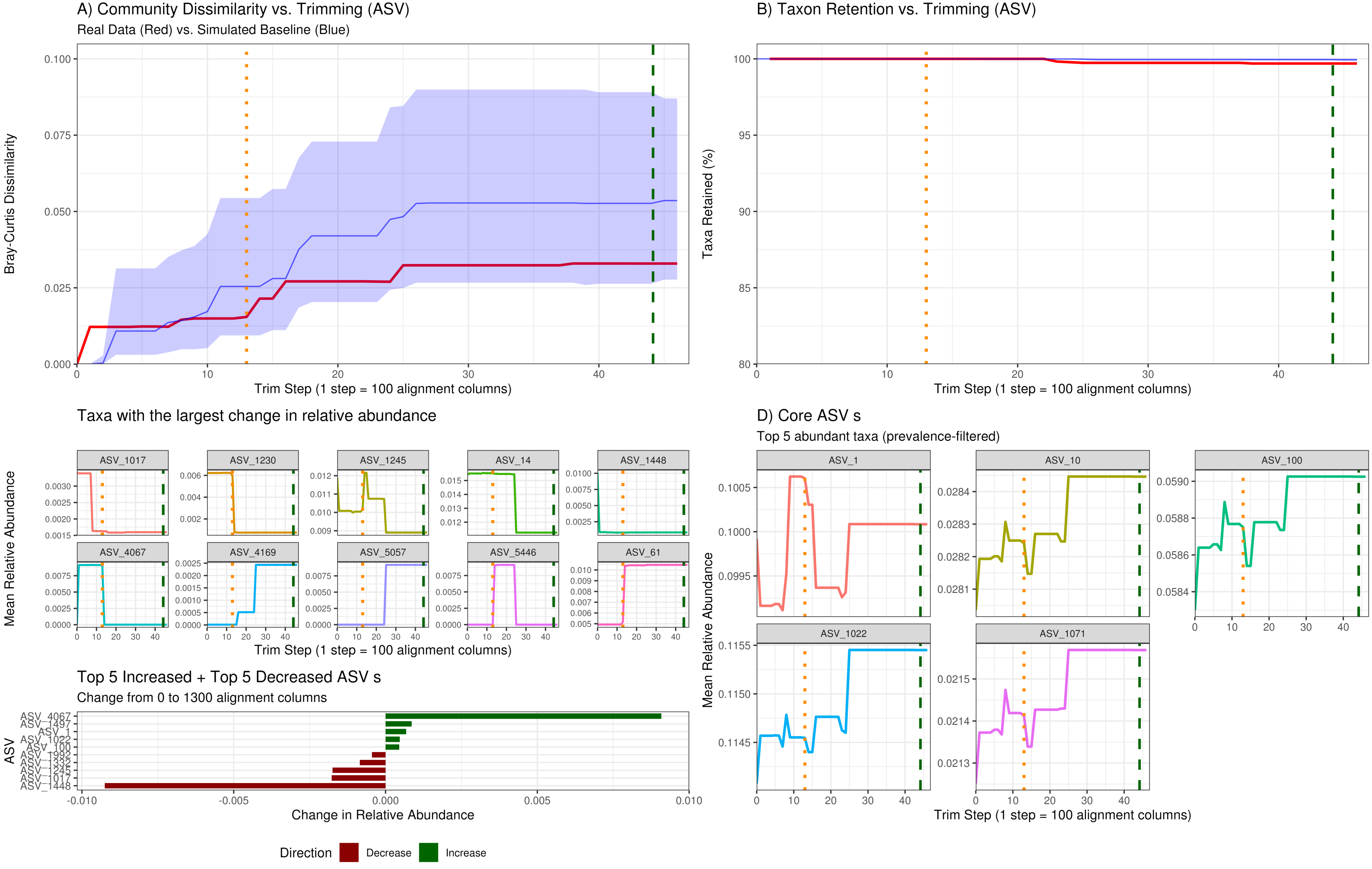


Generated: 2026-09-17 | Primer: 515F-Y_926R | Valid data to 65 alignment columns

### SeCAT Analysis Report: PRJEB90443

Level: ASV | Method: Warning ... Changepoint only (1/2 methods) | Observed: 2400 bp (1-method warning) | Verdict: CAUTION

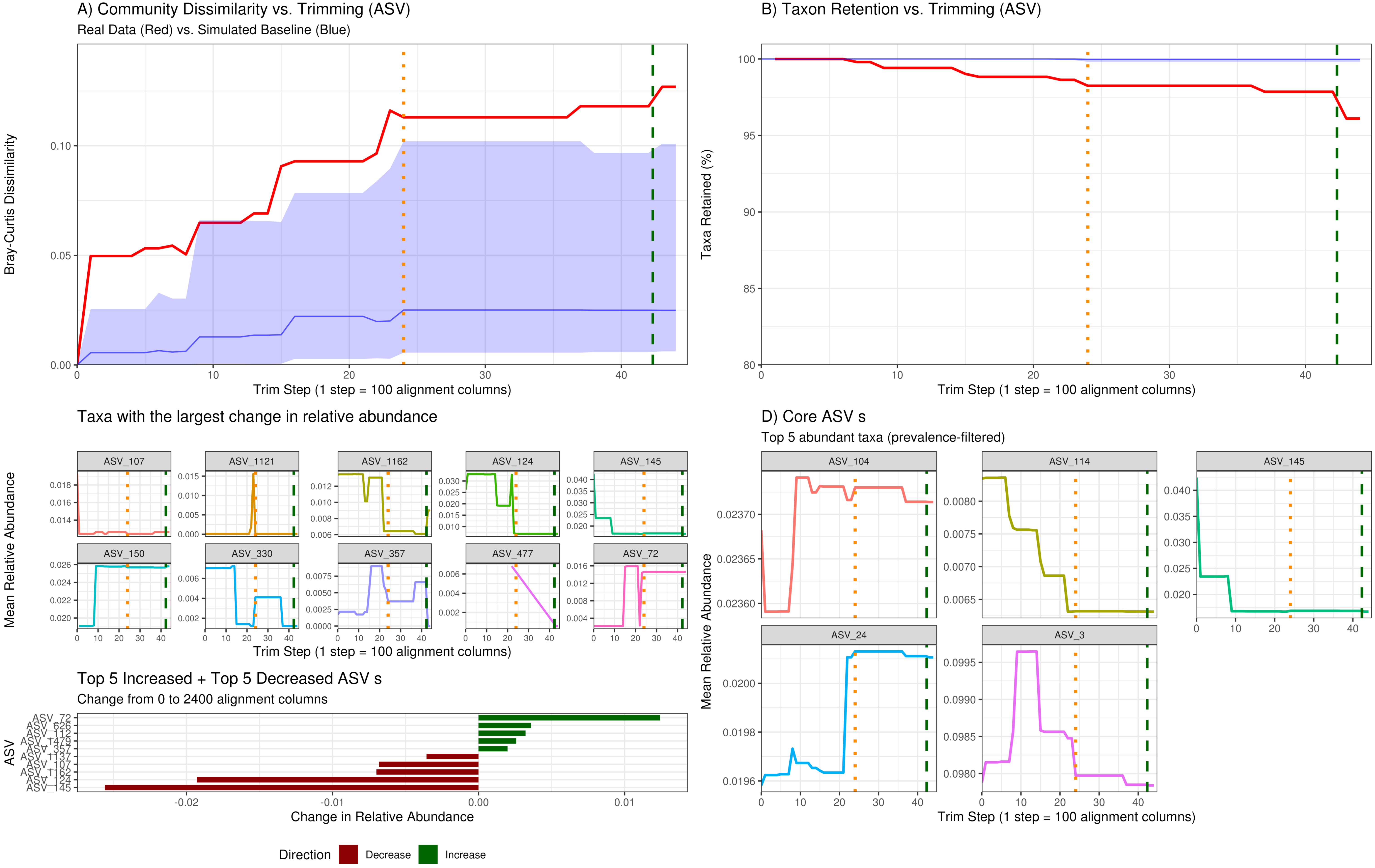


Generated: 2026-09-17 | Primer: 515F-Y_926R | Valid data to 63 alignment columns

### SeCAT Analysis Report: PRJEB42499

Level: ASV | Method: Warning ... Changepoint only (1/1 methods) | Observed: 1600 bp (1-method warning) | Verdict: CAUTION

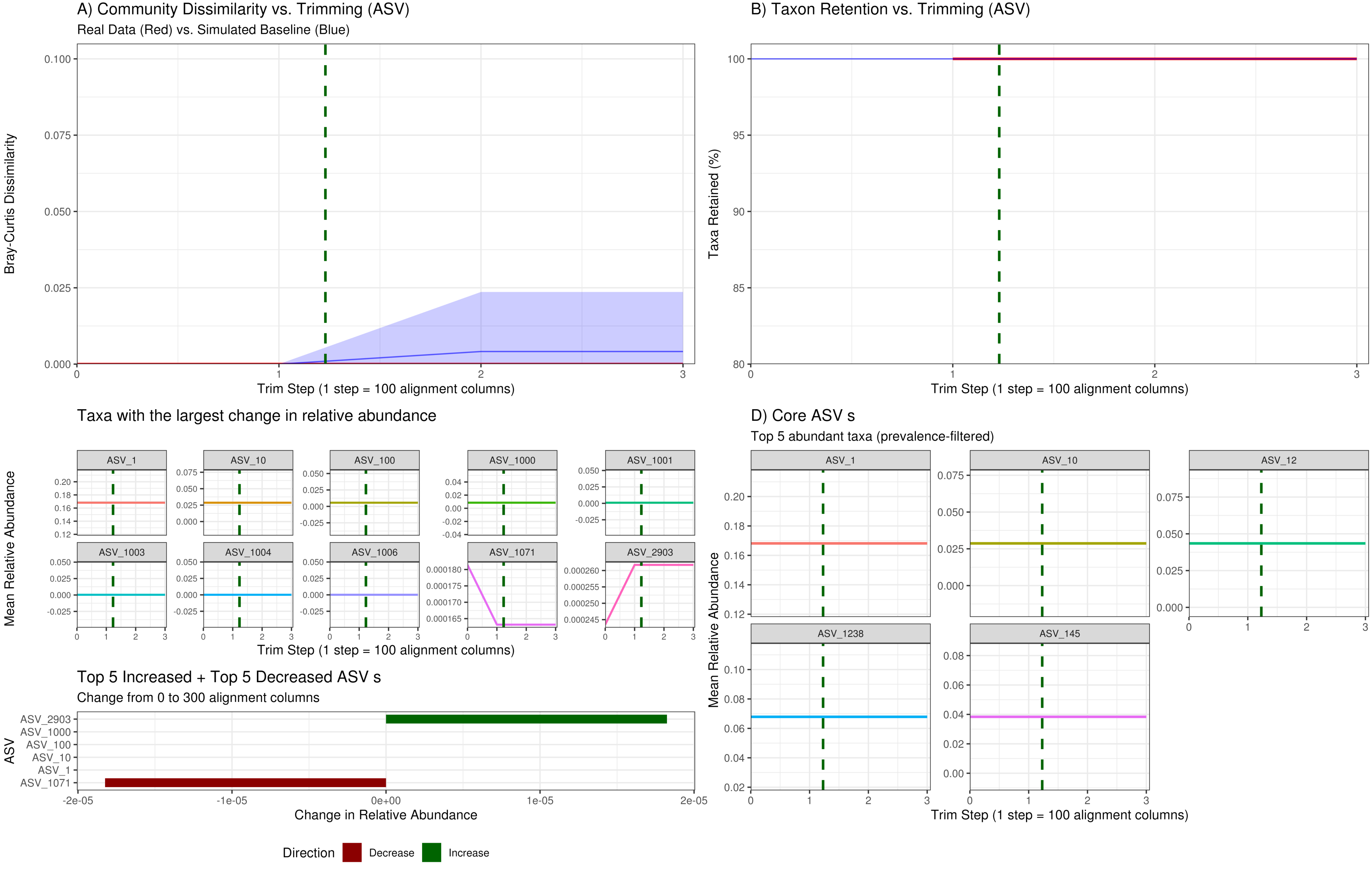


Generated: 2026-09-17 | Primer: 515F_806R | Valid data to 22 alignment columns

### SeCAT Analysis Report: PRJNA734169_SAO

Level: ASV | Method: Warning ... Changepoint only (1/1 methods) | Observed: 1400 bp (1-method warning) | Verdict: CAUTION

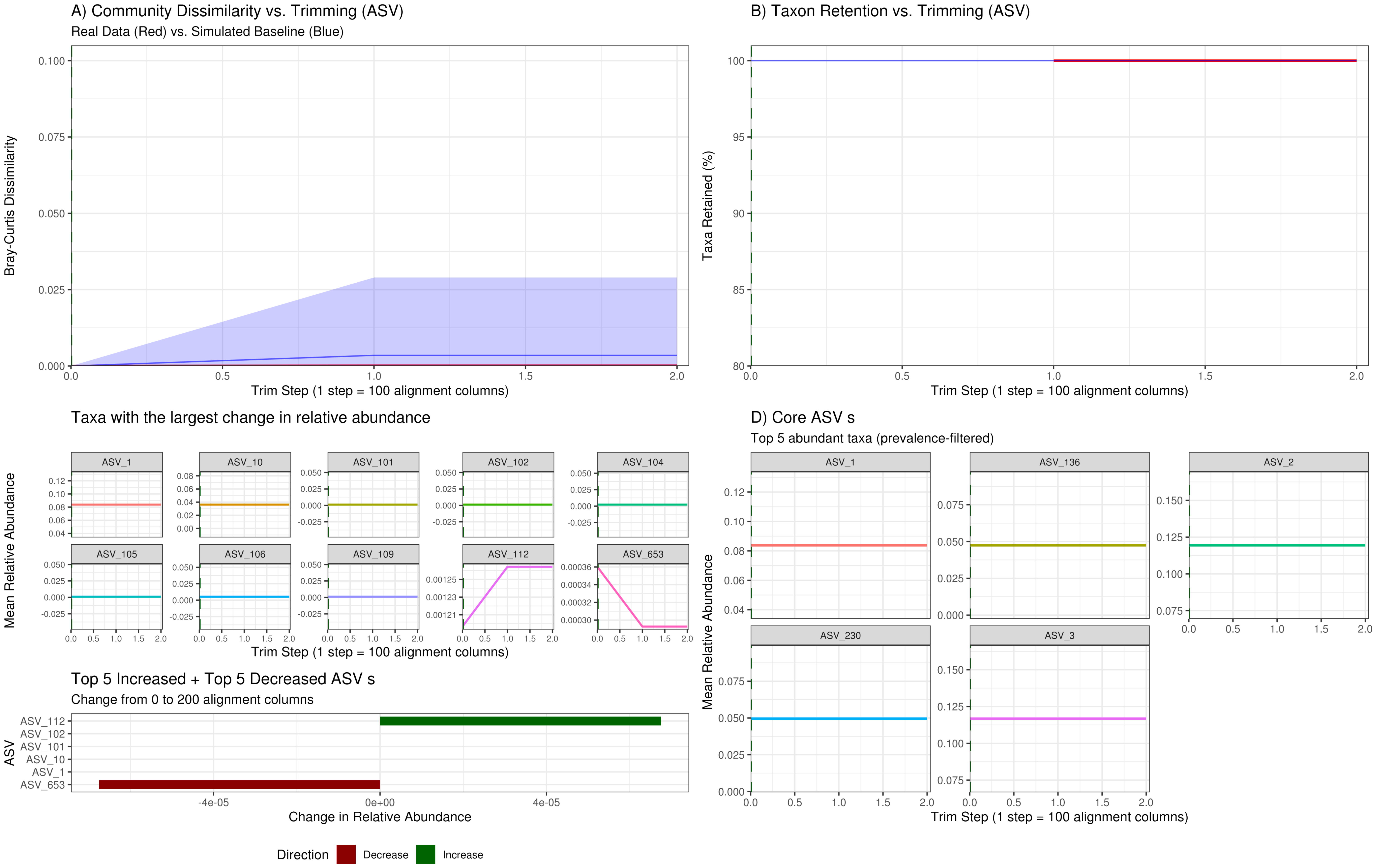


Generated: 2026-09-17 | Primer: 515F_806R | Valid data to 20 alignment columns

### SeCAT Analysis Report: PRJNA903199_PE

Level: ASV | Method: Warning ... Changepoint only (1/1 methods) | Observed: 2000 bp (1-method warning) | Verdict: CAUTION

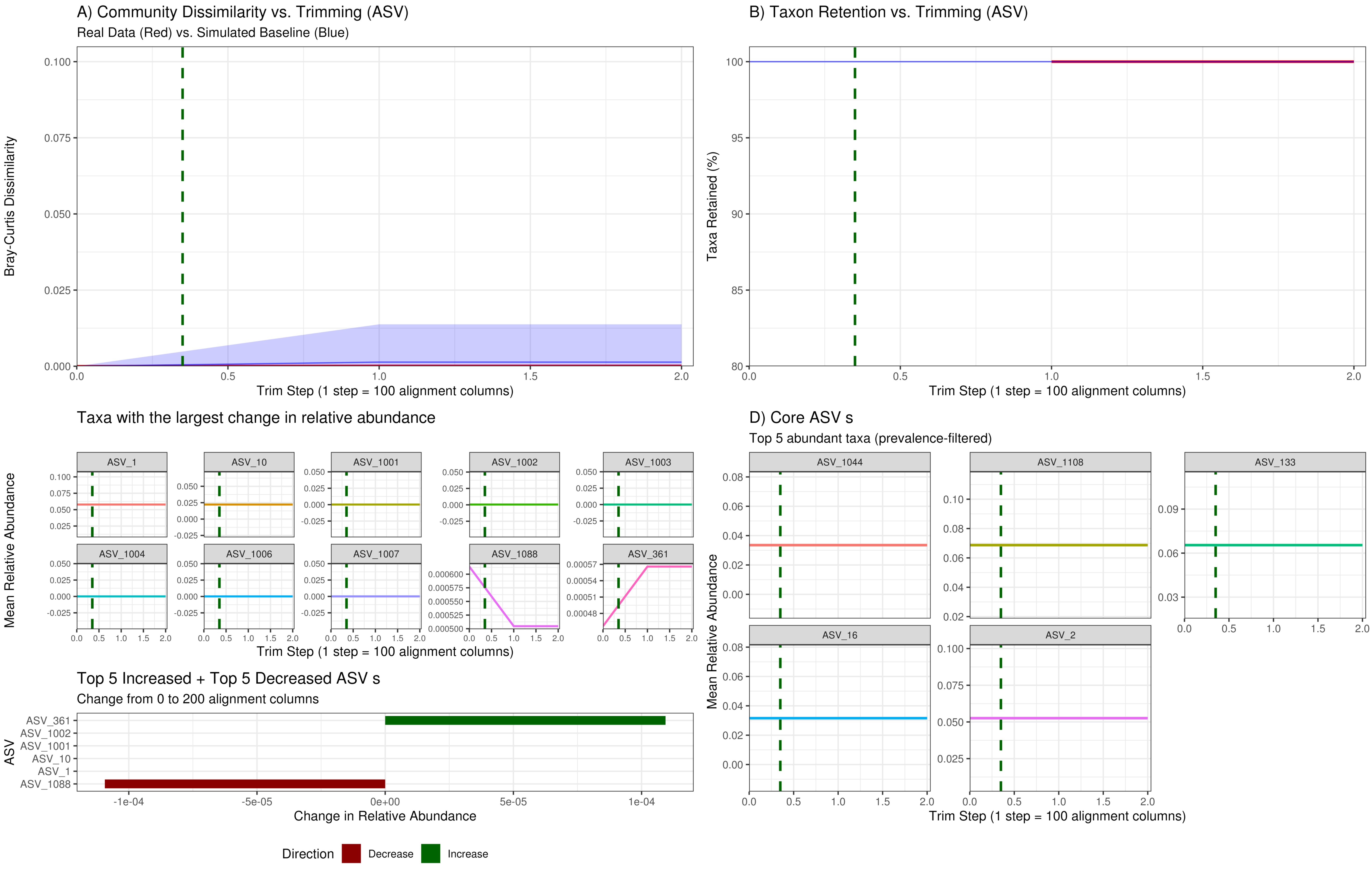
